## Supplementary Material for "Hidden structure in polygenic scores and the challenge of disentangling ancestry interactions in admixed populations"

This PDF file also includes:

– Penn Medicine Biobank Banner Author List and Contributions (p. 35)

– Supplementary Figures S1 to S20

– Supplementary Tables S1 to S5

### Supplementary Proofs and Discussions

#### S1 Computing Polygenic Scores

In computing polygenic scores, we demean haplotypes by ancestry-specific allele frequencies. This is to ensure that polygenic scores in the local model are *invariant* to the binarization scheme at any locus, which is often arbitrary. To see why, we first consider an example of a single dipoid locus that carries two alleles, `C` and `G`. Suppose `C` is the allele assigned 1 (i.e., the ALT allele). We write ancestry-specific allele frequencies at this locus as $f_{\texttt{C}}^{\text{Eur}}$ and $f_{\texttt{C}}^{\text{Afr}}$. On the other hand, if `G` is the ALT allele we write ancestry-specific allele frequencies as $f_{\texttt{G}}^{\text{Eur}}$ and $f_{\texttt{G}}^{\text{Afr}}$. Note that the allele frequencies sum to 1 regardless of ancestry,

$$f_{\texttt{C}}^{\text{Eur}} + f_{\texttt{G}}^{\text{Eur}} = f_{\texttt{C}}^{\text{Afr}} + f_{\texttt{G}}^{\text{Afr}} = 1. \tag{S1}$$

Let us consider how the designation of ALT allele affects the calculation of individual genetic effect, if we simply used allelic dosages. Consider all ten possible genotype-by-local ancestry combinations at this locus, which we have named `1` to `10` for easy reference later. (There are $2^4 = 16$ if the ordering of haplotypes matters, but here the ordering does not matter.) Recall from our Main Text (see Figure 1) that European ancestry is coloured blue and African ancestry is coloured red.

| | 1 | 2 | 3 | 4 | 5 | 6 | 7 | 8 | 9 | 10 |
|---|---|---|---|---|---|---|---|---|---|---|
| Genotype | CG | CC | GC | GG | CG | CC | GG | CG | CC | GG |

First suppose that `C` is designated the ALT allele. The binarized genotype is

| | 1 | 2 | 3 | 4 | 5 | 6 | 7 | 8 | 9 | 10 |
|---|---|---|---|---|---|---|---|---|---|---|
| Genotype | 10 | 11 | 01 | 00 | 10 | 11 | 00 | 10 | 11 | 00 |

Letting $\beta^{\text{Eur}}$ and $\beta^{\text{Afr}}$ be the effect of carrying the `C` allele in European and in African ancestries respectively, the vector of genetic effects across these genotypes is (under the local model, see Eq. (6) of Main Text) given by

| | 1 | 2 | 3 | 4 | 5 | 6 | 7 | 8 | 9 | 10 |
|---|---|---|---|---|---|---|---|---|---|---|
| Genetic Effect | $\beta^{\text{Eur}}$ | $(\beta^{\text{Eur}} + \beta^{\text{Afr}})$ | $\beta^{\text{Afr}}$ | $0$ | $\beta^{\text{Eur}}$ | $2\beta^{\text{Eur}}$ | $0$ | $\beta^{\text{Afr}}$ | $2\beta^{\text{Afr}}$ | $0$ |

Now suppose that `G` is designated the ALT allele. It can be verified that the vector of genetic effects across these genotypes is

| | 1 | 2 | 3 | 4 | 5 | 6 | 7 | 8 | 9 | 10 |
|---|---|---|---|---|---|---|---|---|---|---|
| Genetic Effect | $-\beta^{\text{Afr}}$ | $0$ | $-\beta^{\text{Eur}}$ | $-(\beta^{\text{Eur}} + \beta^{\text{Afr}})$ | $-\beta^{\text{Eur}}$ | $0$ | $-2\beta^{\text{Eur}}$ | $-\beta^{\text{Afr}}$ | $0$ | $-2\beta^{\text{Afr}}$ |

Notice that if the local ancestries are distinct (`1` to `4`), then the difference in genetic effect by binarization is $(\beta^{\text{Eur}} + \beta^{\text{Afr}})$. If the local ancestries are the same on the other hand (`5` to `10`), then the difference in genetic effect by binarization is twice the ancestry-specific effect. This shows that genetic effects at a single locus depend on that locus's binarization scheme, and that the genetic effects across individuals *do not differ by the same shift*. Therefore, if summing across $p$ loci, there are $2^p$ possible binarization schemes that lead to potentially $2^p$ different polygenic score vectors for a sample that are not equivalent up to constant shifts. This raises issues for evaluating polygenic scores, since the non-constant shifts between any pair of score vectors would change the performance (e.g., correlation with phenotype). To make this point clear, for non-admixed cohorts this problem does not occur, because the shifts in genetic effects are the same across all individuals (and so correlation with phenotype would be the same).

We now demonstrate that demeaning allelic dosages leads to polygenic score vectors that are invariant to binarization schemes. First consider the case where C is designated the ALT allele. Demeaning by ancestry-specific allele frequencies $f_{\texttt{C}}^{\text{Eur}}$ and $f_{\texttt{C}}^{\text{Afr}}$, we see for example that the genetic effect for Genotype 1 (CG) is $\beta^{\text{Eur}}(1-f_{\texttt{C}}^{\text{Eur}})+\beta^{\text{Afr}}(-f_{\texttt{C}}^{\text{Afr}})$. We can check that the vector of genetic effects across the ten genotypes is (listed from top to bottom rather than from left to right now)

$$\begin{pmatrix} \beta^{\text{Eur}}(1-f_{\texttt{C}}^{\text{Eur}})+\beta^{\text{Afr}}(-f_{\texttt{C}}^{\text{Afr}}) \\ \beta^{\text{Eur}}(1-f_{\texttt{C}}^{\text{Eur}})+\beta^{\text{Afr}}(1-f_{\texttt{C}}^{\text{Afr}}) \\ \beta^{\text{Eur}}(-f_{\texttt{C}}^{\text{Eur}})+\beta^{\text{Afr}}(1-f_{\texttt{C}}^{\text{Afr}}) \\ \beta^{\text{Eur}}(-f_{\texttt{C}}^{\text{Eur}})+\beta^{\text{Afr}}(-f_{\texttt{C}}^{\text{Afr}}) \\ \beta^{\text{Eur}}(1-f_{\texttt{C}}^{\text{Eur}})+\beta^{\text{Eur}}(-f_{\texttt{C}}^{\text{Eur}}) \\ 2\beta^{\text{Eur}}(1-f_{\texttt{C}}^{\text{Eur}}) \\ 2\beta^{\text{Eur}}(-f_{\texttt{C}}^{\text{Eur}}) \\ \beta^{\text{Afr}}(1-f_{\texttt{C}}^{\text{Afr}})+\beta^{\text{Afr}}(-f_{\texttt{C}}^{\text{Afr}}) \\ 2\beta^{\text{Afr}}(1-f_{\texttt{C}}^{\text{Afr}}) \\ 2\beta^{\text{Afr}}(-f_{\texttt{C}}^{\text{Afr}}) \end{pmatrix}.$$

Next consider the case where G is designated ALT. Remembering to flip the sign of effect and to use the other set of ancestry-specific allele frequencies $f_{\texttt{G}}^{\text{Eur}}$ and $f_{\texttt{G}}^{\text{Afr}}$ for demeaning, we yield

$$\begin{pmatrix} (-\beta^{\text{Eur}})(-f_{\texttt{G}}^{\text{Eur}})+(-\beta^{\text{Afr}})(1-f_{\texttt{G}}^{\text{Afr}}) \\ (-\beta^{\text{Eur}})(-f_{\texttt{G}}^{\text{Eur}})+(-\beta^{\text{Afr}})(-f_{\texttt{G}}^{\text{Afr}}) \\ (-\beta^{\text{Eur}})(1-f_{\texttt{G}}^{\text{Eur}})+(-\beta^{\text{Afr}})(-f_{\texttt{G}}^{\text{Afr}}) \\ (-\beta^{\text{Eur}})(1-f_{\texttt{G}}^{\text{Eur}})+(-\beta^{\text{Afr}})(1-f_{\texttt{G}}^{\text{Afr}}) \\ (-\beta^{\text{Eur}})(-f_{\texttt{G}}^{\text{Eur}})+(-\beta^{\text{Eur}})(1-f_{\texttt{G}}^{\text{Eur}}) \\ 2(-\beta^{\text{Eur}})(-f_{\texttt{G}}^{\text{Eur}}) \\ 2(-\beta^{\text{Eur}})(1-f_{\texttt{G}}^{\text{Eur}}) \\ (-\beta^{\text{Afr}})(-f_{\texttt{G}}^{\text{Afr}})+(-\beta^{\text{Afr}})(1-f_{\texttt{G}}^{\text{Afr}}) \\ 2(-\beta^{\text{Afr}})(-f_{\texttt{G}}^{\text{Afr}}) \\ 2(-\beta^{\text{Afr}})(1-f_{\texttt{G}}^{\text{Afr}}) \end{pmatrix}$$

as the vector of genetic effects. Finally, using Eq. (S1), it is straightforward to check that the two vectors above are equal to each other. Because the vectors are equal, therefore the binarization scheme does not shift genetic effects. So, if we sum genetic effects across $p$ loci after demeaning at each locus, the polygenic effect for an individual does not depend on which of the $2^p$ possible binarization schemes was used.

#### S2 Structural Causal Models Leading to Gene-by-Ancestry Models of Effect Size

There are potentially many causal models that lead to the specific gene-by-ancestry formulation of effect sizes (Eqs. (6) and (7) of Main Text) in our work. Here, we present two that encode G×G interactions, which provides rigour to our claim that "G×A captures epistasis." Other works (e.g., [1]) have investigated the possibility of G×A capturing G×E interactions.

##### Global Model

We introduce a causal model of the marginal effect of a variant in which direct and indirect epistatic effects together contribute to the variant's effect size. This model not only sharpens our intuition about why global ancestry captures *trans non-specific epistasis* [2] across the genome (in particular justifying Eq. (7)), but also motivates future work exploring more data-driven models of *trans* modification of effects.

Consider a focal variant at a gene $\mathtt{f}$, whose marginal effect on a trait measures the contribution to the trait per unit increase in allelic dosage at the focal variant. Assume that the variant's marginal effect is decomposed into its direct effect $b_{\mathtt{f}}$ and the sum of epistatic (or interaction) effects with other peripheral genes — say, $m$ of them — distributed across the genome; see Figure 1C. Note that $b_{\mathtt{f}} = 0$ corresponds to a scenario where the variant effect is entirely composed of its interactions with other variants. This is consistent with the "omnigenic model," which in particular asserts that a peripheral gene's effect on a trait is filtered through multiple networks of genes [3, Figure 1B]. We assume a multiplicative epistasis model governing the interaction effects: for a peripheral gene $\mathtt{g}$ with associated bi-allelic locus, its contribution $b_{\mathtt{g}}$ can be viewed as the effect of carrying one additional copy of the derived allele, conditioned on carrying exactly one copy of the derived allele at the focal variant. We summarize the effect assigned under this model for all other combinations of allelic dosages (for the derived allele) in the table below.

| | | **Focal Variant** | | |
|---|---|---|---|---|
| | **Allelic Dosage** | **0** | **1** | **2** |
| **Peripheral Variant** | **0** | 0 | 0 | 0 |
| | **1** | 0 | $b_{\mathtt{g}}$ | $2b_{\mathtt{g}}$ |
| | **2** | 0 | $2b_{\mathtt{g}}$ | $4b_{\mathtt{g}}$ |

Given these assumptions, what is the marginal effect $\beta$ of the focal variant in an ancestrally homogeneous population? To answer this question, let us assume that allele frequencies for each peripheral gene are known for European and African ancestries: $f_{\mathtt{g}}^{\text{Afr}}$ and $f_{\mathtt{g}}^{\text{Eur}}$, where $\mathtt{g} \in \{1, \dots, m\}$. For a particular individual of fully African ancestry, with allelic dosage vector $\boldsymbol{x} = (x_1, \dots, x_m)$ across the $m$ peripheral genes, their marginal effect is

$$b = b_{\mathtt{f}} + \sum_{\mathtt{g}=1}^{m} b_{\mathtt{g}} x_{\mathtt{g}}. \tag{S2}$$

Taking expectations over individuals in Eq. (S2), the average marginal effect across all individuals of full African ancestry is

$$\beta^{\text{Afr}} := \mathbb{E}[b] = b_{\mathtt{f}} + \sum_{\mathtt{g}=1}^{m} b_{\mathtt{g}} \mathbb{E}[x_{\mathtt{g}}] = b_{\mathtt{f}} + \sum_{\mathtt{g}=1}^{m} 2 b_{\mathtt{g}} f_{\mathtt{g}}^{\text{Afr}}. \tag{S3}$$

The same reasoning shows that the average marginal effect across all individuals of full European ancestry is

$$\beta^{\text{Eur}} = b_{\mathtt{f}} + \sum_{\mathtt{g}=1}^{m} 2 b_{\mathtt{g}} f_{\mathtt{g}}^{\text{Eur}}. \tag{S4}$$

Let us now consider the marginal effect for an admixed individual $i$, whose local ancestry vector and haplotype vector are $\boldsymbol{a} = \left(a_1^{(1)}, \ldots, a_m^{(1)}, a_1^{(2)}, \ldots, a_m^{(2)}\right)$ and $\boldsymbol{x} = \left(x_1^{(1)}, \ldots, x_m^{(1)}, x_1^{(2)}, \ldots, x_m^{(2)}\right)$. Following the notation in the Main Text, let the global ancestry across all peripheral markers be $\overline{a_{i\cdot}} = \frac{1}{2m}\sum_{\mathtt{g}=1}^{m}\left(a_{\mathtt{g}}^{(1)} + a_{\mathtt{g}}^{(2)}\right)$. Their marginal effect is

$$b_i = b_{\mathtt{f}} + \sum_{\mathtt{g}=1}^{m} b_{\mathtt{g}}\left(x_{\mathtt{g}}^{(1)} + x_{\mathtt{g}}^{(2)}\right),$$

which does not involve any local ancestry or global ancestry terms — as it shouldn't. What about the expected effect for the same admixed individual, $\beta_i^{\text{Glo}}$, in case we have neither information about their allelic dosages nor distribution of their local ancestries? We can compute this using the law of iterated expectations.

$$\begin{aligned}
\beta_i^{\text{Glo}} := \mathbb{E}\left[\mathbb{E}\left[b_i|\boldsymbol{a}\right]\right] &= \mathbb{E}\left[b_{\mathtt{f}} + \sum_{\mathtt{g}=1}^{m}\left(\left(a_{\mathtt{g}}^{(1)} + a_{\mathtt{g}}^{(2)}\right) f_{\mathtt{g}}^{\text{Afr}} + \left(2 - a_{\mathtt{g}}^{(1)} - a_{\mathtt{g}}^{(2)}\right) f_{\mathtt{g}}^{\text{Eur}}\right) b_{\mathtt{g}}\right] \\
&= b_{\mathtt{f}} + \sum_{\mathtt{g}=1}^{m}\left(2\overline{a_{i\cdot}} f_{\mathtt{g}}^{\text{Afr}} + 2(1-\overline{a_{i\cdot}}) f_{\mathtt{g}}^{\text{Eur}}\right) b_{\mathtt{g}} \\
&= \overline{a_{i\cdot}}\beta^{\text{Afr}} + (1-\overline{a_{i\cdot}})\beta^{\text{Eur}},
\end{aligned}$$

where in the first equality we used the fact that the average effect at marker $\mathtt{g}$, given local ancestry information, is $\mathbb{E}[b_{\mathtt{g}}|a_{\mathtt{g}}^{(1)}, a_{\mathtt{g}}^{(2)}] = \left[\left(a_{\mathtt{g}}^{(1)} + a_{\mathtt{g}}^{(2)}\right) f_{\mathtt{g}}^{\text{Afr}} + \left(2 - a_{\mathtt{g}}^{(1)} - a_{\mathtt{g}}^{(2)}\right) f_{\mathtt{g}}^{\text{Eur}}\right] b_{\mathtt{g}}$; and the last equality follows from the population-specific marginal effects previously worked out and reported in Eqs. (S3) and (S4). Thus, for an admixed individual $i$, conditioned on knowing just ancestry-specific allele frequencies and their global ancestry across all peripheral markers, the marginal effect of the focal variant for them is $\beta_i^{\text{Glo}} = \overline{a_{i\cdot}}\beta^{\text{Afr}} + (1-\overline{a_{i\cdot}})\beta^{\text{Eur}}$, which corresponds in form to the global model effect size we assumed in Eq. (7). One remaining caveat is that this argument uses just the $m$ peripheral variants that contribute epistatic effects, whereas in practice we compute global ancestry once using all $p$ markers in the genotype array. We perform the latter, because in practice it is difficult to determine, for each focal variant, its set of all other variants that are likely to contribute epistatic effects. It is also possible to use a leave-one-chromosome-out approach to estimate the average local ancestry contributed by the peripheral markers (i.e., excluding the chromosome of the focal marker). However, we find that global ancestry estimation is robust to leaving out chromosome out in practice (Supplementary Figure S1), and so for convenience we relied on the genome-wide global ancestry estimate.

We close this section by noting that the derivations above implicitly assume that the focal marker is independent of the other markers. They would not be correct if the peripheral genes were close to the focal gene (i.e., within the same LD block). To see why the derivation breaks down when this last assumption does not hold, consider the scenario where all of the genes lie within the same LD block, and suppose that having information about the allelic dosages at the peripheral markers allows us to infer the allelic dosage at the focal marker. A special case of this is if we have just one peripheral marker that is completely linked to the focal marker. The conditioning argument would then require us to multiply the direct effect with ancestral allele frequencies (i.e., in Eqs. (S3) and (S4) we have $b_{\mathtt{f}} f_{\mathtt{f}}^{\mathrm{Afr}}$ and $b_{\mathtt{f}} f_{\mathtt{f}}^{\mathrm{Eur}}$ in place of $b_{\mathtt{f}}$). We would also have to multiply each epistatic effect by the relevant ancestral allele frequency of the focal variant. A more mathematical way to think about this is, if we treat allele frequencies as random variables, then the $\sigma$-algebra spanned by the peripheral marker allele frequencies contains the focal marker allele frequency: $f_{\mathtt{f}}^{\mathrm{Eur}} \in \sigma\left(f_1^{\mathrm{Eur}}, \ldots, f_m^{\mathrm{Eur}}\right)$ and $f_{\mathtt{f}}^{\mathrm{Afr}} \in \sigma\left(f_1^{\mathrm{Afr}}, \ldots, f_m^{\mathrm{Afr}}\right)$. So we would have to include the focal variant allelic dosages and its ancestry-specific allele frequencies when computing the expectations. These modifications to the derivation would imply that our conclusion of $\beta_i^{\mathrm{Glo}} = \overline{a_{i\cdot}}\beta^{\mathrm{Afr}} + (1-\overline{a_{i\cdot}})\beta^{\mathrm{Eur}}$ no longer holds.

**Local Model**

The arguments presented above for the global model can be modified to show how the local model arises from *cis* epistasis. Below, we introduce a causal model that incorporates *cis* epistasis; note that similar ideas were explored in Aschard *et al.* [4].

Specifically, we now assume that there are $m$ loci tightly linked with the focal locus carrying the focal SNP. Because of tight linkage, the allelic dosage at each linked locus is roughly the same as the allelic dosage at the focal locus. Let us assume, as above, that a multiplicative epistasis model governs the interaction effects between the $m$ loci and the focal locus.

Given these assumptions, the marginal effect $\beta$ of the focal variant in an ancestrally homogeneous population can be worked out in a similar manner to the global model case. First, similar to Eq. (S2), an individual of fully African ancestry, with allelic dosage vector $\boldsymbol{x} = (x_1, \ldots, x_m)$ across the $m$ other loci, has marginal effect

$$b = b_{\mathtt{f}} + \sum_{\mathtt{g}=1}^{m} b_{\mathtt{g}} x_{\mathtt{g}} \approx b_{\mathtt{f}} + \left(\sum_{\mathtt{g}=1}^{m} b_{\mathtt{g}}\right) x_{\mathtt{f}},$$

where the approximation is owing to tight linkage. Taking expectations over individuals, the average marginal effect across all individuals of full African ancestry is

$$\beta^{\mathrm{Afr}} := \mathbb{E}[b] \approx b_{\mathtt{f}} + \left(\sum_{\mathtt{g}=1}^{m} b_{\mathtt{g}}\right) \mathbb{E}[x_{\mathtt{f}}] = b_{\mathtt{f}} + 2\left(\sum_{\mathtt{g}=1}^{m} b_{\mathtt{g}}\right) f_{\mathtt{f}}^{\mathrm{Afr}}. \tag{S5}$$

The same reasoning shows that the average marginal effect across all individuals of full European ancestry is

$$\beta^{\mathrm{Eur}} \approx b_{\mathtt{f}} + 2\left(\sum_{\mathtt{g}=1}^{m} b_{\mathtt{g}}\right) f_{\mathtt{f}}^{\mathrm{Eur}}. \tag{S6}$$

For an admixed individual $i$, suppose first that their local ancestry vector and haplotype vector are available as $\boldsymbol{a} = \left(a_1^{(1)}, \ldots, a_m^{(1)}, a_1^{(2)}, \ldots, a_m^{(2)}\right)$ and $\boldsymbol{x} = \left(x_1^{(1)}, \ldots, x_m^{(1)}, x_1^{(2)}, \ldots, x_m^{(2)}\right)$. Then their marginal effect is

$$b_i \approx b_{\mathtt{f}} + \sum_{\mathtt{g}=1}^{m} b_{\mathtt{g}} \left(x_{\mathtt{f}}^{(1)} + x_{\mathtt{f}}^{(2)}\right).$$

What about the expected effect for the same admixed individual, $\beta_i^{\mathrm{Loc}}$, in case we have neither information about their allelic dosages nor distribution of their local ancestries? We can compute this using the law of iterated expectations.

$$\begin{aligned}
\beta_i^{\mathrm{Loc}} := \mathbb{E}\left[\mathbb{E}\left[b_i|\boldsymbol{a}\right]\right] &\approx \mathbb{E}\left[b_{\mathtt{f}} + \sum_{\mathtt{g}=1}^{m}\left(\left(a_{\mathtt{f}}^{(1)} + a_{\mathtt{f}}^{(2)}\right) f_{\mathtt{f}}^{\mathrm{Afr}} + \left(2 - a_{\mathtt{f}}^{(1)} - a_{\mathtt{f}}^{(2)}\right) f_{\mathtt{f}}^{\mathrm{Eur}}\right) b_{\mathtt{g}}\right] \\
&= \left(a_{\mathtt{f}}^{(1)} + a_{\mathtt{f}}^{(2)}\right)\left[\frac{1}{2}b_{\mathtt{f}} + \left(\sum_{\mathtt{g}=1}^{m} b_{\mathtt{g}}\right) f_{\mathtt{f}}^{\mathrm{Afr}}\right] + \left(2 - a_{\mathtt{f}}^{(1)} - a_{\mathtt{f}}^{(2)}\right)\left[\frac{1}{2}b_{\mathtt{f}} + \left(\sum_{\mathtt{g}=1}^{m} b_{\mathtt{g}}\right) f_{\mathtt{f}}^{\mathrm{Eur}}\right] \\
&= \left(a_{\mathtt{f}}^{(1)} + a_{\mathtt{f}}^{(2)}\right)\frac{\beta^{\mathrm{Afr}}}{2} + \left(2 - a_{\mathtt{f}}^{(1)} - a_{\mathtt{f}}^{(2)}\right)\frac{\beta^{\mathrm{Eur}}}{2},
\end{aligned}$$

where we use the same reasoning as in the global model derivation above, and in particular the last equality following from the population-specific marginal effects previously worked out and reported in Eqs. (S5) and (S6). Thus, for an admixed individual $i$, conditioned on knowing just ancestry-specific allele frequencies and their local ancestries across all peripheral markers, the marginal effect of the focal variant for them is the sum of marginal effects on their two haplotypes: $\beta_i^{\mathrm{Loc}} = \frac{1}{2}\sum_{h=1,2}\left[a_{\mathtt{f}}^{(h)}\beta^{\mathrm{Afr}} + \left(1 - a_{\mathtt{f}}^{(h)}\right)\beta^{\mathrm{Eur}}\right]$. Up to a scaling factor, this corresponds in form to the local model effect size we assumed in Eq. (6).

#### S3 Relationship Between Causal and Tagging Effect Sizes

We show how tagging effect size depends on causal effect size, allele frequency and LD for quantitative traits, per Eqs. (9) and (10). We mirror the proofs presented in Zaidi [5] and Vukcevic *et al.* [6] for case-control traits, which formulate the causal effect as an average treatment effect in an ancestrally homogeneous population.

Denote the causal and tagging variant allele dosages by $X'$ and $X$, and the phenotype by $Y$. First, the structural causal model we have in mind is the following: the causal variant $X'$ has a causal effect on the trait $Y$, but the tagging variant $X$ is correlated with it. (Imagine a graph with three nodes, $Y, X'$ and $X$. The node for $X'$ points to the node for $Y$, while the nodes $X$ and $X'$ are connected by a non-directional segment to indicate linkage. To be pedantic, there should be a fourth node $A$, denoting ancestry, parent to both $X$ and $X'$, but we are conditioning on it, which induces correlation between the children $X'$ and $X$.) Let the average effects of carrying zero and one copy of the causal allele be $\mu$ and $\mu + \beta'$ (both $\mu$ and $\beta'$ are real-valued):

$$\begin{aligned} \mathbb{E}\left[Y|X'=0\right] &= \mu \\ \mathbb{E}\left[Y|X'=1\right] &= \mu+\beta' \end{aligned}$$

From the equations above, it is clear that the causal effect is just the average treatment effect: $\beta' = \mathbb{E}[Y|X'=1] - \mathbb{E}[Y|X'=0]$. We assume that the causal effect is additive, so that the effect of carrying two copies of the causal allele (i.e., $X'=2$) is just $2\beta'$. This means we do not have to worry about diploidy for the remainder of the argument. Now, let the average effects of carrying zero and one copy of the tagging allele be $\gamma_0 = \mathbb{E}[Y|X=0]$ and $\gamma_1 = \mathbb{E}[Y|X=1]$. We are interested in the quantity $\beta = \gamma_1 - \gamma_0$, which describes the tagging effect.

To compute $\beta$, we require causal and tagging allele frequencies:

$$\begin{aligned} \mathbb{P}(X'=1) &= f' \\ \mathbb{P}(X'=0) &= 1-f' \\ \mathbb{P}(X=1) &= f \\ \mathbb{P}(X=0) &= 1-f \end{aligned}$$

Let us introduce shorthand notation for conditional probabilities of observing particular dosages of tagging variant alleles:

$$\begin{aligned} \mathbb{P}(X=0|X'=0) &= q_{00} \\ \mathbb{P}(X=1|X'=0) &= 1-q_{00} \\ \mathbb{P}(X=0|X'=1) &= q_{10} \\ \mathbb{P}(X=1|X'=1) &= 1-q_{10} \end{aligned}$$

With the notation above, we can express linkage equilibrium (LD) compactly. The LD between the causal and tagging variants is $\lambda = [\mathbb{P}(X=1, X'=1) - ff']/\sqrt{f(1-f)f'(1-f')}$. Since $f = \mathbb{P}(X=1) = \mathbb{P}(X=1, X'=1) + \mathbb{P}(X=1, X'=0) = (1-q_{10})f' + (1-q_{00})(1-f')$, we may substitute the long expression for $f$ in the numerator of LD, which gives

$$\begin{aligned}
\lambda &= \frac{\mathbb{P}(X=1,X'=1) - f'[(1-q_{10})f' + (1-q_{00})(1-f')]}{\sqrt{f(1-f)f'(1-f')}} \\
&= \frac{(1-q_{10})f' - f'[(1-q_{10})f' + (1-q_{00})(1-f')]}{\sqrt{f(1-f)f'(1-f')}} \\
&= \frac{[(1-q_{10}) - (1-q_{00})](1-f')f'}{\sqrt{f(1-f)f'(1-f')}} \\
&= (q_{00} - q_{10})\sqrt{\frac{f'(1-f')}{f(1-f)}}.
\end{aligned}$$

Finally, denote the joint densities of phenotype and the allele dosage variables by $p_{Y,X'}$ and $p_{Y,X}$, and similarly denote all conditional densities using the "$|$" symbol so that there are no ambiguities (e.g., the conditional density of phenotype $Y$ given that $X'=0$ is $p_{Y|X'}(y|0)$).

Back to computing $\beta$, it suffices to compute $\gamma_0$ and $\gamma_1$. Observe that

$$\begin{aligned}
\gamma_0 &= \frac{\int_{\mathbb{R}} y \cdot p_{Y,X}(y,0)dy}{\mathbb{P}(X=0)} \\
&= \frac{1}{(1-f)}\int_{\mathbb{R}} y \cdot \left[p_{Y,X|X'}(y,0|0)\mathbb{P}(X'=0) + p_{Y,X|X'}(y,0|1)\mathbb{P}(X'=1)\right] dy \\
&= \frac{1}{(1-f)}\int_{\mathbb{R}} y \cdot \left[p_{Y|X'}(y|0)\mathbb{P}(X=0|X'=0)(1-f') + p_{Y|X'}(y|1)\mathbb{P}(X=0|X'=1)f'\right] dy \quad (*) \\
&= \frac{1}{(1-f)}\left[q_{00}(1-f')\int_{\mathbb{R}} y \cdot p_{Y|X'}(y|0)dy + q_{10}f'\int_{\mathbb{R}} p_{Y|X'}(y|1)dy\right] \\
&= \frac{q_{00}(1-f')\mu + q_{10}f'(\mu+\beta')}{(1-f)},
\end{aligned}$$

where in $(*)$ we used the fact that $X$ and $Y$ are independent conditioned on $X'$. By reasoning similarly, we obtain

$$\gamma_1 = \frac{(1-q_{00})(1-f')\mu + (1-q_{10})f'(\mu+\beta')}{f}.$$

Taking the difference of these two quantities (which is $\beta$), we obtain

$$\begin{aligned}
\gamma_1 - \gamma_0 &= \left[\frac{(1-q_{00})(1-f') + (1-q_{10})f'}{f} - \frac{q_{00}(1-f') + q_{10}f'}{(1-f)}\right]\mu + \left[\frac{(1-q_{10})f'}{f} - \frac{q_{10}f'}{(1-f)}\right]\beta' \\
&= \left(\frac{(1-f) - [f'q_{10} + (1-f')q_{00}]}{f(1-f)}\right)\mu + \left(\frac{f'(1-q_{10}-f)}{f(1-f)}\right)\beta'.
\end{aligned}$$

We now show that (1) the coefficient of $\mu$ is zero; and (2) the coefficient of $\beta'$ reduces to the quantity analogous to the expressions in Eqs. (9) and (10). First, the numerator of the coefficient of $\mu$, $(1-f) - [f'q_{10} + (1-f')q_{00}]$ is zero, because both $1-f$ and $f'q_{10} + (1-f')q_{00}$ are equal to $\mathbb{P}(X=0)$. This proves (1). Next, using the same substitution trick for $f$ that we used to obtain the expression for LD earlier, we have

$$\begin{aligned}
\frac{f'(1-q_{10}-f)}{f(1-f)} &= \frac{f'[1-q_{10}-(1-q_{10})f'-(1-q_{00})(1-f')]}{f(1-f)} \\
&= \frac{f'(1-f')[(1-q_{10})-(1-q_{00})]}{f(1-f)} \\
&= (q_{00}-q_{10})\frac{f'(1-f')}{f(1-f)} \\
&= \lambda\sqrt{\frac{f'(1-f')}{f(1-f)}},
\end{aligned}$$

which proves (2). Summarizing the above, we obtain

$$\gamma_1 - \gamma_0 = \beta' \ \lambda\sqrt{\frac{f'(1-f')}{f(1-f)}},$$

which is the desired expression.

#### S4 Analytical Properties of Causal Variant Polygenic Scores

We present analytical expressions for the squared correlation between total or standard polygenic scores (TotPGS) and phenotype, as well as between partial polygenic scores (ParPGS) and phenotype, assuming that causal variants and their effects are known. The calculations here will be useful for deriving analytical expressions in the case of tagging variants (Supplementary Material Subsection S6).

We begin by stating that our models treat the following quantities as fixed: phased genotype matrix $\mathbf{X}$ and inferred haplotype local ancestry matrix $\mathbf{A}$. Ancestry-specific causal and tagging allele frequencies, $\hat{f}_j'^{\text{Eur}}, \hat{f}_j'^{\text{Afr}}$ and $\hat{f}_j^{\text{Eur}}, \hat{f}_j^{\text{Afr}}$ as defined in the Main Text (Subsection 3.1), are estimated from phased genotypes and local ancestries. The random quantities are the causal genetic effects $\beta_j'^{\text{Eur}}$ and $\beta_j'^{\text{Afr}}$ as defined by Eq. (2) in the Main Text, as well as the noise term $\varepsilon_i$ that will be irrelevant to our technical derivations. To arrive at the analytical expressions, we first work out empirical (i.e., sample) quantities assuming the genetic effects and exogenous noise are assigned (Step 1), and then work out the expectation of these quantities under the distributional assumptions of genetic effects and noise (Step 2).

Let us begin by recalling the definition of some empirical quantities. The sample correlation between two sample vectors $\boldsymbol{u} = (u_1, \ldots, u_n)$ and $\boldsymbol{v} = (v_1, \ldots, v_n)$ is given by

$$\text{Cor}(\boldsymbol{u}, \boldsymbol{v}) = \frac{\text{Cov}(\boldsymbol{u}, \boldsymbol{v})}{\sqrt{\text{Var}(\boldsymbol{u})\text{Var}(\boldsymbol{v})}}, \tag{S7}$$

where $\text{Cov}(\boldsymbol{u}, \boldsymbol{v})$ and $\text{Var}(\boldsymbol{u})$ are the sample covariance and sample variance, which are defined by

$$\text{Cov}(\boldsymbol{u}, \boldsymbol{v}) = \frac{1}{n-1}\sum_{i=1}^{n}(u_i - \bar{u})(v_i - \bar{v}) \tag{S8}$$

and

$$\text{Var}(\boldsymbol{u}) = \frac{1}{n-1}\sum_{i=1}^{n}(u_i - \bar{u})^2. \tag{S9}$$

($\text{Var}(\boldsymbol{v})$ is defined in a similar manner to Eq. (S9).) Here, $\bar{u} = (u_1 + \ldots + u_n)/n$ and $\bar{v} = (v_1 + \ldots + v_n)/n$ are sample means. Throughout this document, we will always use Var, Cov and Cor to denote sample quantities; properties of random variables and vectors, like mean and variance, will be denoted $\mathbb{E}$ and $\mathbb{V}$. We use $i$ to index individuals unless specified otherwise. To reduce symbolic clutter, we will also collect individual quantities into vectors by boldfacing the quantity, for example writing the vector $(x_{ij} : i = 1, \ldots, n)$ as $\boldsymbol{x_{\cdot j}}$.

Step 1: Empirical quantities under assigned genetic effects and exogenous noise.

**Variance of TotPGS.** Polygenic scores are computed with causal variants and their effect sizes in this scenario. Applying algebraic rules of the sample variance to Eq. (12) of the Main Text yields

$$\begin{aligned}
\mathrm{Var}(\mathbf{TotPGS}) &= \mathrm{Var}\left(\sum_{j=1}^{p} \beta_j'^{\mathrm{Eur}} \left(\hat{\boldsymbol{x}}_{\boldsymbol{\cdot j}}'^{\boldsymbol{(1)}} + \hat{\boldsymbol{x}}_{\boldsymbol{\cdot j}}'^{\boldsymbol{(2)}}\right)\right) \\
&= \sum_{j=1}^{p} \left(\beta_j'^{\mathrm{Eur}}\right)^2 \mathrm{Var}\left(\hat{\boldsymbol{x}}_{\boldsymbol{\cdot j}}'^{\boldsymbol{(1)}} + \hat{\boldsymbol{x}}_{\boldsymbol{\cdot j}}'^{\boldsymbol{(2)}}\right) \qquad \text{(independence of markers)} \\
&= \sum_{j=1}^{p} \Bigg[ \underbrace{\left(\beta_j'^{\mathrm{Eur}}\right)^2 \mathrm{Var}\left(\hat{\boldsymbol{x}}_{\boldsymbol{\cdot j}}'^{\boldsymbol{(1)}}\right)}_{\text{1'}} + \underbrace{\left(\beta_j'^{\mathrm{Eur}}\right)^2 \mathrm{Var}\left(\hat{\boldsymbol{x}}_{\boldsymbol{\cdot j}}'^{\boldsymbol{(2)}}\right)}_{\text{2'}} + \underbrace{2 \times \left(\beta_j'^{\mathrm{Eur}}\right)^2 \mathrm{Cov}\left(\hat{\boldsymbol{x}}_{\boldsymbol{\cdot j}}'^{\boldsymbol{(1)}}, \hat{\boldsymbol{x}}_{\boldsymbol{\cdot j}}'^{\boldsymbol{(2)}}\right)}_{\text{3'}} \Bigg].
\end{aligned}$$

This shows that each marker contributing to the variance of the total polygenic score can be decomposed into three terms: two haplotype-specific genetic variances ((1') and (2')) followed by a cross-haplotype covariance ((3')). Let us simplify each term into expressions involving only fixed and random quantities.

$$\begin{aligned}
\text{(1')} &= \left(\beta_j'^{\mathrm{Eur}}\right)^2 \frac{1}{n-1} \sum_{i=1}^{n} \left(\hat{x}_{ij}'^{(1)} - \frac{1}{n}\sum_{i=1}^{n} \hat{x}_{ij}'^{(1)}\right)^2 \qquad \text{(by Eq. (S9))} \\
&= \left(\beta_j'^{\mathrm{Eur}}\right)^2 \frac{1}{n-1} \sum_{i=1}^{n} \left(\hat{x}_{ij}'^{(1)}\right)^2 \qquad ([\hat{x}_{ij}'^{(1)} : a_{ij}'^{(1)} = 0] \text{ and } [\hat{x}_{ij}'^{(1)} : a_{ij}'^{(1)} = 1] \text{ are demeaned}) \\
&= \left(\beta_j'^{\mathrm{Eur}}\right)^2 \frac{1}{n-1} \left[ \sum_{i: a_{ij}'^{(1)}=0} \left(\hat{x}_{ij}'^{(1)}\right)^2 + \sum_{i: a_{ij}'^{(1)}=1} \left(\hat{x}_{ij}'^{(1)}\right)^2 \right] \\
&= \left(\beta_j'^{\mathrm{Eur}}\right)^2 \frac{n}{n-1} \left[ \left(1 - \overline{a_{\cdot j}'^{(1)}}\right) \hat{f}_j'^{\mathrm{Eur}} \left(1 - \hat{f}_j'^{\mathrm{Eur}}\right) + \overline{a_{\cdot j}'^{(1)}} \hat{f}_j'^{\mathrm{Afr}} \left(1 - \hat{f}_j'^{\mathrm{Afr}}\right) \right],
\end{aligned}$$

where $\overline{a_{\cdot j}'^{(h)}} = \frac{1}{n}\left(a_{1j}'^{(h)} + \ldots + a_{nj}'^{(h)}\right)$ denotes the African ancestry fraction at the $j$th marker on haplotype $h$ ($h = 1, 2$). Similarly,

$$\text{(2')} = \left(\beta_j'^{\mathrm{Eur}}\right)^2 \frac{n}{n-1} \left[ \left(1 - \overline{a_{\cdot j}'^{(2)}}\right) \hat{f}_j'^{\mathrm{Eur}} \left(1 - \hat{f}_j'^{\mathrm{Eur}}\right) + \overline{a_{\cdot j}'^{(2)}} \hat{f}_j'^{\mathrm{Afr}} \left(1 - \hat{f}_j'^{\mathrm{Afr}}\right) \right].$$

Finally,

$$\begin{aligned}
\text{(3')} &= 2 \times \left(\beta_j'^{\mathrm{Eur}}\right)^2 \frac{1}{n-1} \sum_{i=1}^{n} \left(\hat{x}_{ij}'^{(1)} - \frac{1}{n}\sum_{i=1}^{n} \hat{x}_{ij}'^{(1)}\right)\left(\hat{x}_{ij}'^{(2)} - \frac{1}{n}\sum_{i=1}^{n} \hat{x}_{ij}'^{(2)}\right) \qquad \text{(by Eq. (S8))} \\
&= 2 \times \left(\beta_j'^{\mathrm{Eur}}\right)^2 \frac{1}{n-1} \sum_{i=1}^{n} \hat{x}_{ij}'^{(1)} \hat{x}_{ij}'^{(2)} \qquad \text{(allele dosages are demeaned)}
\end{aligned}$$

**Variance of ParPGS.** Applying properties of the sample variance to Eq. (13) of the Main Text,

$$\begin{aligned}
\text{Var}(\textbf{ParPGS}) &= \text{Var}\left(\sum_{j=1}^{p} \beta_j^{\prime\text{Eur}} \left[\left(\mathbf{1} - \boldsymbol{a}_{\cdot\boldsymbol{j}}^{\boldsymbol{\prime(1)}}\right) \hat{\boldsymbol{x}}_{\cdot\boldsymbol{j}}^{\boldsymbol{\prime(1)}} + \left(\mathbf{1} - \boldsymbol{a}_{\cdot\boldsymbol{j}}^{\boldsymbol{\prime(2)}}\right) \hat{\boldsymbol{x}}_{\cdot\boldsymbol{j}}^{\boldsymbol{\prime(2)}}\right]\right) \\
&= \sum_{j=1}^{p} \left(\beta_j^{\prime\text{Eur}}\right)^2 \text{Var}\left(\left(\mathbf{1} - \boldsymbol{a}_{\cdot\boldsymbol{j}}^{\boldsymbol{\prime(1)}}\right) \hat{\boldsymbol{x}}_{\cdot\boldsymbol{j}}^{\boldsymbol{\prime(1)}} + \left(\mathbf{1} - \boldsymbol{a}_{\cdot\boldsymbol{j}}^{\boldsymbol{\prime(2)}}\right) \hat{\boldsymbol{x}}_{\cdot\boldsymbol{j}}^{\boldsymbol{\prime(2)}}\right) \\
&\qquad\qquad\qquad\qquad\qquad\qquad\qquad\qquad\qquad\qquad\text{(independence of markers)} \\
&= \sum_{j=1}^{p} \Bigg[\underbrace{\left(\beta_j^{\prime\text{Eur}}\right)^2 \text{Var}\left(\left(\mathbf{1} - \boldsymbol{a}_{\cdot\boldsymbol{j}}^{\boldsymbol{\prime(1)}}\right) \hat{\boldsymbol{x}}_{\cdot\boldsymbol{j}}^{\boldsymbol{\prime(1)}}\right)}_{\text{(4')}} + \underbrace{\left(\beta_j^{\prime\text{Eur}}\right)^2 \text{Var}\left(\left(\mathbf{1} - \boldsymbol{a}_{\cdot\boldsymbol{j}}^{\boldsymbol{\prime(2)}}\right) \hat{\boldsymbol{x}}_{\cdot\boldsymbol{j}}^{\boldsymbol{\prime(2)}}\right)}_{\text{(5')}} \\
&\qquad + \underbrace{2 \times \left(\beta_j^{\prime\text{Eur}}\right)^2 \text{Cov}\left(\left(\mathbf{1} - \boldsymbol{a}_{\cdot\boldsymbol{j}}^{\boldsymbol{\prime(1)}}\right) \hat{\boldsymbol{x}}_{\cdot\boldsymbol{j}}^{\boldsymbol{\prime(1)}}, \left(\mathbf{1} - \boldsymbol{a}_{\cdot\boldsymbol{j}}^{\boldsymbol{\prime(2)}}\right) \hat{\boldsymbol{x}}_{\cdot\boldsymbol{j}}^{\boldsymbol{\prime(2)}}\right)}_{\text{(6')}}\Bigg].
\end{aligned}$$

The terms here are similar to those obtained in the Variance of TotPGS above, and they simplify as follows:

$$\begin{aligned}
\text{(4')} &= \left(\beta_j^{\prime\text{Eur}}\right)^2 \frac{n}{n-1} \left(1 - \overline{a_{\cdot j}^{\prime(1)}}\right) \hat{f}_j^{\prime\text{Eur}} \left(1 - \hat{f}_j^{\prime\text{Eur}}\right), \\
\text{(5')} &= \left(\beta_j^{\prime\text{Eur}}\right)^2 \frac{n}{n-1} \left(1 - \overline{a_{\cdot j}^{\prime(2)}}\right) \hat{f}_j^{\prime\text{Eur}} \left(1 - \hat{f}_j^{\prime\text{Eur}}\right), \\
\text{(6')} &= 2 \times \left(\beta_j^{\prime\text{Eur}}\right)^2 \frac{1}{n-1} \sum_{i=1}^{n} \left(1 - a_{ij}^{\prime(1)}\right) \left(1 - a_{ij}^{\prime(2)}\right) \hat{x}_{ij}^{\prime(1)} \hat{x}_{ij}^{\prime(2)}.
\end{aligned}$$

**Variance of Phenotype.** In our simulations, $\text{Var}(\boldsymbol{y})$ is 1 in expectation. We find in practice that $\text{Var}(\boldsymbol{y}) \approx 1$ even for a single draw from our models, so we approximate $\text{Var}(\boldsymbol{y})$ by its expectation.

**Covariance of TotPGS with Phenotype.** These depend on whether the local model or global model is assumed. We work out the expressions under each model.

*Local Model.* By Eq. (6) of the Main Text, the phenotype of individual $i$ is

$$y_i = \sum_{j=1}^{p} \left(\left[\beta_j^{\prime\text{Eur}} \left(1 - a_{ij}^{\prime(1)}\right) + \beta_j^{\prime\text{Afr}} a_{ij}^{\prime(1)}\right] \hat{x}_{ij}^{\prime(1)} + \left[\beta_j^{\prime\text{Eur}} \left(1 - a_{ij}^{\prime(2)}\right) + \beta_j^{\prime\text{Afr}} a_{ij}^{\prime(2)}\right] \hat{x}_{ij}^{\prime(2)}\right) + \varepsilon_i.$$

Because the noise term $\varepsilon_i$ is exogenous, its contribution to the overall covariance is, on average, zero. In practice the covariance is close to 0 even for a single draw, so we drop it from our calculations.

By independence of markers and algebraic rules of the sample covariance, we obtain

$$
\begin{aligned}
\operatorname{Cov}(\mathbf{TotPGS}, \boldsymbol{y}) \;=\; \sum_{j=1}^{p} \Bigg\{ & \underbrace{\operatorname{Cov}\left(\beta_j'^{\text{Eur}} \hat{\boldsymbol{x}}_{\cdot j}'^{(1)}, \beta_j'^{\text{Eur}} \left(\mathbf{1} - \boldsymbol{a}_{\cdot j}'^{(1)}\right) \hat{\boldsymbol{x}}_{\cdot j}'^{(1)} + \beta_j'^{\text{Afr}} \boldsymbol{a}_{\cdot j}'^{(1)} \hat{\boldsymbol{x}}_{\cdot j}'^{(1)}\right)}_{\text{(7L')}} \\
& + \underbrace{\operatorname{Cov}\left(\beta_j'^{\text{Eur}} \hat{\boldsymbol{x}}_{\cdot j}'^{(1)}, \beta_j'^{\text{Eur}} \left(\mathbf{1} - \boldsymbol{a}_{\cdot j}'^{(2)}\right) \hat{\boldsymbol{x}}_{\cdot j}'^{(2)} + \beta_j'^{\text{Afr}} \boldsymbol{a}_{\cdot j}'^{(2)} \hat{\boldsymbol{x}}_{\cdot j}'^{(2)}\right)}_{\text{(8L')}} \\
& + \underbrace{\operatorname{Cov}\left(\beta_j'^{\text{Eur}} \hat{\boldsymbol{x}}_{\cdot j}'^{(2)}, \beta_j'^{\text{Eur}} \left(\mathbf{1} - \boldsymbol{a}_{\cdot j}'^{(1)}\right) \hat{\boldsymbol{x}}_{\cdot j}'^{(1)} + \beta_j'^{\text{Afr}} \boldsymbol{a}_{\cdot j}'^{(1)} \hat{\boldsymbol{x}}_{\cdot j}'^{(1)}\right)}_{\text{(9L')}} \\
& + \underbrace{\operatorname{Cov}\left(\beta_j'^{\text{Eur}} \hat{\boldsymbol{x}}_{\cdot j}'^{(2)}, \beta_j'^{\text{Eur}} \left(\mathbf{1} - \boldsymbol{a}_{\cdot j}'^{(2)}\right) \hat{\boldsymbol{x}}_{\cdot j}'^{(2)} + \beta_j'^{\text{Afr}} \boldsymbol{a}_{\cdot j}'^{(2)} \hat{\boldsymbol{x}}_{\cdot j}'^{(2)}\right)}_{\text{(10L')}} \Bigg\}
\end{aligned}
$$

The four terms simplify as follows:

$$
\begin{aligned}
\text{(7L')} &= \underbrace{\left(\beta_j'^{\text{Eur}}\right)^2 \frac{n}{n-1} \left(1 - \overline{a_{\cdot j}'^{(1)}}\right) \hat{f}_j'^{\text{Eur}} \left(1 - \hat{f}_j'^{\text{Eur}}\right)}_{\text{(7L')}_1} + \underbrace{\beta_j'^{\text{Eur}} \beta_j'^{\text{Afr}} \frac{n}{n-1} \overline{a_{\cdot j}'^{(1)}} \hat{f}_j'^{\text{Afr}} \left(1 - \hat{f}_j'^{\text{Afr}}\right)}_{\text{(7L')}_2}, \\
\text{(8L')} &= \underbrace{\left(\beta_j'^{\text{Eur}}\right)^2 \frac{1}{n-1} \sum_{i=1}^{n} \left(1 - a_{ij}'^{(2)}\right) \hat{x}_{ij}'^{(1)} \hat{x}_{ij}'^{(2)}}_{\text{(8L')}_1} + \underbrace{\beta_j'^{\text{Eur}} \beta_j'^{\text{Afr}} \frac{1}{n-1} \sum_{i=1}^{n} a_{ij}'^{(2)} \hat{x}_{ij}'^{(1)} \hat{x}_{ij}'^{(2)}}_{\text{(8L')}_2}, \\
\text{(9L')} &= \underbrace{\left(\beta_j'^{\text{Eur}}\right)^2 \frac{1}{n-1} \sum_{i=1}^{n} \left(1 - a_{ij}'^{(1)}\right) \hat{x}_{ij}'^{(1)} \hat{x}_{ij}'^{(2)}}_{\text{(9L')}_1} + \underbrace{\beta_j'^{\text{Eur}} \beta_j'^{\text{Afr}} \frac{1}{n-1} \sum_{i=1}^{n} a_{ij}'^{(1)} \hat{x}_{ij}'^{(1)} \hat{x}_{ij}'^{(2)}}_{\text{(9L')}_2}, \\
\text{(10L')} &= \underbrace{\left(\beta_j'^{\text{Eur}}\right)^2 \frac{n}{n-1} \left(1 - \overline{a_{\cdot j}'^{(2)}}\right) \hat{f}_j'^{\text{Eur}} \left(1 - \hat{f}_j'^{\text{Eur}}\right)}_{\text{(10L')}_1} + \underbrace{\beta_j'^{\text{Eur}} \beta_j'^{\text{Afr}} \frac{n}{n-1} \overline{a_{\cdot j}'^{(2)}} \hat{f}_j'^{\text{Afr}} \left(1 - \hat{f}_j'^{\text{Afr}}\right)}_{\text{(10L')}_2}.
\end{aligned}
$$

*Global Model.* By Eq. (7) of the Main Text, the phenotype of individual $i$ is

$$
y_i = \sum_{j=1}^{p} \left( \left[\beta_j'^{\text{Eur}} (1 - \overline{a_{i\cdot}}) + \beta_j'^{\text{Afr}} \overline{a_{i\cdot}}\right] \hat{x}_{ij}'^{(1)} + \left[\beta_j'^{\text{Eur}} (1 - \overline{a_{i\cdot}}) + \beta_j'^{\text{Afr}} \overline{a_{i\cdot}}\right] \hat{x}_{ij}'^{(2)} \right) + \varepsilon_i.
$$

Again, we drop the exogenous noise term $\varepsilon_i$ from our calculations. By independence of markers and algebraic rules of the sample covariance, we obtain (note $\overline{\boldsymbol{a_{\cdot\cdot}}} = (\overline{a_{i\cdot}} : i = 1, \ldots, n)$)

$$\begin{aligned}
\mathrm{Cov}\left(\mathbf{TotPGS}, \boldsymbol{y}\right) &= \sum_{j=1}^{p}\Bigg\{\underbrace{\mathrm{Cov}\left(\beta_j'^{\mathrm{Eur}}\hat{\boldsymbol{x}}_{\boldsymbol{\cdot j}}'^{(1)}, \beta_j'^{\mathrm{Eur}}(\mathbf{1}-\overline{\boldsymbol{a_{\cdot\cdot}}})\hat{\boldsymbol{x}}_{\boldsymbol{\cdot j}}'^{(1)} + \beta_j'^{\mathrm{Afr}}\overline{\boldsymbol{a_{\cdot\cdot}}}\hat{\boldsymbol{x}}_{\boldsymbol{\cdot j}}'^{(1)}\right)}_{\text{(7G')}} \\
&\quad + \underbrace{\mathrm{Cov}\left(\beta_j'^{\mathrm{Eur}}\hat{\boldsymbol{x}}_{\boldsymbol{\cdot j}}'^{(1)}, \beta_j'^{\mathrm{Eur}}(\mathbf{1}-\overline{\boldsymbol{a_{\cdot\cdot}}})\hat{\boldsymbol{x}}_{\boldsymbol{\cdot j}}'^{(2)} + \beta_j'^{\mathrm{Afr}}\overline{\boldsymbol{a_{\cdot\cdot}}}\hat{\boldsymbol{x}}_{\boldsymbol{\cdot j}}'^{(2)}\right)}_{\text{(8G')}} \\
&\quad + \underbrace{\mathrm{Cov}\left(\beta_j'^{\mathrm{Eur}}\hat{\boldsymbol{x}}_{\boldsymbol{\cdot j}}'^{(2)}, \beta_j'^{\mathrm{Eur}}(\mathbf{1}-\overline{\boldsymbol{a_{\cdot\cdot}}})\hat{\boldsymbol{x}}_{\boldsymbol{\cdot j}}'^{(1)} + \beta_j'^{\mathrm{Afr}}\overline{\boldsymbol{a_{\cdot\cdot}}}\hat{\boldsymbol{x}}_{\boldsymbol{\cdot j}}'^{(1)}\right)}_{\text{(9G')}} \\
&\quad + \underbrace{\mathrm{Cov}\left(\beta_j'^{\mathrm{Eur}}\hat{\boldsymbol{x}}_{\boldsymbol{\cdot j}}'^{(2)}, \beta_j'^{\mathrm{Eur}}(\mathbf{1}-\overline{\boldsymbol{a_{\cdot\cdot}}})\hat{\boldsymbol{x}}_{\boldsymbol{\cdot j}}'^{(2)} + \beta_j'^{\mathrm{Afr}}\overline{\boldsymbol{a_{\cdot\cdot}}}\hat{\boldsymbol{x}}_{\boldsymbol{\cdot j}}'^{(2)}\right)}_{\text{(10G')}}\Bigg\}
\end{aligned}$$

The four terms simplify as follows:

$$\text{(7G')} = \underbrace{\left(\beta_j'^{\mathrm{Eur}}\right)^2\frac{1}{n-1}\sum_{i=1}^{n}(1-\overline{a_{i\cdot}})\left(\hat{x}_{ij}'^{(1)}\right)^2}_{\text{(7G')}_1} + \underbrace{\beta_j'^{\mathrm{Eur}}\beta_j'^{\mathrm{Afr}}\frac{1}{n-1}\sum_{i=1}^{n}\overline{a_{i\cdot}}\left(\hat{x}_{ij}'^{(1)}\right)^2}_{\text{(7G')}_2},$$

$$\text{(8G')} = \underbrace{\left(\beta_j'^{\mathrm{Eur}}\right)^2\frac{1}{n-1}\sum_{i=1}^{n}(1-\overline{a_{i\cdot}})\,\hat{x}_{ij}'^{(1)}\hat{x}_{ij}'^{(2)}}_{\text{(8G')}_1} + \underbrace{\beta_j'^{\mathrm{Eur}}\beta_j'^{\mathrm{Afr}}\frac{1}{n-1}\sum_{i=1}^{n}\overline{a_{i\cdot}}\hat{x}_{ij}'^{(1)}\hat{x}_{ij}'^{(2)}}_{\text{(8G')}_2},$$

$$\text{(9G')} = \underbrace{\left(\beta_j'^{\mathrm{Eur}}\right)^2\frac{1}{n-1}\sum_{i=1}^{n}(1-\overline{a_{i\cdot}})\,\hat{x}_{ij}'^{(1)}\hat{x}_{ij}'^{(2)}}_{\text{(9G')}_1} + \underbrace{\beta_j'^{\mathrm{Eur}}\beta_j'^{\mathrm{Afr}}\frac{1}{n-1}\sum_{i=1}^{n}\overline{a_{i\cdot}}\hat{x}_{ij}'^{(1)}\hat{x}_{ij}'^{(2)}}_{\text{(9G')}_2},$$

$$\text{(10G')} = \underbrace{\left(\beta_j'^{\mathrm{Eur}}\right)^2\frac{1}{n-1}\sum_{i=1}^{n}(1-\overline{a_{i\cdot}})\left(\hat{x}_{ij}'^{(2)}\right)^2}_{\text{(10G')}_1} + \underbrace{\beta_j'^{\mathrm{Eur}}\beta_j'^{\mathrm{Afr}}\frac{1}{n-1}\sum_{i=1}^{n}\overline{a_{i\cdot}}\left(\hat{x}_{ij}'^{(2)}\right)^2}_{\text{(10G')}_2}.$$

**Covariance of ParPGS with Phenotype.** Again, these depend on the model assumed, so we work out the expressions under the global model and the local model.

*Local Model.* Similar to the arguments for obtaining the covariance of TotPGS with phenotype, we leverage independence of markers, independence of the exogenous noise term $\varepsilon_i$, and algebraic rules of the sample covariance, to obtain

$$
\begin{aligned}
\operatorname{Cov}\left(\mathbf{ParPGS}, \boldsymbol{y}\right) &= \sum_{j=1}^{p}\Bigg\{\underbrace{\operatorname{Cov}\left(\beta_j^{\prime\text{Eur}}\left(\mathbf{1}-\boldsymbol{a}_{\cdot j}^{\prime(1)}\right)\hat{\boldsymbol{x}}_{\cdot j}^{\prime(1)}, \beta_j^{\prime\text{Eur}}\left(\mathbf{1}-\boldsymbol{a}_{\cdot j}^{\prime(1)}\right)\hat{\boldsymbol{x}}_{\cdot j}^{\prime(1)} + \beta_j^{\prime\text{Afr}}\boldsymbol{a}_{\cdot j}^{\prime(1)}\hat{\boldsymbol{x}}_{\cdot j}^{\prime(1)}\right)}_{\text{(11L)}} \\
&\quad + \underbrace{\operatorname{Cov}\left(\beta_j^{\prime\text{Eur}}\left(\mathbf{1}-\boldsymbol{a}_{\cdot j}^{\prime(1)}\right)\hat{\boldsymbol{x}}_{\cdot j}^{\prime(1)}, \beta_j^{\prime\text{Eur}}\left(\mathbf{1}-\boldsymbol{a}_{\cdot j}^{\prime(2)}\right)\hat{\boldsymbol{x}}_{\cdot j}^{\prime(2)} + \beta_j^{\prime\text{Afr}}\boldsymbol{a}_{\cdot j}^{\prime(2)}\hat{\boldsymbol{x}}_{\cdot j}^{\prime(2)}\right)}_{\text{(12L')}} \\
&\quad + \underbrace{\operatorname{Cov}\left(\beta_j^{\prime\text{Eur}}\left(\mathbf{1}-\boldsymbol{a}_{\cdot j}^{\prime(2)}\right)\hat{\boldsymbol{x}}_{\cdot j}^{\prime(2)}, \beta_j^{\prime\text{Eur}}\left(\mathbf{1}-\boldsymbol{a}_{\cdot j}^{\prime(1)}\right)\hat{\boldsymbol{x}}_{\cdot j}^{\prime(1)} + \beta_j^{\prime\text{Afr}}\boldsymbol{a}_{\cdot j}^{\prime(1)}\hat{\boldsymbol{x}}_{\cdot j}^{\prime(1)}\right)}_{\text{(13L')}} \\
&\quad + \underbrace{\operatorname{Cov}\left(\beta_j^{\prime\text{Eur}}\left(\mathbf{1}-\boldsymbol{a}_{\cdot j}^{\prime(2)}\right)\hat{\boldsymbol{x}}_{\cdot j}^{\prime(2)}, \beta_j^{\prime\text{Eur}}\left(\mathbf{1}-\boldsymbol{a}_{\cdot j}^{\prime(2)}\right)\hat{\boldsymbol{x}}_{\cdot j}^{\prime(2)} + \beta_j^{\prime\text{Afr}}\boldsymbol{a}_{\cdot j}^{\prime(2)}\hat{\boldsymbol{x}}_{\cdot j}^{\prime(2)}\right)}_{\text{(14L')}}\Bigg\}
\end{aligned}
$$

The four terms simplify as follows:

$$
\begin{aligned}
\text{(11L')} &= \left(\beta_j^{\prime\text{Eur}}\right)^2\frac{n}{n-1}\left(1-\overline{a_{\cdot j}^{\prime(1)}}\right)\hat{f}_j^{\prime\text{Eur}}\left(1-\hat{f}_j^{\prime\text{Eur}}\right)+0=\text{(4')}, \\
\text{(12L')} &= \text{(6')} + \underbrace{\beta_j^{\prime\text{Eur}}\beta_j^{\prime\text{Afr}}\frac{1}{n-1}\sum_{i=1}^{n}a_{ij}^{\prime(2)}\left(1-a_{ij}^{\prime(1)}\right)\hat{x}_{ij}^{\prime(1)}\hat{x}_{ij}^{\prime(2)}}_{\text{(12L')}_2}, \\
\text{(13L')} &= \text{(6')} + \underbrace{\beta_j^{\prime\text{Eur}}\beta_j^{\prime\text{Afr}}\frac{1}{n-1}\sum_{i=1}^{n}a_{ij}^{\prime(1)}\left(1-a_{ij}^{\prime(2)}\right)\hat{x}_{ij}^{\prime(1)}\hat{x}_{ij}^{\prime(2)}}_{\text{(13L')}_2}, \\
\text{(14L')} &= \left(\beta_j^{\prime\text{Eur}}\right)^2\frac{n}{n-1}\left(1-\overline{a_{\cdot j}^{\prime(2)}}\right)\hat{f}_j^{\prime\text{Eur}}\left(1-\hat{f}_j^{\prime\text{Eur}}\right)+0=\text{(5')}.
\end{aligned}
$$

Note that the simplifications of (11L') and (14L') rely on the fact that $\left(1-a_{ij}^{\prime(h)}\right)a_{ij}^{\prime(h)}=0$.

*Global Model.* Similar to the arguments for obtaining the covariance of TotPGS with phenotype, we leverage independence of markers, independence of the exogenous noise term $\varepsilon_i$, and algebraic rules of the sample covariance, to obtain

$$
\begin{aligned}
\operatorname{Cov}\left(\mathbf{ParPGS}, \boldsymbol{y}\right) &= \sum_{j=1}^{p} \Bigg\{ \underbrace{\operatorname{Cov}\left(\beta_j^{\text{Eur}}\left(\mathbf{1}-\boldsymbol{a}'^{(1)}_{\cdot j}\right)\hat{\boldsymbol{x}}'^{(1)}_{\cdot j}, \beta_j'^{\text{Eur}}\left(\mathbf{1}-\overline{\boldsymbol{a}_{\cdot\cdot}}\right)\hat{\boldsymbol{x}}'^{(1)}_{\cdot j} + \beta_j'^{\text{Afr}}\overline{\boldsymbol{a}_{\cdot\cdot}}\hat{\boldsymbol{x}}'^{(1)}_{\cdot j}\right)}_{\text{(11G')}} \\
&\quad + \underbrace{\operatorname{Cov}\left(\beta_j'^{\text{Eur}}\left(\mathbf{1}-\boldsymbol{a}'^{(1)}_{\cdot j}\right)\hat{\boldsymbol{x}}'^{(1)}_{\cdot j}, \beta_j'^{\text{Eur}}\left(\mathbf{1}-\overline{\boldsymbol{a}_{\cdot\cdot}}\right)\hat{\boldsymbol{x}}'^{(2)}_{\cdot j} + \beta_j'^{\text{Afr}}\overline{\boldsymbol{a}_{\cdot\cdot}}\hat{\boldsymbol{x}}'^{(2)}_{\cdot j}\right)}_{\text{(12G')}} \\
&\quad + \underbrace{\operatorname{Cov}\left(\beta_j'^{\text{Eur}}\left(\mathbf{1}-\boldsymbol{a}'^{(2)}_{\cdot j}\right)\hat{\boldsymbol{x}}'^{(2)}_{\cdot j}, \beta_j^{\text{Eur}}\left(\mathbf{1}-\overline{\boldsymbol{a}_{\cdot\cdot}}\right)\hat{\boldsymbol{x}}'^{(1)}_{\cdot j} + \beta_j^{\text{Afr}}\overline{\boldsymbol{a}_{\cdot\cdot}}\hat{\boldsymbol{x}}'^{(1)}_{\cdot j}\right)}_{\text{(13G')}} \\
&\quad + \underbrace{\operatorname{Cov}\left(\beta_j'^{\text{Eur}}\left(\mathbf{1}-\boldsymbol{a}'^{(2)}_{\cdot j}\right)\hat{\boldsymbol{x}}'^{(2)}_{\cdot j}, \beta_j'^{\text{Eur}}\left(\mathbf{1}-\overline{\boldsymbol{a}_{\cdot\cdot}}\right)\hat{\boldsymbol{x}}'^{(2)}_{\cdot j} + \beta_j'^{\text{Afr}}\overline{\boldsymbol{a}_{\cdot\cdot}}\hat{\boldsymbol{x}}'^{(2)}_{\cdot j}\right)}_{\text{(14G')}} \Bigg\}
\end{aligned}
$$

The four terms simplify as follows:

$$
\begin{aligned}
\text{(11G')} &= \underbrace{\left(\beta_j'^{\text{Eur}}\right)^2 \frac{1}{n-1}\sum_{i=1}^{n}\left(1-a'^{(1)}_{ij}\right)\left(1-\overline{a_{i\cdot}}\right)\left(\hat{x}'^{(1)}_{ij}\right)^2}_{\text{(11G')}_1} \\
&\quad + \underbrace{\beta_j'^{\text{Eur}}\beta_j'^{\text{Afr}}\frac{1}{n-1}\sum_{i=1}^{n}\left(1-a'^{(1)}_{ij}\right)\overline{a_{i\cdot}}\left(\hat{x}'^{(1)}_{ij}\right)^2}_{\text{(11G')}_2},
\end{aligned}
$$

$$
\begin{aligned}
\text{(12G')} &= \underbrace{\left(\beta_j'^{\text{Eur}}\right)^2 \frac{1}{n-1}\sum_{i=1}^{n}\left(1-a'^{(1)}_{ij}\right)\left(1-\overline{a_{i\cdot}}\right)\hat{x}'^{(1)}_{ij}\hat{x}'^{(2)}_{ij}}_{\text{(12G')}_1} \\
&\quad + \underbrace{\beta_j'^{\text{Eur}}\beta_j'^{\text{Afr}}\frac{1}{n-1}\sum_{i=1}^{n}\left(1-a'^{(1)}_{ij}\right)\overline{a_{i\cdot}}\hat{x}'^{(1)}_{ij}\hat{x}'^{(2)}_{ij}}_{\text{(12G')}_2},
\end{aligned}
$$

$$
\textcircled{13G'} = \underbrace{\left(\beta_j'^{\text{Eur}}\right)^2 \frac{1}{n-1} \sum_{i=1}^{n} \left(1 - a_{ij}'^{(2)}\right) \left(1 - \overline{a_{i\cdot}}\right) \hat{x}_{ij}'^{(1)} \hat{x}_{ij}'^{(2)}}_{\textcircled{13G'}_1}
+ \underbrace{\beta_j'^{\text{Eur}} \beta_j'^{\text{Afr}} \frac{1}{n-1} \sum_{i=1}^{n} \left(1 - a_{ij}'^{(2)}\right) \overline{a_{i\cdot}} \hat{x}_{ij}'^{(1)} \hat{x}_{ij}'^{(2)}}_{\textcircled{13G}_2},
$$

$$
\textcircled{14G'} = \underbrace{\left(\beta_j'^{\text{Eur}}\right)^2 \frac{1}{n-1} \sum_{i=1}^{n} \left(1 - a_{ij}'^{(2)}\right) \left(1 - \overline{a_{i\cdot}}\right) \left(\hat{x}_{ij}'^{(2)}\right)^2}_{\textcircled{14G'}_1}
+ \underbrace{\beta_j'^{\text{Eur}} \beta_j'^{\text{Afr}} \frac{1}{n-1} \sum_{i=1}^{n} \left(1 - a_{ij}'^{(2)}\right) \overline{a_{i\cdot}} \left(\hat{x}_{ij}'^{(2)}\right)^2}_{\textcircled{14G'}_2}.
$$

We summarize all of the above calculations of PGS-phenotype correlations in **Box A** and **Box B** for easy reference.

**Box A: Correlations Under the Local Model when Causal Variants are Known**

$$
\text{Cor}(\mathbf{TotPGS}, \boldsymbol{y}) = \frac{\sum_{j=1}^{p} \left(\textcircled{7L'} + \textcircled{8L'} + \textcircled{9L'} + \textcircled{10L'}\right)}{\sqrt{\sum_{j=1}^{p} \left(\textcircled{1'} + \textcircled{2'} + \textcircled{3'}\right)}}
$$

$$
\text{Cor}(\mathbf{ParPGS}, \boldsymbol{y}) = \frac{\sum_{j=1}^{p} \left(\textcircled{11L'} + \textcircled{12L'} + \textcircled{13L'} + \textcircled{14L'}\right)}{\sqrt{\sum_{j=1}^{p} \left(\textcircled{4'} + \textcircled{5'} + \textcircled{6'}\right)}}
$$

**Box B: Correlations Under the Global Model when Causal Variants are Known**

$$\text{Cor}(\mathbf{TotPGS}, \boldsymbol{y}) = \frac{\sum_{j=1}^{p}\left(\textcircled{7G'} + \textcircled{8G'} + \textcircled{9G'} + \textcircled{10G'}\right)}{\sqrt{\sum_{j=1}^{p}\left(\textcircled{1'} + \textcircled{2'} + \textcircled{3'}\right)}}$$

$$\text{Cor}(\mathbf{ParPGS}, \boldsymbol{y}) = \frac{\sum_{j=1}^{p}\left(\textcircled{11G'} + \textcircled{12G'} + \textcircled{13G'} + \textcircled{14G'}\right)}{\sqrt{\sum_{j=1}^{p}\left(\textcircled{4'} + \textcircled{5'} + \textcircled{6'}\right)}}$$

Step 2: Expectation of quantities under distributional assumptions of genetic effects and noise. To compute the expected correlations, we rely on the approximation $\mathbb{E}[X/Y] \approx \mathbb{E}[X]/\mathbb{E}[Y]$, which is not strong in general but works well in our setting, as we have verified through simulations.[1] We also rely on the approximation $\mathbb{E}[\sqrt{X}] \approx \sqrt{\mathbb{E}[X]}$.[2] Thus, we have, by linearity of expectation,

$$\mathbb{E}_{\text{Loc}}\left[\text{Cor}(\mathbf{TotPGS}, \boldsymbol{y})\right] \approx \frac{\sum_{j=1}^{p}\left(\mathbb{E}\left[\textcircled{7L'}\right] + \mathbb{E}\left[\textcircled{8L'}\right] + \mathbb{E}\left[\textcircled{9L'}\right] + \mathbb{E}\left[\textcircled{10L'}\right]\right)}{\sqrt{\sum_{j=1}^{p}\left(\mathbb{E}\left[\textcircled{1'}\right] + \mathbb{E}\left[\textcircled{2'}\right] + \mathbb{E}\left[\textcircled{3'}\right]\right)}}, \quad \text{(S10)}$$

$$\mathbb{E}_{\text{Loc}}\left[\text{Cor}(\mathbf{ParPGS}, \boldsymbol{y})\right] \approx \frac{\sum_{j=1}^{p}\left(\mathbb{E}\left[\textcircled{11L'}\right] + \mathbb{E}\left[\textcircled{12L'}\right] + \mathbb{E}\left[\textcircled{13L'}\right] + \mathbb{E}\left[\textcircled{14L'}\right]\right)}{\sqrt{\sum_{j=1}^{p}\left(\mathbb{E}\left[\textcircled{4'}\right] + \mathbb{E}\left[\textcircled{5'}\right] + \mathbb{E}\left[\textcircled{6'}\right]\right)}}. \quad \text{(S11)}$$

Similarly for the global model,

$$\mathbb{E}_{\text{Glo}}\left[\text{Cor}(\mathbf{TotPGS}, \boldsymbol{y})\right] \approx \frac{\sum_{j=1}^{p}\left(\mathbb{E}\left[\textcircled{7G'}\right] + \mathbb{E}\left[\textcircled{8G'}\right] + \mathbb{E}\left[\textcircled{9G'}\right] + \mathbb{E}\left[\textcircled{10G'}\right]\right)}{\sqrt{\sum_{j=1}^{p}\left(\mathbb{E}\left[\textcircled{1'}\right] + \mathbb{E}\left[\textcircled{2'}\right] + \mathbb{E}\left[\textcircled{3'}\right]\right)}}, \quad \text{(S12)}$$

$$\mathbb{E}_{\text{Glo}}\left[\text{Cor}(\mathbf{ParPGS}, \boldsymbol{y})\right] \approx \frac{\sum_{j=1}^{p}\left(\mathbb{E}\left[\textcircled{11G'}\right] + \mathbb{E}\left[\textcircled{12G'}\right] + \mathbb{E}\left[\textcircled{13G'}\right] + \mathbb{E}\left[\textcircled{14G'}\right]\right)}{\sqrt{\sum_{j=1}^{p}\left(\mathbb{E}\left[\textcircled{4'}\right] + \mathbb{E}\left[\textcircled{5'}\right] + \mathbb{E}\left[\textcircled{6'}\right]\right)}}. \quad \text{(S13)}$$

To simplify these expectations (Eqs. (S10)-(S13)) into expressions involving only fixed quantities, we must evaluate the following expectations: $\mathbb{E}\left[\left(\beta_j'^{\text{Eur}}\right)^2\right]$ and $\mathbb{E}\left[\beta_j'^{\text{Eur}}\beta_j'^{\text{Afr}}\right]$. By Eqs. (2)-(5) these are $\sigma_{\text{Eur}}'^2$ and $\tau'$ respectively. Therefore, expectations of quantities are computed by substituting each occurrence of the relevant expectation into Eqs. (S10)-(S13).

[1]A better approximation, derived via Taylor expansion, is $\mathbb{E}[X/Y] \approx \mathbb{E}[X]/\mathbb{E}[Y] - \text{cov}(X,Y)/(\mathbb{E}[Y])^2 + \left[\text{var}(Y)\mathbb{E}[X]/(\mathbb{E}[Y])^3\right]$, but this would require tedious second moment calculations.

[2]Again, a better but more tedious approximation is $\mathbb{E}[\sqrt{X}] \approx \sqrt{\mathbb{E}[X]} - \text{var}(X)/(8\mathbb{E}[X]\sqrt{E[X]})$.

#### S5 Causal Variant Polygenic Score Behaviour Under Additional Assumptions

This section proves Proposition 4.2 of the Main Text, which states the following conclusions under some additional assumptions about the causal variants (which are furthermore assumed known).

1. The squared phenotype correlations with TotPGS are approximately equal between the local and the global models, and are a quadratic function of global African ancestry.
2. Under the global model, the squared phenotytpe correlation with ParPGS is approximately a cubic function of global African ancestry.
3. Under the local model, the squared phenotype correlation with ParPGS is approximately a linear function of global African ancestry.

The additional assumptions are as follows.

(A) Ancestry-specific allele frequencies are identical across markers.

(B) Distribution of African ancestries is the same across individuals.

(C) Ancestry assignment is identical between two haplotypes of a diploid individual.

(D) At each marker, the difference between ancestry-specific allele frequencies is small.

(E) Distribution of alleles is approximately uncorrelated between two haplotypes at a diploid locus.

We note that these assumptions do not hold in practice, and are assumed for mathematical convenience. To arrive at the conclusions of Proposition 4.2, we will first state what each assumption entails mathematically (Step 1; all involve simplifying complicated quantities), and then apply them to simplify the expressions (1') - (6'), (7L') - (14L'), (7G') - (14G'), $\sigma'^2_{\text{Eur}}$, $\sigma'^2_{\text{Afr}}$ and $\tau'$ (Step 2). From there, arriving at the quantities stated in Proposition 4.2 is straightforward (Step 3).

Step 1: Mathematical entailment of additional assumptions.

(A) This means that

$$\begin{aligned} \hat{f}_1'^{\text{Eur}} = \hat{f}_2'^{\text{Eur}} = \ldots = \hat{f}_p'^{\text{Eur}} &= \hat{f}'^{\text{Eur}}, \\ \hat{f}_1'^{\text{Afr}} = \hat{f}_2'^{\text{Afr}} = \ldots = \hat{f}_p'^{\text{Afr}} &= \hat{f}'^{\text{Afr}}. \end{aligned}$$

(B) This means that

$$a_{1j}'^{(h)} = a_{2j}'^{(h)} = \ldots = a_{nj}'^{(h)} = a_j'^{(h)}$$

for each haplotype $h$ and marker $j$. Specifically,

$$a_j'^{(h)} = \begin{cases} 1 & \text{if marker } j \text{ on haplotype } h \text{ is assigned African ancestry} \\ 0 & \text{otherwise} \end{cases}$$

(C) This means that $a_{ij}'^{(1)} = a_{ij}'^{(2)}$ for each individual $i$ at marker $j$. Combined with Assumption (B), this means that

$$a_j'^{(1)} = a_j'^{(2)} = a_j' = \begin{cases} 1 & \text{if marker } j \text{ is assigned African ancestry} \\ 0 & \text{otherwise} \end{cases}$$

(D) This means that $\hat{f}_j^{\prime\text{Eur}} - \hat{f}_j^{\prime\text{Afr}} = o\left(\min\{\hat{f}_j^{\prime\text{Eur}}, \hat{f}_j^{\prime\text{Afr}}\}\right)$.[3] Note that $o(\cdot)$ is "little $o$" with respect to sample size $n$, which is to say that

$$\lim_{n\to\infty}\left(\frac{\hat{f}^{\prime\text{Eur}} - \hat{f}^{\prime\text{Afr}}}{\min\{\hat{f}^{\prime\text{Eur}}, \hat{f}^{\prime\text{Afr}}\}}\right) = 0.$$

(E) Combined with Assumptions (B) and (C), this means that for any marker $j$,

$$\sum_{i=1}^{n} \hat{x}_{ij}^{\prime(1)} \hat{x}_{ij}^{\prime(2)} \approx 0.$$

Step 2: Application of Step 1 to expressions.

**Simplifying ① - ⑥.** We obtain the following simplified expressions.

$$\begin{aligned}
① &= \left(\beta_j^{\prime\text{Eur}}\right)^2 \frac{n}{n-1}\left[(1-a_j')\hat{f}^{\prime\text{Eur}}\left(1-\hat{f}^{\prime\text{Eur}}\right) + a_j'\hat{f}^{\prime\text{Afr}}\left(1-\hat{f}^{\prime\text{Afr}}\right)\right]\\
② &= \left(\beta_j^{\prime\text{Eur}}\right)^2 \frac{n}{n-1}\left[(1-a_j')\hat{f}^{\prime\text{Eur}}\left(1-\hat{f}^{\prime\text{Eur}}\right) + a_j'\hat{f}^{\prime\text{Afr}}\left(1-\hat{f}^{\prime\text{Afr}}\right)\right]\\
③ &\approx 0\\
④ &= \left(\beta_j^{\prime\text{Eur}}\right)^2 \frac{n}{n-1}(1-a_j)\hat{f}^{\prime\text{Eur}}\left(1-\hat{f}^{\prime\text{Eur}}\right)\\
⑤ &= \left(\beta_j^{\prime\text{Eur}}\right)^2 \frac{n}{n-1}(1-a_j)\hat{f}^{\prime\text{Eur}}\left(1-\hat{f}^{\prime\text{Eur}}\right)\\
(6') &\approx 0
\end{aligned}$$

**Simplifying (7L') - (14L').** We obtain the following simplified expressions.

$$\begin{aligned}
(7L') &= \frac{n}{n-1}\left[\left(\beta_j^{\prime\text{Eur}}\right)^2(1-a_j')\hat{f}^{\prime\text{Eur}}\left(1-\hat{f}^{\prime\text{Eur}}\right) + \beta_j^{\prime\text{Eur}}\beta_j^{\prime\text{Afr}}a_j\hat{f}^{\prime\text{Afr}}\left(1-\hat{f}^{\prime\text{Afr}}\right)\right]\\
(8L') &\approx 0\\
(9L') &\approx 0\\
(10L') &= \frac{n}{n-1}\left[\left(\beta_j^{\prime\text{Eur}}\right)^2(1-a_j')\hat{f}^{\prime\text{Eur}}\left(1-\hat{f}^{\prime\text{Eur}}\right) + \beta_j^{\prime\text{Eur}}\beta_j^{\prime\text{Afr}}a_j'\hat{f}^{\prime\text{Afr}}\left(1-\hat{f}^{\prime\text{Afr}}\right)\right]
\end{aligned}$$

$$\begin{aligned}
(11L') &= \left(\beta_j^{\prime\text{Eur}}\right)^2 \frac{n}{n-1}(1-a_j')\hat{f}^{\prime\text{Eur}}\left(1-\hat{f}^{\prime\text{Eur}}\right)\\
(12L') &\approx 0\\
(13L') &\approx 0\\
(14L') &= \left(\beta_j^{\prime\text{Eur}}\right)^2 \frac{n}{n-1}(1-a_j')\hat{f}^{\prime\text{Eur}}\left(1-\hat{f}^{\prime\text{Eur}}\right)
\end{aligned}$$

[3] With probability 1, since allele frequencies are, technically speaking, finite-sample estimates. We will ignore this detail.

**Simplifying (7G') - (14G').** We obtain the following simplified expressions.

$$\begin{aligned}
\text{(7G')} &= \frac{n}{n-1}\left[\left(\beta_j'^{\text{Eur}}\right)^2(1-\overline{a})+\beta_j'^{\text{Eur}}\beta_j'^{\text{Afr}}\overline{a}\right]\left[(1-a_j')\hat{f}'^{\text{Eur}}\left(1-\hat{f}'^{\text{Eur}}\right)+a_j'\hat{f}'^{\text{Afr}}\left(1-\hat{f}'^{\text{Afr}}\right)\right] \\
\text{(8G')} &\approx 0 \\
\text{(9G')} &\approx 0 \\
\text{(10G')} &= \frac{n}{n-1}\left[\left(\beta_j'^{\text{Eur}}\right)^2(1-\overline{a})+\beta_j'^{\text{Eur}}\beta_j'^{\text{Afr}}\overline{a}\right]\left[(1-a_j')\hat{f}'^{\text{Eur}}\left(1-\hat{f}'^{\text{Eur}}\right)+a_j'\hat{f}'^{\text{Afr}}\left(1-\hat{f}'^{\text{Afr}}\right)\right] \\
\text{(11G')} &= \frac{n}{n-1}\left[\left(\beta_j'^{\text{Eur}}\right)^2(1-\overline{a})+\beta_j'^{\text{Eur}}\beta_j'^{\text{Afr}}\overline{a}\right](1-a_j')\hat{f}'^{\text{Eur}}\left(1-\hat{f}'^{\text{Eur}}\right) \\
\text{(12G')} &\approx 0 \\
\text{(13G')} &\approx 0 \\
\text{(14G')} &= \frac{n}{n-1}\left[\left(\beta_j'^{\text{Eur}}\right)^2(1-\overline{a})+\beta_j'^{\text{Eur}}\beta_j'^{\text{Afr}}\overline{a}\right](1-a_j')\hat{f}'^{\text{Eur}}\left(1-\hat{f}'^{\text{Eur}}\right)
\end{aligned}$$

**Simplifying the remaining quantities.** We obtain the following simplified expressions.

$$\begin{aligned}
\sigma'^2_{\text{Eur}} &= \frac{r^2}{2p\hat{f}'^{\text{Eur}}\left(1-\hat{f}'^{\text{Eur}}\right)} \\
\sigma'^2_{\text{Afr}} &= \frac{r^2}{2p\hat{f}'^{\text{Afr}}\left(1-\hat{f}'^{\text{Afr}}\right)} \\
\tau' &= \frac{r^2\rho}{2p\sqrt{\hat{f}'^{\text{Eur}}\hat{f}'^{\text{Afr}}\left(1-\hat{f}'^{\text{Eur}}\right)\left(1-\hat{f}'^{\text{Afr}}\right)}}
\end{aligned}$$

Step 3: Verifying quantities reported in Proposition 4.2. We plug the simplified expressions from Step 2 directly into the relevant quantities.

**Expected correlations between phenotype and TotPGS ($\mathbb{E}$[Cor(TotPGS, $\boldsymbol{y}$)]).** Under the local model, the numerator of Eq. (S10) is

$$\begin{aligned}
\textcircled{\scriptsize T}^{\text{Num}}_{\text{Loc}} &\approx \frac{n}{n-1}\Bigg\{\hat{f}'^{\text{Eur}}\left(1-\hat{f}'^{\text{Eur}}\right)\sum_{j=1}^{p}(1-a_j')\left[\mathbb{E}\left[\left(\beta_j'^{\text{Eur}}\right)^2\right]\right. \\
&\qquad \left. +\ \mathbb{E}\left[\left(\beta_j'^{\text{Eur}}\right)^2\right]\right] + \hat{f}'^{\text{Afr}}\left(1-\hat{f}'^{\text{Afr}}\right)\sum_{j=1}^{p}a_j'\left[\mathbb{E}\left[\beta_j'^{\text{Eur}}\beta_j'^{\text{Afr}}\right]+\mathbb{E}\left[\beta_j'^{\text{Eur}}\beta_j'^{\text{Afr}}\right]\right]\Bigg\} \\
&= \frac{n}{n-1}\left[2\sigma_{\text{Eur}}'^2\hat{f}'^{\text{Eur}}\left(1-\hat{f}'^{\text{Eur}}\right)p(1-\overline{a'})+2\tau'\hat{f}'^{\text{Afr}}\left(1-\hat{f}'^{\text{Afr}}\right)p\overline{a'}\right] \\
&= \frac{n}{n-1}\left[r^2(1-\overline{a})+r^2\rho\overline{a}\sqrt{\frac{\hat{f}'^{\text{Afr}}\left(1-\hat{f}'^{\text{Afr}}\right)}{\hat{f}'^{\text{Eur}}\left(1-\hat{f}'^{\text{Eur}}\right)}}\right],
\end{aligned}$$

where we used the assumption that global ancestry calculated across causal variants is the same as if it were calculated across tagging variants ($\overline{a} = \overline{a'}$) in the last equality. The last expression can be further approximated as

$$\frac{n}{n-1}\left[r^2(1-\overline{a})+r^2\rho\overline{a}(1+o(1))\right] \approx r^2(1-\overline{a}+\rho\overline{a}).$$

The squared denominator is

$$\begin{aligned}
\left[\textcircled{\scriptsize T}^{\text{Den}}_{\text{Loc}}\right]^2 &\approx \frac{n}{n-1}\Bigg\{\hat{f}'^{\text{Eur}}\left(1-\hat{f}'^{\text{Eur}}\right)\sum_{j=1}^{p}(1-a_j')\left[\mathbb{E}\left[\left(\beta_j'^{\text{Eur}}\right)^2\right]\right. \\
&\qquad \left. +\ \mathbb{E}\left[\left(\beta_j'^{\text{Eur}}\right)^2\right]\right] + \hat{f}'^{\text{Afr}}\left(1-\hat{f}'^{\text{Afr}}\right)\sum_{j=1}^{p}a_j'\left[\mathbb{E}\left[\left(\beta_j'^{\text{Eur}}\right)^2\right]+\mathbb{E}\left[\left(\beta_j'^{\text{Eur}}\right)^2\right]\right]\Bigg\} \\
&= \frac{n}{n-1}2\sigma_{\text{Eur}}'^2\left[\hat{f}'^{\text{Eur}}\left(1-\hat{f}'^{\text{Eur}}\right)p(1-\overline{a'})+\hat{f}'^{\text{Afr}}\left(1-\hat{f}'^{\text{Afr}}\right)p\overline{a'}\right] \\
&= \frac{n}{n-1}\left[r^2(1-\overline{a})+r^2\overline{a}\frac{\hat{f}'^{\text{Afr}}\left(1-\hat{f}'^{\text{Afr}}\right)}{\hat{f}'^{\text{Eur}}\left(1-\hat{f}'^{\text{Eur}}\right)}\right] \\
&= \frac{n}{n-1}\left[r^2(1-\overline{a})+r^2\overline{a}(1+o(1))^2\right] \\
&\approx r^2
\end{aligned}$$

Therefore

$$\mathbb{E}_{\text{Loc}}\left[\text{Cor}(\textbf{TotPGS}, \boldsymbol{y})\right] \approx \frac{\textcircled{\scriptsize T}^{\text{Num}}_{\text{Loc}}}{\textcircled{\scriptsize T}^{\text{Den}}_{\text{Loc}}} = r(1-\overline{a}+\rho\overline{a}),$$

and using $\mathbb{E}[X] \approx (\mathbb{E}[\sqrt{X}])^2$, $\mathbb{E}_{\text{Loc}}\left[\text{Cor}^2(\textbf{TotPGS}, \boldsymbol{y})\right] \approx r^2(1-\overline{a}+\rho\overline{a})^2$.

Performing a similar calculation under the global model, we have

$$\begin{aligned}
\textcircled{T}^{\text{Num}}_{\text{Glo}} &\approx \frac{n}{n-1}\left[2\sigma'^2_{\text{Eur}}(1-\overline{a}) + 2\tau'\overline{a}\right] \\
&\quad \times \left[\hat{f}'^{\text{Eur}}\left(1-\hat{f}'^{\text{Eur}}\right)\sum_{j=1}^{p}(1-a'_j) + \hat{f}'^{\text{Afr}}\left(1-\hat{f}'^{\text{Afr}}\right)\sum_{j=1}^{p}a'_j\right] \\
&= \frac{n}{n-1}\left[\frac{r^2(1-\overline{a})}{p\hat{f}'^{\text{Eur}}\left(1-\hat{f}'^{\text{Eur}}\right)} + \frac{\rho r^2\overline{a}}{p\sqrt{\hat{f}'^{\text{Eur}}\hat{f}'^{\text{Afr}}\left(1-\hat{f}'^{\text{Eur}}\right)\left(1-\hat{f}'^{\text{Afr}}\right)}}\right] \\
&\quad \times \left[\hat{f}'^{\text{Eur}}\left(1-\hat{f}'^{\text{Eur}}\right)p(1-\overline{a'}) + \hat{f}'^{\text{Afr}}\left(1-\hat{f}'^{\text{Afr}}\right)p\overline{a'}\right] \\
&= \frac{n}{n-1}\times r^2 \times \left[(1-\overline{a})^2 + \rho\overline{a}(1-\overline{a})(1-o(1)) + \overline{a}(1-\overline{a})(1+o(1))^2 + \rho\overline{a}^2(1+o(1))\right] \\
&\approx r^2(1-\overline{a}+\rho\overline{a}).
\end{aligned}$$

Because the squared denominator in Eq. (S12) is the same as in the local model,

$$\mathbb{E}_{\text{Glo}}\left[\text{Cor}(\textbf{TotPGS}, \boldsymbol{y})\right] \approx \frac{\textcircled{T}^{\text{Num}}_{\text{Glo}}}{\textcircled{T}^{\text{Den}}_{\text{Loc}}} = r(1-\overline{a}+\rho\overline{a}),$$

and so $\mathbb{E}_{\text{Glo}}\left[\text{Cor}^2(\textbf{TotPGS}, \boldsymbol{y})\right] \approx r^2(1-\overline{a}+\rho\overline{a})^2$. This proves Statement 1 of Main Text Proposition 4.2.

**Expected correlations between phenotype and ParPGS ($\mathbb{E}$[Cor(ParPGS, $\boldsymbol{y}$)]).** Under the local model, the numerator of Eq. (S11) is

$$\begin{aligned}
\textcircled{P}^{\text{Num}}_{\text{Loc}} &\approx \frac{n}{n-1}\left[\hat{f}'^{\text{Eur}}\left(1-\hat{f}'^{\text{Eur}}\right)\sum_{j=1}^{p}\left(\mathbb{E}\left[\left(\beta'^{\text{Eur}}_j\right)^2\right] + \mathbb{E}\left[\left(\beta'^{\text{Eur}}_j\right)^2\right]\right)(1-a'_j)\right] \\
&= \frac{n}{n-1}\left[2\sigma'^2_{\text{Eur}}\hat{f}'^{\text{Eur}}\left(1-\hat{f}'^{\text{Eur}}\right)\sum_{j=1}^{p}(1-a'_j)\right] \\
&\approx r^2(1-\overline{a}).
\end{aligned}$$

Observe from the simplified expressions that the squared denominator is

$$\left[\textcircled{P}^{\text{Den}}_{\text{Loc}}\right]^2 = r^2(1-\overline{a})$$

as well, so we obtain

$$\mathbb{E}_{\text{Loc}}\left[\text{Cor}(\textbf{ParPGS}, \boldsymbol{y})\right] \approx \frac{\textcircled{P}^{\text{Num}}_{\text{Loc}}}{\textcircled{P}^{\text{Den}}_{\text{Loc}}} = r\sqrt{1-\overline{a}},$$

and subsequently $\mathbb{E}_{\text{Loc}}\left[\text{Cor}^2(\textbf{ParPGS}, \boldsymbol{y})\right] \approx r^2(1-\overline{a})$. This proves Statement 3 of Main Text Proposition 4.2.

Under the global model, the numerator of Eq. (S13) is

$$\begin{aligned}
\text{(P)}^{\text{Num}}_{\text{Glo}} &\approx \frac{n}{n-1} \cdot \hat{f}'^{\text{Eur}} \left(1 - \hat{f}'^{\text{Eur}}\right) \Bigg[ (1-\overline{a}) \sum_{j=1}^{p} \left( \mathbb{E}\left[ \left(\beta_j'^{\text{Eur}}\right)^2 \right] + \mathbb{E}\left[ \left(\beta_j'^{\text{Eur}}\right)^2 \right] \right) (1 - a_j') \\
&\quad + \overline{a} \sum_{j=1}^{p} \left( \mathbb{E}\left[ \beta_j'^{\text{Eur}} \beta_j'^{\text{Afr}} \right] + \mathbb{E}\left[ \beta_j'^{\text{Eur}} \beta_j'^{\text{Afr}} \right] \right) (1 - a_j') \Bigg] \\
&= \frac{n}{n-1} \cdot \hat{f}'^{\text{Eur}} \left(1 - \hat{f}'^{\text{Eur}}\right) \left[ 2\sigma'^2_{\text{Eur}} (1-\overline{a}) p (1 - \overline{a'}) + 2\tau' \overline{a} p (1 - \overline{a'}) \right] \\
&= \frac{n}{n-1} \left[ r^2 (1-\overline{a})^2 + \overline{a}(1-\overline{a}) \rho r^2 \sqrt{\frac{\hat{f}'^{\text{Eur}} \left(1 - \hat{f}'^{\text{Eur}}\right)}{\hat{f}'^{\text{Afr}} \left(1 - \hat{f}'^{\text{Afr}}\right)}} \right] \\
&= \frac{n}{n-1} \cdot r^2 \cdot (1-\overline{a}) \left[ 1 - \overline{a} + \rho \overline{a} (1 - o(1)) \right] \\
&\approx r^2 (1-\overline{a})(1 - \overline{a} + \rho \overline{a}).
\end{aligned}$$

Because the squared denominator in Eq. (S12) is the same as in the local model,

$$\mathbb{E}_{\text{Glo}} \left[ \text{Cor}(\textbf{ParPGS}, \boldsymbol{y}) \right] \approx \frac{\text{(P)}^{\text{Num}}_{\text{Glo}}}{\text{(P)}^{\text{Den}}_{\text{Loc}}} = r\sqrt{1-\overline{a}}(1 - \overline{a} + \rho \overline{a}),$$

and so $\mathbb{E}_{\text{Glo}} \left[ \text{Cor}^2(\textbf{ParPGS}, \boldsymbol{y}) \right] \approx r^2 (1-\overline{a})(1 - \overline{a} + \rho \overline{a})^2$. This proves Statement 2 of Main Text Proposition 4.2.

#### S6 Analytical Properties of Tagging Variant Polygenic Scores

When causal variants are not known, polygenic scores are computed using tagging variants. To obtain the expressions for squared correlations, we mirror the strategy used to derive the behaviour of causal variant PGSs. First, we compute correlations for a single draw of the effect sizes, as in Supplementary Material Subsection S4. The key steps are the same, except that in all calculations involving the **TotPGS** and **ParPGS**, the "prime" expression is removed (i.e., $\beta_j'^{\text{Eur}}$ becomes $\beta_j^{\text{Eur}}$, $\beta_j'^{\text{Afr}}$ becomes $\beta_j^{\text{Afr}}$, and so on). After this is accomplished, we take expectations over the causal and tagging effect sizes, whose distributions are described in Eqs. (2) and (11) of the Main Text. Because the second step involves the approximation $\mathbb{E}[X/Y] \approx \mathbb{E}[X]/\mathbb{E}[Y]$, the final result is an approximation to the behaviour of tagging variant polygenic scores. We omit the labour of repeating the calculations, and instead list the final expressions in Supplementary Table S3, S4 and S5 (under the "Simplest Approximation" columns), while summarizing the main result in Proposition S1.

**Proposition S1** (Behaviour of Tagging Variant PGSs Under the Local and Global Models)**.** *Suppose causal variants and their effect sizes are not known, so that polygenic scores are computed using tagging variants. Let the expected squared phenotype correlations with partial polygenic score and total polygenic score under the local model (resp., global model) be denoted by* $\mathbb{E}_{Loc}[Cor^2(ParPGS, y)]$ *and* $\mathbb{E}_{Loc}[Cor^2(TotPGS, y)]$ *(resp.,* $\mathbb{E}_{Glo}[Cor^2(ParPGS, y)]$ *and* $\mathbb{E}_{Glo}[Cor^2(TotPGS, y)]$*). Then the following are true.*

1. *Under the local model,*

$$\mathbb{E}_{Loc}[Cor^2(TotPGS, y)] \approx \frac{\sum_{j=1}^{p}\left(\mathbb{E}\left[\textcircled{7L}\right] + \mathbb{E}\left[\textcircled{8L}\right] + \mathbb{E}\left[\textcircled{9L}\right] + \mathbb{E}\left[\textcircled{10L}\right]\right)}{\sqrt{\sum_{j=1}^{p}\left(\mathbb{E}\left[\textcircled{1}\right] + \mathbb{E}\left[\textcircled{2}\right] + \mathbb{E}\left[\textcircled{3}\right]\right)}} \tag{S14}$$

$$\mathbb{E}_{Loc}[Cor^2(ParPGS, y)] \approx \frac{\sum_{j=1}^{p}\left(\mathbb{E}\left[\textcircled{11L}\right] + \mathbb{E}\left[\textcircled{12L}\right] + \mathbb{E}\left[\textcircled{13L}\right] + \mathbb{E}\left[\textcircled{14L}\right]\right)}{\sqrt{\sum_{j=1}^{p}\left(\mathbb{E}\left[\textcircled{4}\right] + \mathbb{E}\left[\textcircled{5}\right] + \mathbb{E}\left[\textcircled{6}\right]\right)}} \tag{S15}$$

2. *Under the global model,*

$$\mathbb{E}_{Glo}[Cor^2(TotPGS, y)] \approx \frac{\sum_{j=1}^{p}\left(\mathbb{E}\left[\textcircled{7G}\right] + \mathbb{E}\left[\textcircled{8G}\right] + \mathbb{E}\left[\textcircled{9G}\right] + \mathbb{E}\left[\textcircled{10G}\right]\right)}{\sqrt{\sum_{j=1}^{p}\left(\mathbb{E}\left[\textcircled{1}\right] + \mathbb{E}\left[\textcircled{2}\right] + \mathbb{E}\left[\textcircled{3}\right]\right)}} \tag{S16}$$

$$\mathbb{E}_{Glo}[Cor^2(ParPGS, y)] \approx \frac{\sum_{j=1}^{p}\left(\mathbb{E}\left[\textcircled{11G}\right] + \mathbb{E}\left[\textcircled{12G}\right] + \mathbb{E}\left[\textcircled{13G}\right] + \mathbb{E}\left[\textcircled{14G}\right]\right)}{\sqrt{\sum_{j=1}^{p}\left(\mathbb{E}\left[\textcircled{4}\right] + \mathbb{E}\left[\textcircled{5}\right] + \mathbb{E}\left[\textcircled{6}\right]\right)}} \tag{S17}$$

*In Statements 1 and 2 above, the quantities* $\mathbb{E}[\textcircled{1}]$-$\mathbb{E}[\textcircled{6}]$ *are defined in Supplementary Table S3, the quantities* $\mathbb{E}\left[\textcircled{7L}\right]$-$\mathbb{E}\left[\textcircled{14L}\right]$ *are defined in Supplementary Table S4 and the quantities* $\mathbb{E}\left[\textcircled{7G}\right]$-$\mathbb{E}\left[\textcircled{14G}\right]$ *are defined in Supplementary Table S5.*

#### S7 Simulation Study Details

##### Estimating Individual Global Ancestries

To compute the global ancestry of individual $i$ in the PMBB ADM cohort, denoted $\overline{a_i}$, we take the average of local African ancestries across their tagging variants:

$$\overline{a_i} = \frac{1}{2p}\sum_{j=1}^{p}\left(a_{ij}^{(1)} + a_{ij}^{(2)}\right).$$

For **Simulation Study 1** this is fine, because we treat tagging variants as causal variants (i.e., $\lambda_j^{\text{Eur}} = \lambda_j^{\text{Afr}} = 1$, $f_j'^{\text{Eur}} = f_j^{\text{Eur}}$ and $f_j'^{\text{Afr}} = f_j^{\text{Afr}}$ for all $j$). For **Simulation Study 2**, we check that this quantity roughly matches the average of local African ancestries across causal variants, $\overline{a_i'} = \frac{1}{2p}\sum_{j=1}^{p}\left(a_{ij}'^{(1)} + a_{ij}'^{(2)}\right)$. This is indeed true: for each of the six traits, the correlation between these two quantities across all individuals is $> 0.999999$. This also justifies the assumption $\overline{a} = \overline{a'}$, which we used in Supplementary Material Subsection S5. The values of $\overline{a}$ and $\overline{a'}$ for each trait are reported in Supplementary Table S2.

##### Simulation Study 1

We elaborate on how our 50 seeds, each consisting of 1563 approximately independent markers, were prepared.

Selecting and Verifying Approximate Independence of Markers

As mentioned in Subsection 3.2 of the Main Text, approximately independent markers were selected by intersecting LD blocks for homogeneous African and European populations. These LD blocks are generated by LDetect [7], and are publicly available at their project directory, `https://bitbucket.org/nygcresearch/ldetect-data/src/master/`. The following steps were performed.

1. Download genome-wide LD block files under `EUR` and `AFR` subdirectories.
2. Intersect the LD split points.
3. Lift over from hg19 to hg38 coordinates using liftOver.

At the end of Step 3, non-overlapping LD blocks shared in common between AFR and EUR are available. Of these blocks, 1563 had non-empty intersection with our variant set. The median number of variants in a block was 134 (min = 1, max = 863), and 98% of all blocks contained at least five variants (see Supplementary Figure S18A for a visualization). To generate 50 seeds of approximately independent markers, we randomly selected one variant per block.

As an additional principled check of approximate independence of markers, flinty [8] was run on the 50 seeds. For each seed, the 1563 markers were treated as independent features and the haplotype allele dosages were summed to obtain the genotype matrix (so local ancestry calls are irrelevant here). The test of exchangeable sample and independent features (ES&IF) null was then run with metric set to the default Euclidean distance, and $p$-values were computed using large-sample approximation as described in their paper. We found that the first subgroup (called "Q1"), consisting of individuals in the lowest quantile of global African ancestry, had very small $p$-values — all were $< 10^{-10}$ — indicating violation of ES&IF (Supplementary Figure S18B). We next chose a threshold $\alpha = 0.05/(50 \times 4) = 2.5 \times 10^{-4}$, which controls the family-wise error rate at 0.05. We counted the number of subgroups within each seed for which the $p$-value was larger than $\alpha$ (i.e., fail to reject ES&IF). We found that 49 out of 50 seeds had at least two out of four subgroups that had large $p$-values (Supplementary Figure S18C), suggesting that violation of ES&IF was largely driven by the Q1 subgroup. To boost power through using larger samples, and to encourage our simulation study to mirror realistic analyses that may include linked variants (e.g., pairs of markers in long range LD), we decided to keep all four subgroups and all seeds for our simulations.

**Simulation Study 2**

As mentioned in Subsection 3.1 of the Main Text, polygenic scores were trained to decide the number of causal and tagging variants to assign for each trait. We also used the same polygenic scores to decide the range of $r^2$ values for phenotype-specific simulations. We therefore describe how polygenic scores (PGS) were constructed, and then explain how the constructed PGS were used in this simulation study.

Polygenic Score (PGS) Construction

We performed clumping and thresholding (C&T) on the PMBB EUR cohort. We first intersected variants from the processed GWAS summary statistic file (see **GWAS Summary Statistics and Causal-Tagging Variant Assignment** in Subsection 3.1) and the genotypes. Variants were then clumped using a variety of physical window sizes and index SNP $p$-value thresholds (PLINK `clump-p1` values were `0.01` or `0.0001`, `clump-kb` values were `100` or `250`). Variants in high LD were clumped (`--clump-r2 0.5` in PLINK). For each combination of physical window size and index SNP $p$-value threshold, we construct optimal C&T scores separately on median and mean traits, by selecting, among 14 $p$-value cutoffs for polygenic score SNP inclusion, the cutoff that produced the highest incremental $R^2$ over a base model that included age, sex, age$\times$sex, age$^2$ and leading principal components as covariates. To ensure robustness to population stratification confounding during this last step, we optimize C&T scores by including varying number of leading principal components (10 or 20). Altogether, for each trait we constructed 16 C&T PGSs (four combinations of physical window size and index SNP $p$-value threshold, two measurements of the trait, and two choices of PCs included in base model).

PGS Usage 1: Deciding Number of Causal and Tagging Variants

The median polygenicity ($p_{\text{median}}$) of C&T PGSs for the six traits are as follows: Standing Height ($18,000$), Weight ($16,000$), Body Mass Index ($19,000$), Triglycerides ($13,000$), Neutrophil Count ($3,700$) and Platelet Count ($14,000$). For each phenotype, we took the top $5p_{\text{median}}$ GWAS hits from the GWAS summary statistic file as candidates for causal variants. We next computed an LD table for the PMBB EUR cohort, restricting to 1000 variants within a 500 kb window and using a threshold of $\lambda^2 \geq 0.8$ for inclusion (PLINK command: `--r2 --ld-window-r2 0.8 --ld-window 1000 --ld-window-kb 500`). This produces a table of pairs of variants in high LD. We next intersected the LD table with the list of GWAS hits, where we filtered out (1) pairs of variants whereby both appear in the list of GWAS hits; (2) rows where the same variant appearing in the list of GWAS hits shows up twice. This produces a restricted table where each GWAS hit variant appears exactly one, and only one of the pairs of variants in high LD is a GWAS hit. Finally, for each "GWAS hit, non-GWAS hit" pair, we assigned the GWAS hit as the causal variant and the non-hit as the tagging variant. The final number of variant pairs for each phenotype is reported in Supplementary Table S2; they appear qualitatively consistent with previous work estimating common variant polygenicity of complex traits [9].

PGS Usage 2: Choosing Range of $r^2$ for Simulations

We regressed phenotypes against C&T PGSs on PMBB EUR cohort. We computed the coefficient of determination (model goodness-of-fit statistic), defined as

$$R^2_{\text{PGS}} = 1 - \frac{\text{sum squared residuals}}{\text{total sum of squares}},$$

and computed as `R2_PGS = summary(lm(y~PGS))$r.squared` in R. The median values of $R^2_{\text{PGS}}$ are as follows: Standing Height (0.22), Weight (0.06), Body Mass Index (0.045), Triglycerides (0.055), Neutrophil Count (0.035) and Platelet Count (0.085). The quantity $R^2_{\text{PGS}}$ is an approximately unbiased estimate of the *tagging variant* $r^2$, analogous to the causal variant $r^2$ in our model except that it is a measure of phenotypic variation explained by tagging variant genetic effects. This would result in underestimation of the causal variant $r^2$, with the downward bias being roughly equal to the mean causal-tagging squared LD, $\overline{\lambda^2_{\text{Eur}}} = \frac{1}{p}\sum_{j=1}^{p}(\lambda_j^{\text{Eur}})^2$. We thus computed $R^2_{\text{PGS}}/\overline{\lambda^2_{\text{Eur}}}$ for each trait, and assigned a grid of 10 values roughly centered on this quantity as the range of $r^2$ in our phenotype-specific simulations. The range of $r^2$ for each trait is reported in Supplementary Table S2.

#### S8 Local Ancestry Average Causal and Tagging Effects

Here, we derive the joint distribution of local ancestry average causal effects under the local and global models, as reported in Proposition 4.1 of the Main Text. Using the joint distributions, we subsequently define and derive the genome-wide correlation of local ancestry average causal effects. We also derive the joint distribution and genome-wide correlation for average tagging variant effects.

##### Proof of Proposition 4.1

As described in Subsection 2.2 of the Main Text, given phased genotype and local ancestry matrices, the average causal effect for African local ancestry at locus $j$ is computed by averaging individual effects across all individuals carrying African local ancestry at that locus. Under the local model this is just the African ancestry effect size $\beta_j'^{\text{Afr}}$. Under the global model, we obtain

$$\beta'_{\cdot j}\mid_{\text{LA=Afr}} \;=\; \frac{1}{n\left(\overline{a'^{(1)}_{\cdot j}}+\overline{a'^{(2)}_{\cdot j}}\right)}\sum_{i=1}^{n}\left(a'^{(1)}_{ij}\beta'^{\text{Glo},1}_{ij}+a'^{(2)}_{ij}\beta'^{\text{Glo},2}_{ij}\right). \tag{S18}$$

Similarly, under the local model the average causal effect for European local ancestry is $\beta_j'^{\text{Eur}}$, whereas under the global model it is

$$\beta'_{\cdot j}\mid_{\text{LA=Eur}} \;=\; \frac{1}{n\left(2-\overline{a'^{(1)}_{\cdot j}}-\overline{a'^{(2)}_{\cdot j}}\right)}\sum_{i=1}^{n}\left[\left(1-a'^{(1)}_{ij}\right)\beta'^{\text{Glo},1}_{ij}+\left(1-a'^{(2)}_{ij}\right)\beta'^{\text{Glo},2}_{ij}\right] \tag{S19}$$

We shall use Eqs. (S18) and (S19) to prove Proposition 4.1 below.

*Proof.* It is straightforward to see that under the local model, the joint distribution of average causal effects is the same as the original joint distribution of causal effects in Eq. (2) of the Main Text. We must therefore verify that under the global model the joint distribution is the multivariate Gaussian distribution described in Proposition 4.1.

Recall from Eq. (7) of the Main Text that the causal effect for individual $i$ at marker $j$ is $\beta'^{\text{Glo},1}_{ij} = \beta'^{\text{Glo},2}_{ij} = \beta'^{\text{Eur}}_{j}\left(1-\overline{a_{i\cdot}}\right)+\beta'^{\text{Afr}}_{j}\overline{a_{i\cdot}}$. We also recall from Eq. (2) of the Main Text that the distribution of $\left[\beta'^{\text{Eur}}_{j},\beta'^{\text{Afr}}_{j}\right]$ is a bivariate Gaussian parameterized by three quantities: $\sigma'^2_{\text{Eur}},\sigma'^2_{\text{Afr}}$ and $\tau'$. This shows that the local ancestry average causal effects Eqs. (S18) and (S19) are linear transformations of a multivariate Gaussian distribution, and so their joint distribution is also multivariate Gaussian.

Substitute the definition of $\beta'^{\text{Glo},h}_{ij}$ into Eq. (S18) (they are the same for $h=1,2$) to obtain $\beta'_{\cdot j}\mid_{\text{LA=Afr}} \;=\; \omega'_{1,j}\beta'^{\text{Eur}}_{j}+\omega'_{2,j}\beta'^{\text{Afr}}_{j}$, where

$$\omega'_{1,j} \;=\; \left[n\left(\overline{a'^{(1)}_{\cdot j}}+\overline{a'^{(2)}_{\cdot j}}\right)\right]^{-1}\sum_{i=1}^{n}\left(a'^{(1)}_{ij}+a'^{(2)}_{ij}\right)\left(1-\overline{a_{i\cdot}}\right) \tag{S20}$$

$$\omega'_{2,j} \;=\; \left[n\left(\overline{a'^{(1)}_{\cdot j}}+\overline{a'^{(2)}_{\cdot j}}\right)\right]^{-1}\sum_{i=1}^{n}\left(a'^{(1)}_{ij}+a'^{(2)}_{ij}\right)\overline{a_{i\cdot}}. \tag{S21}$$

This yields

$$\beta'_{\cdot j}\mid_{\text{LA=Afr}}\sim N\left(0,\sigma'^2_{\text{Eur}}\omega'^2_{1,j}+2\tau'\omega'_{1,j}\omega'_{2,j}+\sigma'^2_{\text{Afr}}\omega'^2_{2,j}\right).$$

By similar reasoning, we may obtain from Eq. (S19) the marginal distribution of $\beta'_{\cdot j}\mid_{\text{LA=Eur}}$:

$$\beta'_{\cdot j}\mid_{\text{LA=Eur}}\sim N\left(0,\sigma'^2_{\text{Eur}}\omega'^2_{3,j}+2\tau'\omega'_{3,j}\omega'_{4,j}+\sigma'^2_{\text{Afr}}\omega'^2_{4,j}\right),$$

where

$$\omega'_{3,j} = \left[n\left(2-\overline{a'^{(1)}_{\cdot j}}-\overline{a'^{(2)}_{\cdot j}}\right)\right]^{-1}\sum_{i=1}^{n}\left(2-a'^{(1)}_{ij}-a'^{(2)}_{ij}\right)(1-\overline{a_{i\cdot}}) \quad \text{(S22)}$$

$$\omega'_{4,j} = \left[n\left(2-\overline{a'^{(1)}_{\cdot j}}-\overline{a'^{(2)}_{\cdot j}}\right)\right]^{-1}\sum_{i=1}^{n}\left(2-a'^{(1)}_{ij}-a'^{(2)}_{ij}\right)\overline{a_{i\cdot}}. \quad \text{(S23)}$$

To obtain the full covariance matrix, we work out the off-diagonal term, $\mathrm{cov}\left(\beta'_{\cdot j}\mid_{\mathrm{LA=Eur}}, \beta'_{\cdot j}\mid_{\mathrm{LA=Afr}}\right)$.

By bilinearity of the covariance operator,

$$\begin{aligned}\mathrm{cov}\left(\beta'_{\cdot j}\mid_{\mathrm{LA=Eur}}, \beta'_{\cdot j}\mid_{\mathrm{LA=Afr}}\right) &= \mathrm{cov}\left(\omega'_{1,j}\beta'^{\mathrm{Eur}}_j+\omega'_{2,j}\beta'^{\mathrm{Afr}}_j, \omega'_{3,j}\beta'^{\mathrm{Eur}}_j+\omega'_{4,j}\beta'^{\mathrm{Afr}}_j\right)\\ &= \omega'_{1,j}\omega'_{3,j}\mathrm{var}\left(\beta'^{\mathrm{Eur}}_j\right)+(\omega'_{2,j}\omega'_{3,j}+\omega'_{1,j}\omega'_{4,j})\mathrm{cov}\left(\beta'^{\mathrm{Eur}}_j,\beta'^{\mathrm{Afr}}_j\right)\\ &\quad +\omega'_{2,j}\omega'_{4,j}\mathrm{var}\left(\beta'^{\mathrm{Afr}}_j\right)\\ &= \omega'_{1,j}\omega'_{3,j}\sigma'^2_{\mathrm{Eur}}+\left(\omega'_{2,j}\omega'_{3,j}+\omega'_{1,j}\omega'_{4,j}\right)\tau'+\omega'_{2,j}\omega'_{4,j}\sigma'^2_{\mathrm{Afr}}.\end{aligned}$$

We can therefore gather the terms of the covariance matrix,

$$\begin{aligned}u'_j &= \sigma'^2_{\mathrm{Eur}}\omega'^2_{1,j}+2\tau'\omega'_{1,j}\omega'_{2,j}+\sigma'^2_{\mathrm{Afr}}\omega'^2_{2,j}\\ v'_j &= \sigma'^2_{\mathrm{Eur}}\omega'^2_{3,j}+2\tau'\omega'_{3,j}\omega'_{4,j}+\sigma'^2_{\mathrm{Afr}}\omega'^2_{4,j}\\ w'_j &= \sigma'^2_{\mathrm{Eur}}\omega'_{1,j}\omega'_{3,j}+\tau'(\omega'_{2,j}\omega'_{3,j}+\omega'_{1,j}\omega'_{4,j})+\sigma'^2_{\mathrm{Afr}}\omega'_{2,j}\omega'_{4,j},\end{aligned}$$

which recovers the expressions in Proposition 4.1. □

#### Local Ancestry Average Causal Effect Correlation

We first introduce marker-specific causal effect correlations, before defining the genome-wide causal effect correlation parameter. To obtain marker-specific causal effect correlations, we consider for each marker $j$ the model correlation ($\mathrm{LAACor}'_j$), defined as the ratio of the off-diagonal covariance term to the diagonal variances in the covariance matrix. The quantity $\mathrm{LAACor}'_j$ is conceptually analogous to $r_{\mathrm{admix}}$ defined in Hou *et al.* [10], except that in our case it depends on the marker $j$ owing to differences between our model and theirs.

Under the local model, following Eq. (2) of the Main Text we obtain $\mathrm{LAACor}'^{\mathrm{Loc}}_j = \tau' \Big/ \sqrt{\sigma'^2_{\mathrm{Eur}}\sigma'^2_{\mathrm{Afr}}} = \rho$. Under the global model, however, $\mathrm{LAACor}'^{\mathrm{Glo}}_j = w'_j \Big/ \sqrt{u'_j v'_j}$.

To measure genome-wide average causal effect correlation, a natural quantity to consider is the correlation of average causal effects across all markers $j$: if the average causal effects for African and European local ancestries are the vectors $\boldsymbol{\beta}'|_{\mathrm{LA=Afr}}$ and $\boldsymbol{\beta}'|_{\mathrm{LA=Eur}}$, then this is just the empirical correlation $\mathrm{Cor}(\boldsymbol{\beta}'|_{\mathrm{LA=Eur}}, \boldsymbol{\beta}'|_{\mathrm{LA=Eur}})$. Analogous to the single marker case, we can consider a model parameter that is the expectation of this empirical quantity. This model parameter, the *expected genome-wide causal effect correlation*, is defined as

$$\overline{\mathrm{LAACor}'} = \mathbb{E}\left[\frac{\left\langle \boldsymbol{\beta}'|_{\mathrm{LA=Eur}}-\overline{\boldsymbol{\beta}'|_{\mathrm{LA=Eur}}}, \boldsymbol{\beta}'|_{\mathrm{LA=Afr}}-\overline{\boldsymbol{\beta}'|_{\mathrm{LA=Afr}}}\right\rangle}{\left\|\boldsymbol{\beta}'|_{\mathrm{LA=Eur}}-\overline{\boldsymbol{\beta}'|_{\mathrm{LA=Eur}}}\right\|_2 \cdot \left\|\boldsymbol{\beta}'|_{\mathrm{LA=Afr}}-\overline{\boldsymbol{\beta}'|_{\mathrm{LA=Afr}}}\right\|_2}\right]. \quad \text{(S24)}$$

We work out approximations for $\overline{\text{LAACor}'}$ (Eq. (S24)) under the local and global models. Under the local model, $\boldsymbol{\beta}'|_{\text{LA=Eur}} = (\beta_1'^{\text{Eur}}, \ldots, \beta_p'^{\text{Eur}})$ and $\boldsymbol{\beta}'|_{\text{LA=Afr}} = (\beta_1'^{\text{Afr}}, \ldots, \beta_p'^{\text{Afr}})$, so

$$\begin{aligned}
\overline{\text{LAACor}'_{\text{Loc}}} &= \mathbb{E}\left[\frac{\sum_{j=1}^p \left(\beta_j'^{\text{Eur}} - \overline{\beta_\cdot'^{\text{Eur}}}\right)\left(\beta_j'^{\text{Afr}} - \overline{\beta_\cdot'^{\text{Afr}}}\right)}{\sqrt{\sum_{j=1}^p \left(\beta_j'^{\text{Eur}} - \overline{\beta_\cdot'^{\text{Eur}}}\right)^2}\sqrt{\sum_{j=1}^p \left(\beta_j'^{\text{Afr}} - \overline{\beta_\cdot'^{\text{Afr}}}\right)^2}}\right] \\
&\approx \frac{\mathbb{E}\left[\sum_{j=1}^p \beta_j'^{\text{Afr}}\beta_j'^{\text{Eur}} - p \cdot \overline{\beta_\cdot'^{\text{Afr}}} \cdot \overline{\beta_\cdot'^{\text{Eur}}}\right]}{\sqrt{\mathbb{E}\left[\sum_{j=1}^p \left(\beta_j'^{\text{Eur}} - \overline{\beta_\cdot'^{\text{Eur}}}\right)^2\right]}\sqrt{\mathbb{E}\left[\sum_{j=1}^p \left(\beta_j'^{\text{Afr}} - \overline{\beta_\cdot'^{\text{Afr}}}\right)^2\right]}} \\
&= \frac{\left(\frac{p-1}{p}\right)\sum_{j=1}^p \mathbb{E}\left[\beta_j'^{\text{Afr}}\beta_j'^{\text{Eur}}\right]}{\sqrt{\left(\frac{p-1}{p}\right)\sum_{j=1}^p \mathbb{E}\left[(\beta_j'^{\text{Afr}})^2\right]}\sqrt{\left(\frac{p-1}{p}\right)\sum_{j=1}^p \mathbb{E}\left[(\beta_j'^{\text{Eur}})^2\right]}} \\
&= \frac{(p-1)\tau'}{\sqrt{(p-1)\sigma'^2_{\text{Afr}}}\sqrt{(p-1)\sigma'^2_{\text{Eur}}}} = \rho.
\end{aligned}$$

Under the global model, $\boldsymbol{\beta}'|_{\text{LA=Eur}} = (\beta'_{\cdot 1}\ |_{\text{LA=Eur}}, \ldots, \beta'_{\cdot p}\ |_{\text{LA=Eur}})$ and $\boldsymbol{\beta}'|_{\text{LA=Afr}} = (\beta'_{\cdot 1}\ |_{\text{LA=Afr}}, \ldots, \beta'_{\cdot p}\ |_{\text{LA=Afr}})$. Using the same approximating method, we obtain

$$\begin{aligned}
\overline{\text{LAACor}'_{\text{Glo}}} &\approx \frac{\left(\frac{p-1}{p}\right)\sum_{j=1}^p \mathbb{E}\left[\beta'_{\cdot j}\ |_{\text{LA=Afr}} \cdot \beta'_{\cdot j}\ |_{\text{LA=Eur}}\right]}{\sqrt{\left(\frac{p-1}{p}\right)\sum_{j=1}^p \mathbb{E}\left[(\beta'_{\cdot j}\ |_{\text{LA=Afr}})^2\right]}\sqrt{\left(\frac{p-1}{p}\right)\sum_{j=1}^p \mathbb{E}\left[(\beta'_{\cdot j}\ |_{\text{LA=Eur}})^2\right]}} \\
&= \frac{w'_1 + \ldots + w'_p}{\sqrt{u'_1 + \ldots + u'_p}\sqrt{v'_1 + \ldots + v'_p}}.
\end{aligned}$$

The key results are summarized in **Box C** for easy reference.

**Box C: Expected Genome-wide Causal Effect Correlation**

The genome-wide causal effect correlation by local ancestry is the correlation of average causal effect vectors. When the causal effect vectors themselves are realizations of (i.e., drawn from) a probability distribution, this quantity can be viewed as a random variable. Its expectation, $\overline{\text{LAACor}'}$ (Eq. (S24)), is the *expected genome-wide causal effect correlation.* Under the local model, the expected genome-wide causal effect correlation is

$$\overline{\text{LAACor}'_{\text{Loc}}} \approx \rho. \tag{S25}$$

Under the global model, the expected genome-wide causal effect correlation is

$$\overline{\text{LAACor}'_{\text{Glo}}} \approx \frac{w'_1 + \ldots + w'_p}{\sqrt{u'_1 + \ldots + u'_p}\sqrt{v'_1 + \ldots + v'_p}}. \tag{S26}$$

**Local Ancestry Average Tagging Effect Distribution and Correlation**

Similar to average causal effects, average tagging effects are defined by averaging the tagging effect sizes across all individuals of one local ancestry. Because the results here are not crucial to the core findings of our work, we will outline the main ideas behind the derivation before stating the key results in **Box D**.

Under the local model, the joint distribution of average tagging effects is the same as the original joint distribution of tagging effects in Eq. (11) of the Main Text. To obtain the joint distribution under the global model, we may repeat the strategy used for obtaining the joint distribution of average causal effects (Proposition 4.1). It sounds complicated, but the mathematical steps are just slight modifications of the arguments presented in the proof above, where each occurrence of $\sigma'^2_{\text{Eur}}$ is replaced with $\sigma'^2_{\text{Eur}}(\theta_j^{\text{Eur}})^2$, each occurrence of $\sigma'^2_{\text{Afr}}$ is replaced with $\sigma'^2_{\text{Afr}}(\theta_j^{\text{Afr}})^2$ and so on. By replacing each quantity in the average causal effect argument with the appropriate quantity for the average tagging effect, we can obtain the joint distribution. Once the joint distribution is obtained, we may then analogously define the genome-wide tagging effect correlation (similar to Eq. (S24)) and repeat the steps for deriving their approximations under the local and global models.

**Box D: Joint Distribution of Average Tagging Effects, and Expected Genome-wide Tagging Effect Correlation**

Under the local model, the joint distribution of average tagging effects is the same as the original joint distribution of tagging effects in the base model, Eq. (11) of the Main Text. Under the global model, the joint distribution of average tagging effects is

$$\begin{bmatrix}\beta_{\cdot j}\ |_{\text{LA=Afr}}\\ \beta_{\cdot j}\ |_{\text{LA=Eur}}\end{bmatrix} \sim N\left(\begin{bmatrix}0\\0\end{bmatrix}, \begin{bmatrix}u_j & w_j\\ w_j & v_j\end{bmatrix}\right),$$

where

$$\begin{aligned}
u_j &= \sigma'^2_{\text{Eur}}(\theta_j^{\text{Eur}})^2\omega_{1,j}^2 + 2\tau'\theta_j^{\text{Eur}}\theta_j^{\text{Afr}}\omega_{1,j}\omega_{2,j} + \sigma'^2_{\text{Afr}}(\theta_j^{\text{Afr}})^2\omega_{2,j}^2,\\
v_j &= \sigma'^2_{\text{Eur}}(\theta_j^{\text{Eur}})^2\omega_{3,j}^2 + 2\tau'\theta_j^{\text{Eur}}\theta_j^{\text{Afr}}\omega_{3,j}\omega_{4,j} + \sigma'^2_{\text{Afr}}(\theta_j^{\text{Afr}})^2\omega_{4,j}^2,\\
w_j &= \omega_{1,j}\omega_{3,j}\sigma'^2_{\text{Eur}}(\theta_j^{\text{Eur}})^2 + (\omega_{2,j}\omega_{3,j} + \omega_{1,j}\omega_{4,j})\tau'\theta_j^{\text{Eur}}\theta_j^{\text{Afr}} + \omega_{2,j}\omega_{4,j}\sigma'^2_{\text{Afr}}(\theta_j^{\text{Afr}})^2,
\end{aligned}$$

with $\left(\sigma'^2_{\text{Eur}}, \sigma'^2_{\text{Afr}}, \tau'\right)$ defined in Eq. (2) of the Main Text; $\theta_j^{\text{Eur}}$ and $\theta_j^{\text{Afr}}$ defined in Subsection 2.3 of the Main Text; and

$$\begin{aligned}
\omega_{1,j} &= \left[n\left(\overline{a_{\cdot j}^{(1)}} + \overline{a_{\cdot j}^{(2)}}\right)\right]^{-1}\sum_{i=1}^{n}\left(a_{ij}^{(1)} + a_{ij}^{(2)}\right)(1-\overline{a_{i\cdot}})\\
\omega_{2,j} &= \left[n\left(\overline{a_{\cdot j}^{(1)}} + \overline{a_{\cdot j}^{(2)}}\right)\right]^{-1}\sum_{i=1}^{n}\left(a_{ij}^{(1)} + a_{ij}^{(2)}\right)\overline{a_{i\cdot}}\\
\omega_{3,j} &= \left[n\left(2-\overline{a_{\cdot j}^{(1)}} - \overline{a_{\cdot j}^{(2)}}\right)\right]^{-1}\sum_{i=1}^{n}\left(2-a_{ij}^{(1)} - a_{ij}^{(2)}\right)(1-\overline{a_{i\cdot}})\\
\omega_{4,j} &= \left[n\left(2-\overline{a_{\cdot j}^{(1)}} - \overline{a_{\cdot j}^{(2)}}\right)\right]^{-1}\sum_{i=1}^{n}\left(2-a_{ij}^{(1)} - a_{ij}^{(2)}\right)\overline{a_{i\cdot}}.
\end{aligned}$$

Moreover, this distribution is independent across markers $j$.

The genome-wide tagging effect correlation by local ancestry is the correlation of average tagging effect vectors. When the tagging effect vectors themselves are realizations of a probability distribution, this quantity can be viewed as a random variable. Its expectation, $\overline{\text{LAACor}}$, is the expected genome-wide tagging effect correlation. Under the local model, the expected genome-wide tagging effect correlation is

$$\overline{\text{LAACor}_{\text{Loc}}} \approx \frac{\rho\left(\theta_1^{\text{Eur}}\theta_1^{\text{Afr}} + \ldots + \theta_p^{\text{Eur}}\theta_p^{\text{Afr}}\right)}{\sqrt{(\theta_1^{\text{Eur}})^2 + \ldots + (\theta_p^{\text{Eur}})^2}\sqrt{(\theta_1^{\text{Afr}})^2 + \ldots + (\theta_p^{\text{Afr}})^2}}. \tag{S27}$$

Under the global model, the expected genome-wide tagging effect correlation is

$$\overline{\text{LAACor}_{\text{Glo}}} \approx \frac{w_1 + \ldots + w_p}{\sqrt{u_1 + \ldots + u_p}\sqrt{v_1 + \ldots + v_p}}. \tag{S28}$$

**Simulation Study**

To verify that the correlation approximations in **Box C** and **Box D** are reasonably accurate, we simulated causal and tagging effects using Eqs. (2) and (11) for the six phenotypes for which we assigned putative causal and tagging variants. We considered a wide range of parameters:

- 25 choices of $\rho$ (15 values equally spaced in $[0.2, 0.9]$, and $\{0.91, \ldots, 0.99, 1\}$)
- 10 choices of $r^2$, chosen in a phenotype-specific manner to include empirically observed $r^2$, as estimated from fitting clumping and thresholding polygenic scores in a European-specific cohort

For each pair $(\rho, r^2)$, we computed genome-wide average effect correlations $\overline{\text{LAACor}'}$ and $\overline{\text{LAACor}}$ under the local model (Eqs. (S25) and (S27)) and the global model (Eqs. (S26) and (S28)). Subsequently, we drew 100 pairs of effect size vectors under each model and computed empirical correlations of the average effect vectors. We calculated averages of the latter two quantities and constructed approximate 95% confidence intervals for them, before comparing them against $\overline{\text{LAACor}'}$ and $\overline{\text{LAACor}}$.

Across all six phenotypes, we find good agreement between the model parameters ($\overline{\text{LAACor}'}$ and $\overline{\text{LAACor}}$) and the empirical correlations of local ancestry average effect vectors (Supplementary Figures S19 and S20).

#### Penn Medicine Biobank Team and Contributions

##### Leadership

- Daniel J. Rader, M.D.; Marylyn D. Ritchie, Ph.D.

Contribution: All authors contributed to securing funding, study design and oversight. All authors reviewed the final version of the manuscript.

##### Patient Recruitment and Regulatory Oversight

- JoEllen Weaver; Nawar Naseer, Ph.D., M.P.H.; Giorgio Sirugo, M.D., Ph.D.; Afiya Poindexter; Yi-An Ko, Ph.D.; Kyle P. Nerz

Contributions: JW manages patient recruitment and regulatory oversight of study. NN manages participant engagement, assists with regulatory oversight, and researcher access. GS assists with researcher access. AP, YK, KPN perform recruitment and enrollment of study participants.

##### Lab Operations

- JoEllen Weaver; Meghan Livingstone; Fred Vadivieso; Stephanie DerOhannessian; Teo Tran; Julia Stephanowski; Salma Santos; Ned Haubein, Ph.D.; Joseph Dunn

Contribution: JW, ML, FV, SD conduct oversight of lab operations. ML, FV, AK, SD, TT, JS, SS perform sample processing. NH, JD are responsible for sample tracking and the laboratory information management system.

##### Clinical Informatics

- Anurag Verma, Ph.D.; Colleen Morse Kripke, M.S. DPT, MSA; Marjorie Risman, M.S.; Renae Judy, B.S.; Colin Wollack, M.S.

Contribution: All authors contributed to the development and validation of clinical phenotypes used to identify study subjects and (when applicable) controls.

##### Genome Informatics

- Anurag Verma Ph.D.; Shefali S. Verma, Ph.D.; Scott Damrauer, M.D.; Yuki Bradford, M.S.; Scott Dudek, M.S.; Theodore Drivas, M.D., Ph.D.

Contribution: AV, SSV, and SD are responsible for the analysis, design, and infrastructure needed to quality control genotype and exome data. YB performs the analysis. TD and AV provides variant and gene annotations and their functional interpretation of variants.

### Supplementary Figures and Tables

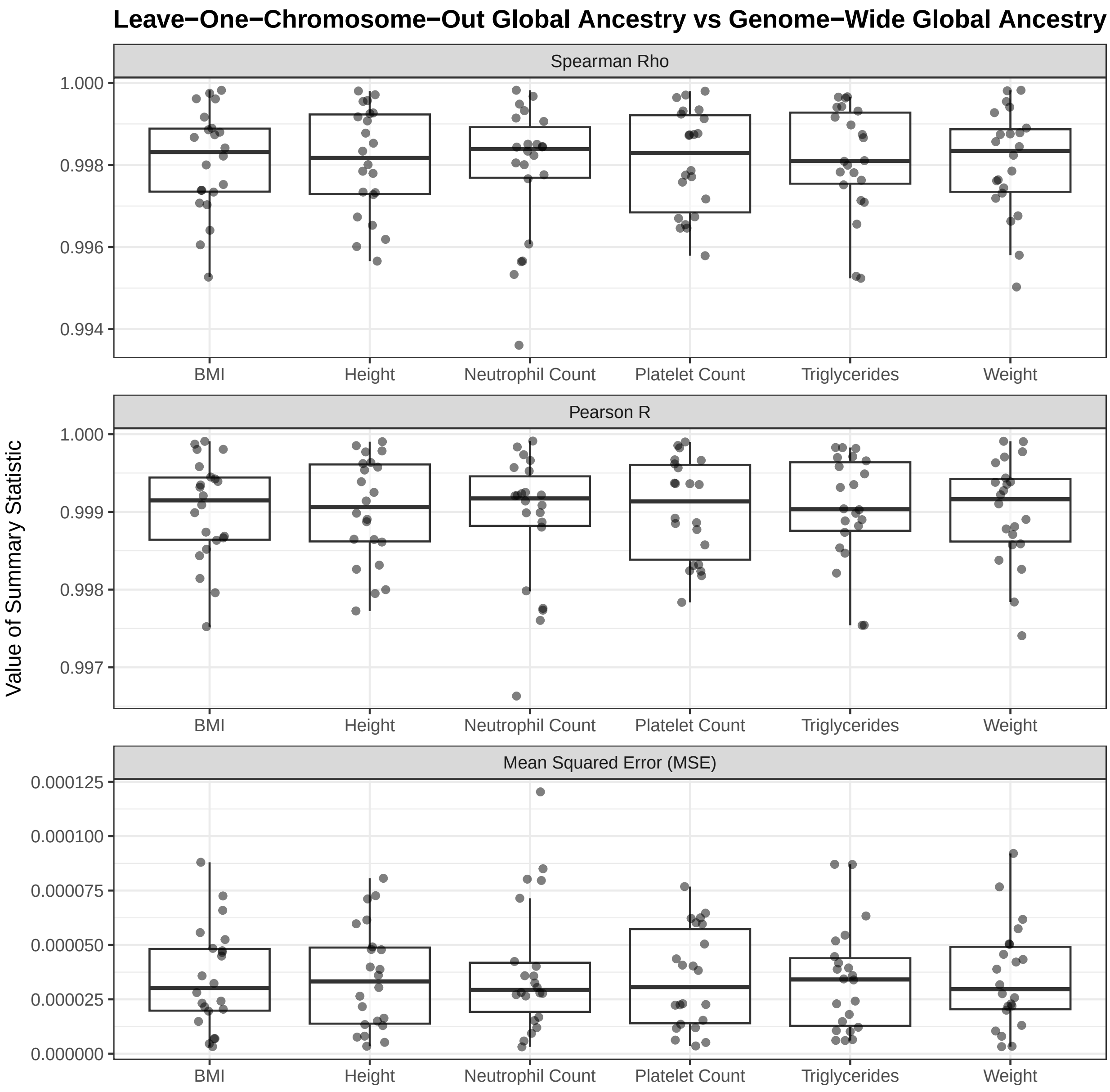

Figure S1: Comparison of Leave-One-Chromosome-Out calculation of individual global ancestries and the genome-wide calculation used in our work. For each autosome $j \in \{1, \ldots, 22\}$, all variants from Chr $j$ were excluded in the calculation of global African ancestry as mean proportion of local African ancestries. The resulting vector of individual global ancestries, $\overline{a_{i\cdot}^{-j}}$, is subsequently compared against the original vector of individual genome-wide global ancestries, $\overline{a_{i\cdot}}$, using three metrics: Spearman and Pearson correlations, as well as the mean squared error (MSE). Across the six traits, the median Spearman Rho $\geq 0.998$, the median Pearson R $\geq 0.999$, and the median MSE $\leq 0.00013$.

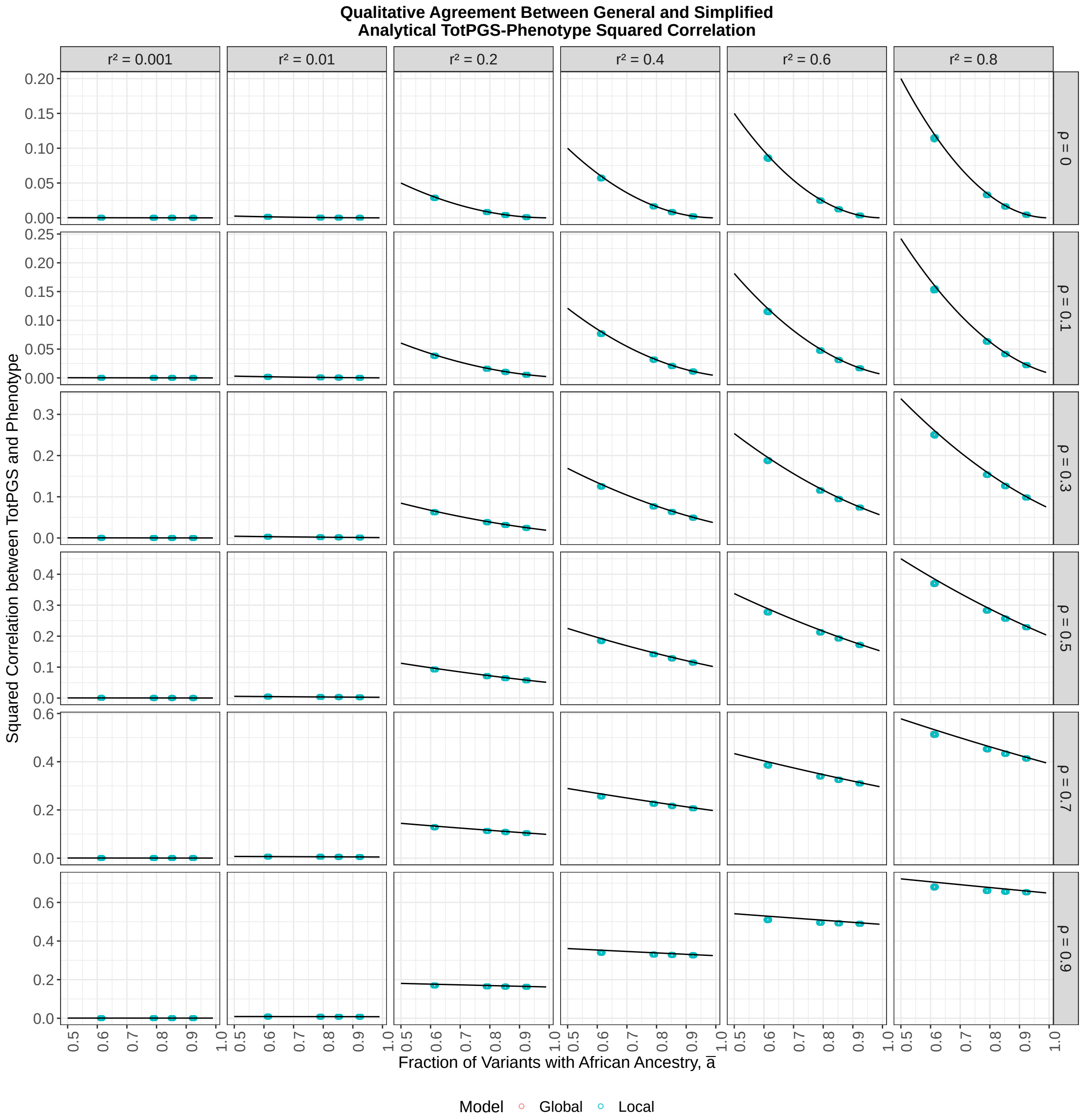

Figure S2: Agreement between analytical predictions of total polygenic score performance (1) under additional assumptions about allele frequencies and ancestry assignments as described in Supplementary Material Section S5, and (2) under no additional assumptions. Solid lines depict the predicted behaviour under additional assumptions, whereas each point depicts the predicted behaviour for a particular seed's distribution of allele frequencies and ancestry assignment as well as quantile. Each plot within the panel has a single line corresponding to the first equation of Theorem 4.2 in the Main Text; and $50 \times 4 \times 2 = 400$ points corresponding to the number of seeds simulated (50), number of quantiles used (4) and whether the local or the global model was assumed when working out the analytical formulae (2).

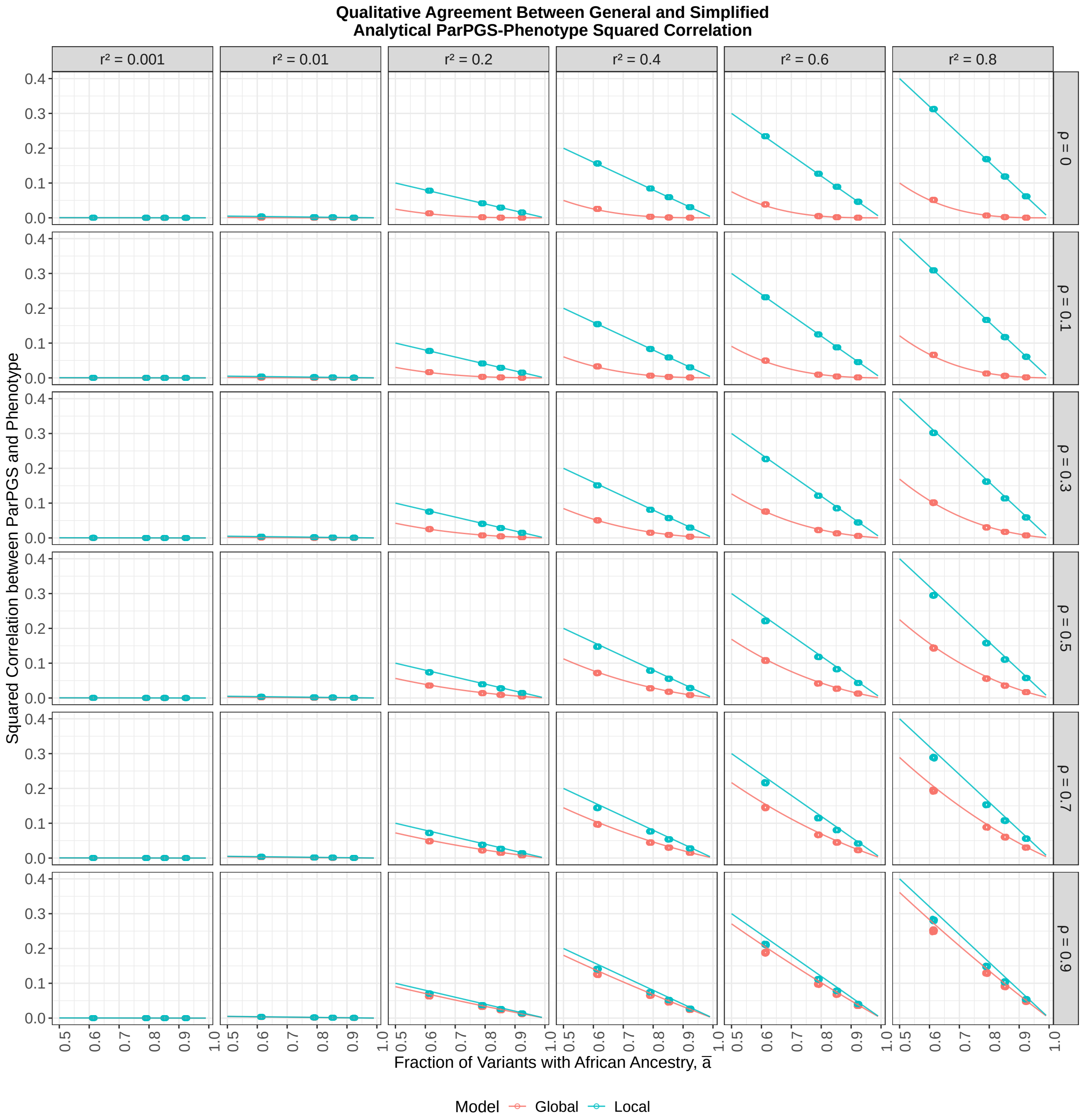

Figure S3: Agreement between analytical predictions of partial polygenic score performance (1) under additional assumptions about allele frequencies and ancestry assignments as described in Supplementary Material Section S5, and (2) under no additional assumptions. Solid lines depict the predicted behaviour under additional assumptions, whereas each point depicts the predicted behaviour for a particular seed's distribution of allele frequencies and ancestry assignment as well as quantile. Each plot within the panel has two lines corresponding to the second and third equations of Theorem 4.2 in the Main Text; and $50 \times 4 \times 2 = 400$ points corresponding to the number of seeds simulated (50), number of quantiles used (4) and whether the local or the global model was assumed when working out the analytical formulae (2).

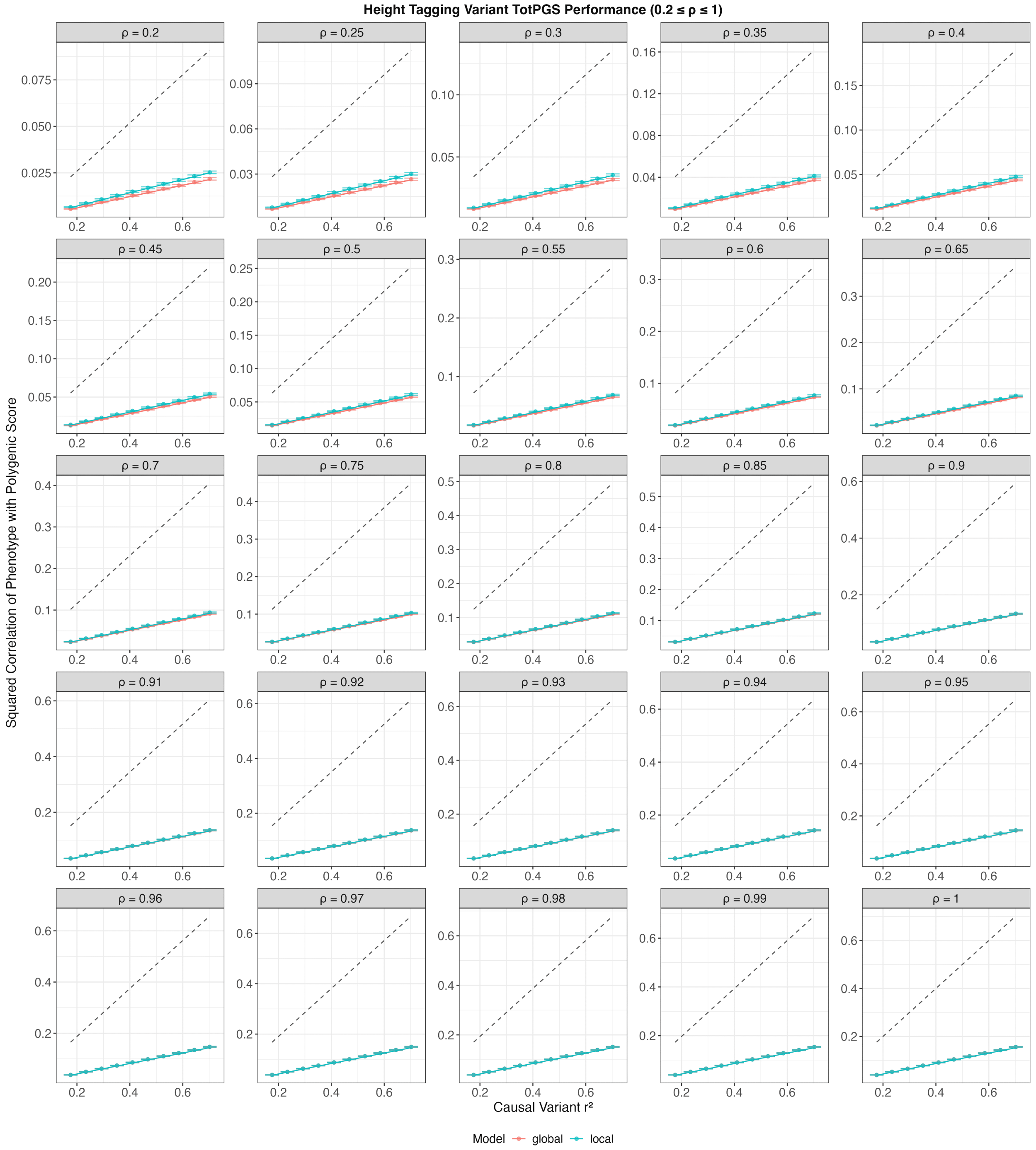

Figure S4: Standard PGS (TotPGS) performance when polygenic scores are computed using height-specific tagging variants and simulated tagging effect sizes under Eq. (11) of Main Text. Solid coloured curves are approximate expectations of the performance under the local and the global model. Black dashed curves are analytical quantities (see Proposition 4.2 of Main Text) describing the approximate performance of TotPGS if height-specific causal variants and their effects are used instead (note $\overline{a} \approx 0.8$).

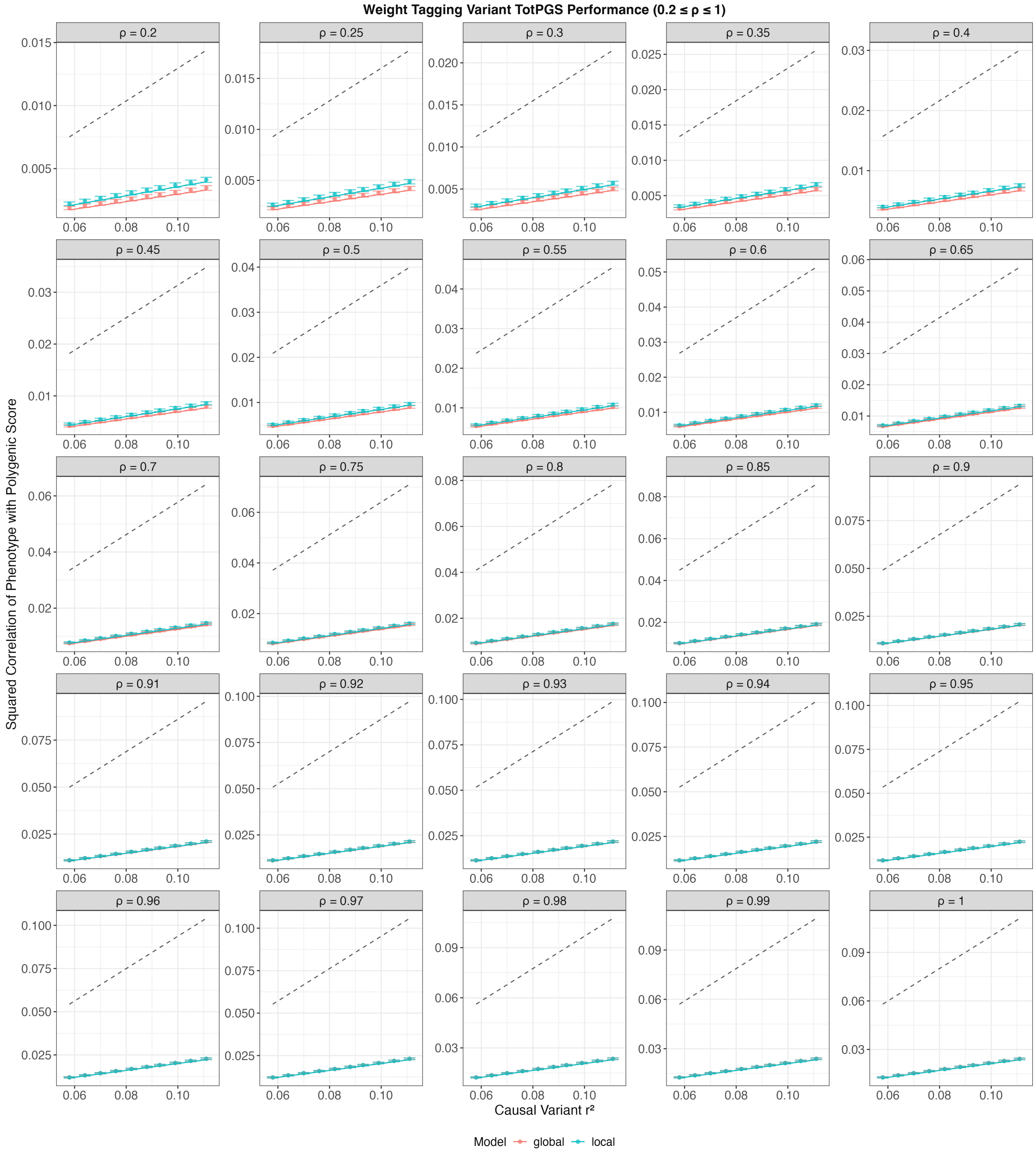

Figure S5: Standard PGS (TotPGS) performance when polygenic scores are computed using weight-specific tagging variants and simulated tagging effect sizes under Eq. (11) of Main Text. Solid coloured curves are approximate expectations of the performance under the local and the global model. Black dashed curves are analytical quantities (see Proposition 4.2 of Main Text) describing the approximate performance of TotPGS if weight-specific causal variants and their effects are used instead (note $\overline{a} \approx 0.8$).

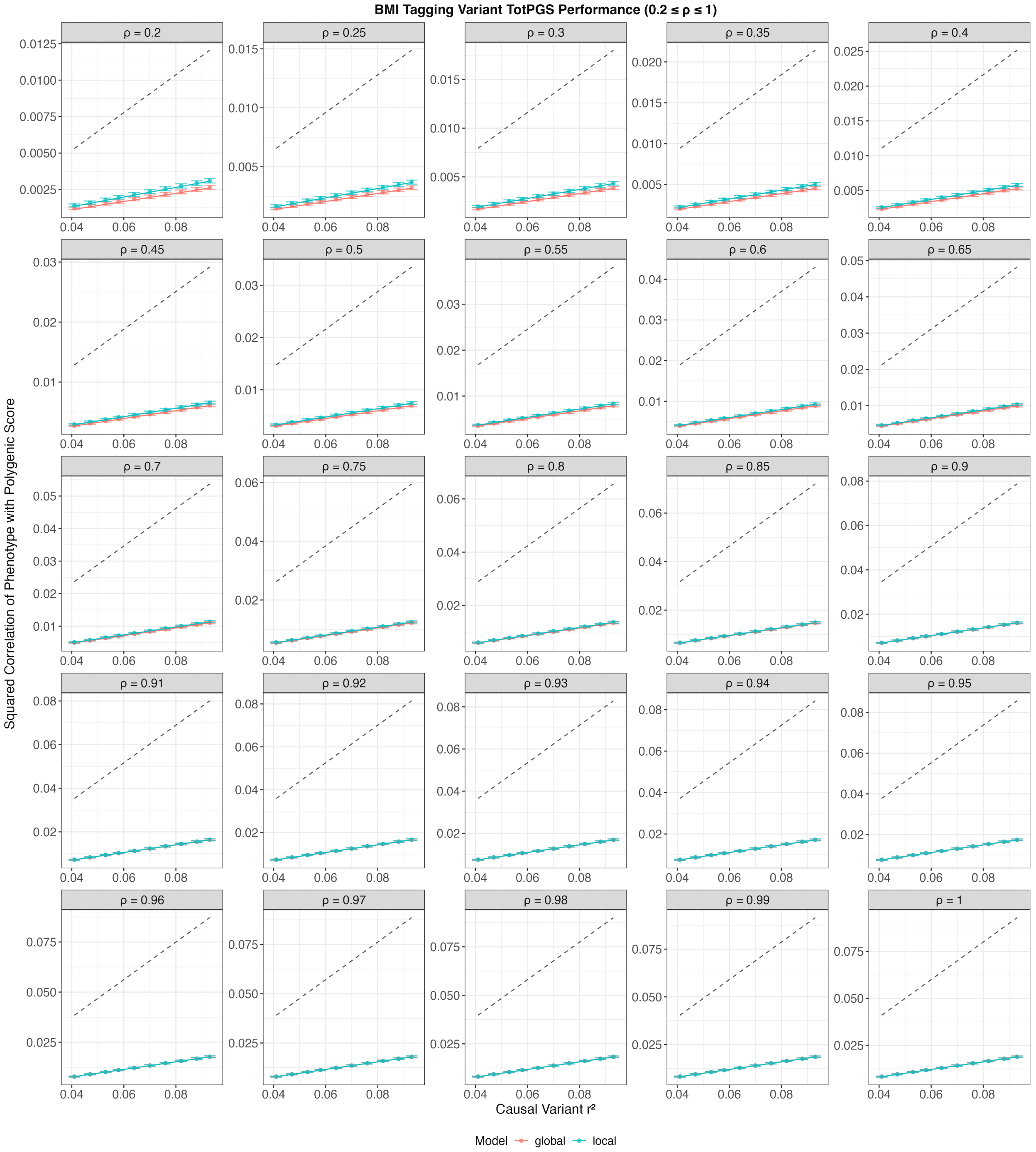

Figure S6: Standard PGS (TotPGS) performance when polygenic scores are computed using BMI-specific tagging variants and simulated tagging effect sizes under Eq. (11) of Main Text. Solid coloured curves are approximate expectations of the performance under the local and the global model. Black dashed curves are analytical quantities (see Proposition 4.2 of Main Text) describing the approximate performance of TotPGS if BMI-specific causal variants and their effects are used instead (note $\overline{a} \approx 0.8$).

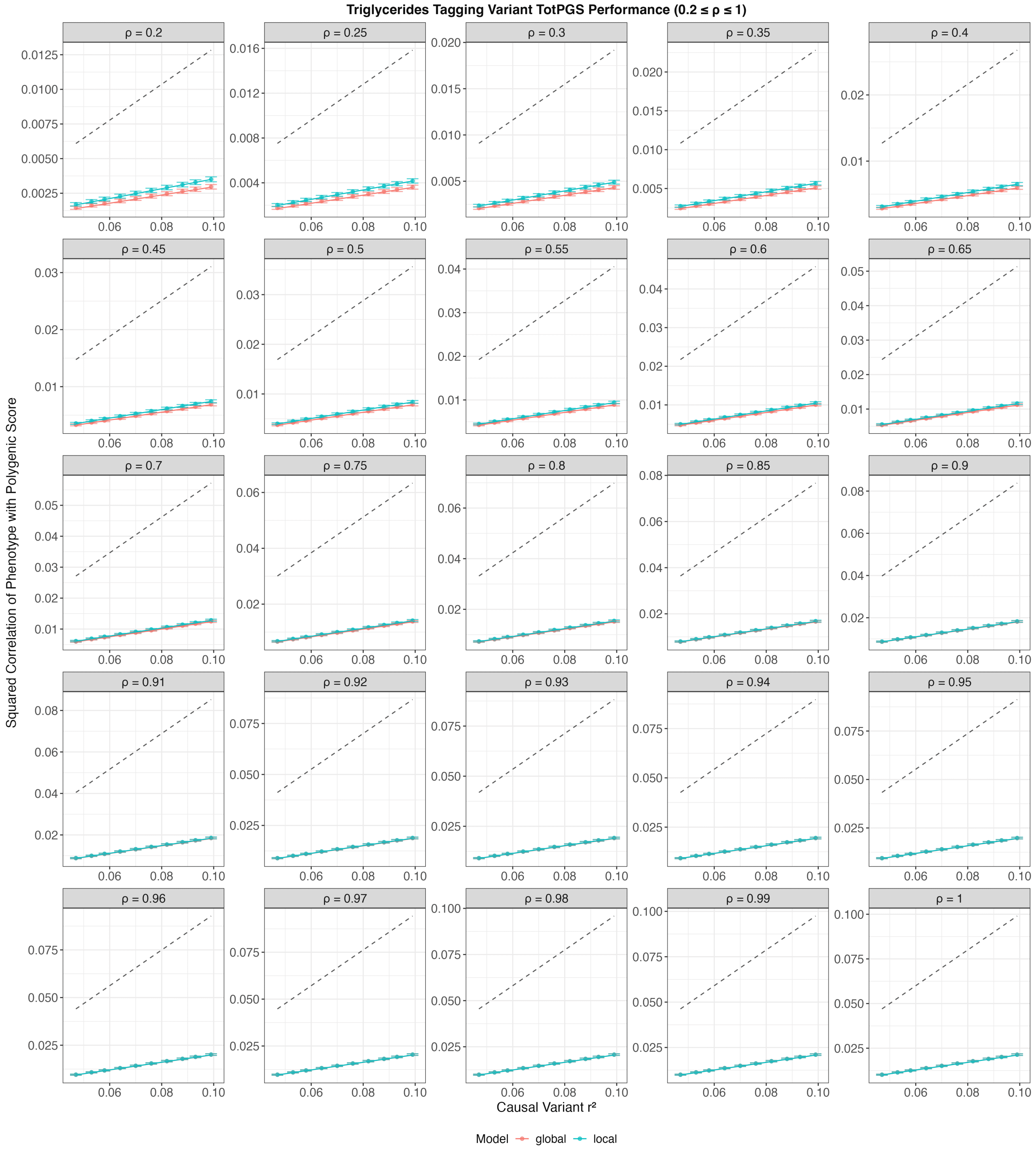

Figure S7: Standard PGS (TotPGS) performance when polygenic scores are computed using triglycerides-specific tagging variants and simulated tagging effect sizes under Eq. (11) of Main Text. Solid coloured curves are approximate expectations of the performance under the local and the global model. Black dashed curves are analytical quantities (see Proposition 4.2 of Main Text) describing the approximate performance of TotPGS if triglycerides-specific causal variants and their effects are used instead (note $\overline{a} \approx 0.8$).

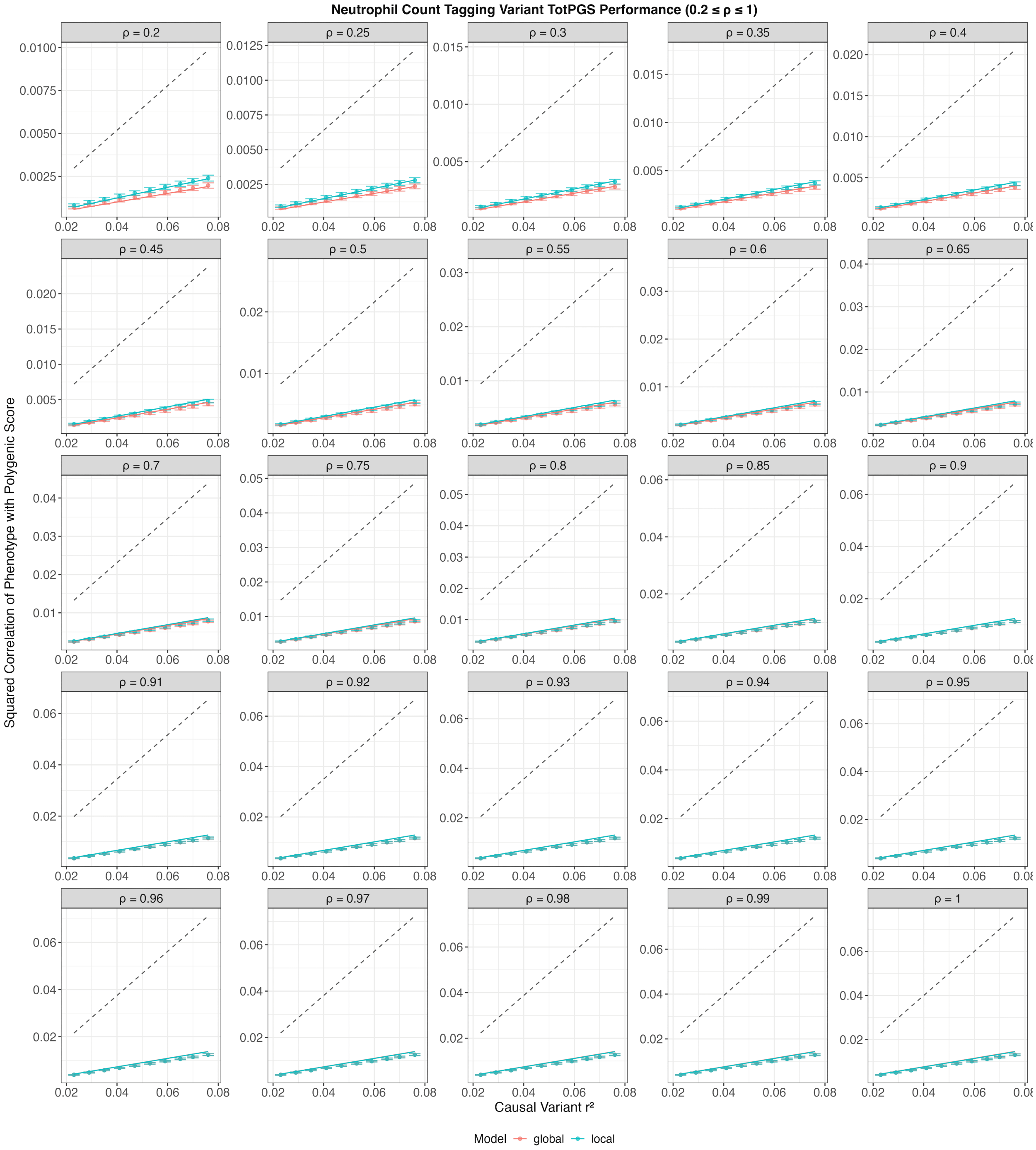

Figure S8: Standard PGS (TotPGS) performance when polygenic scores are computed using neutrophil count-specific tagging variants and simulated tagging effect sizes under Eq. (11) of Main Text. Solid coloured curves are approximate expectations of the performance under the local and the global model. Black dashed curves are analytical quantities (see Proposition 4.2 of Main Text) describing the approximate performance of TotPGS if neutrophil count-specific causal variants and their effects are used instead (note $\overline{a} \approx 0.8$).

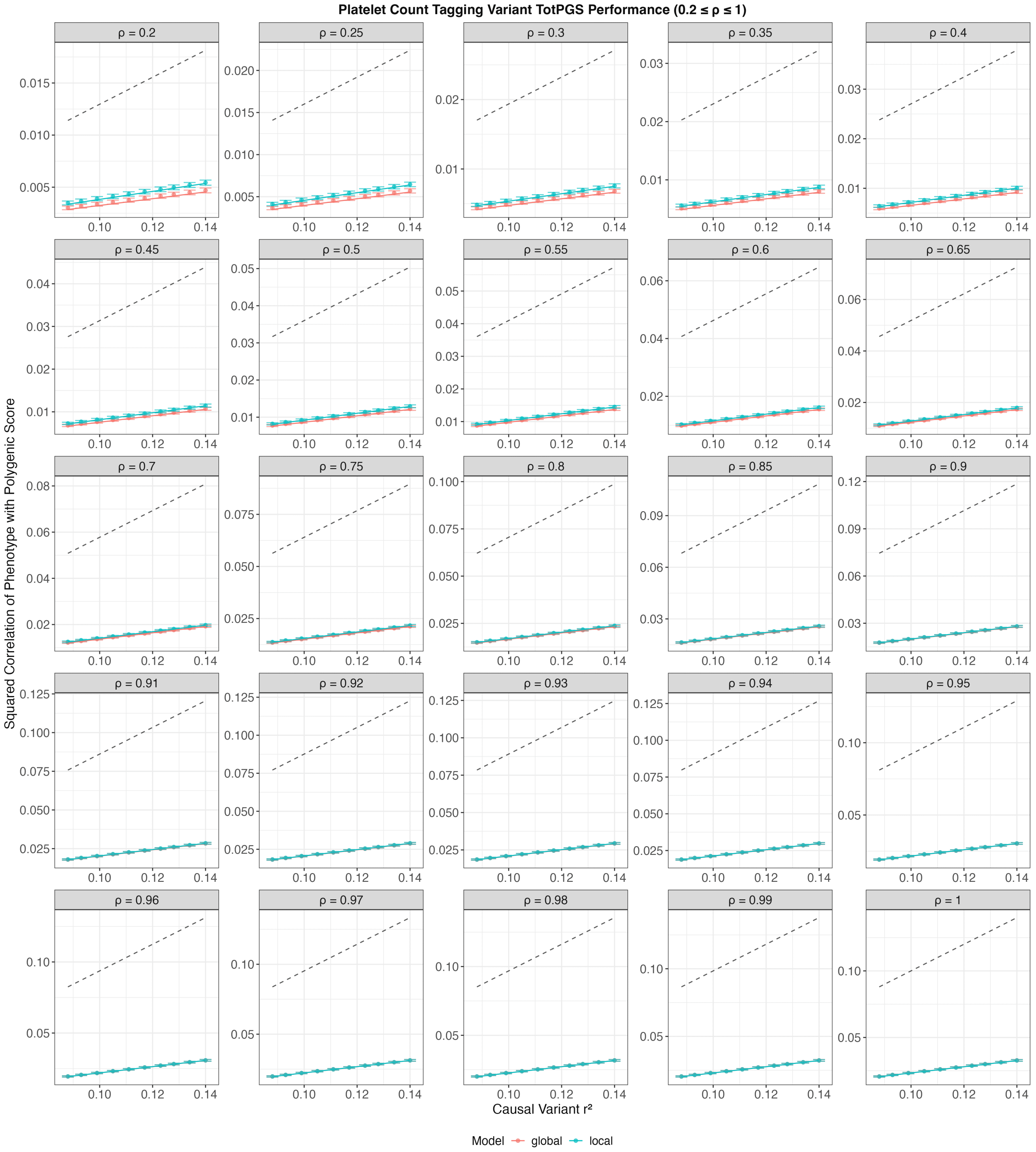

Figure S9: Standard PGS (TotPGS) performance when polygenic scores are computed using platelet count-specific tagging variants and simulated tagging effect sizes under Eq. (11) of Main Text. Solid coloured curves are approximate expectations of the performance under the local and the global model. Black dashed curves are analytical quantities (see Proposition 4.2 of Main Text) describing the approximate performance of TotPGS if platelet count-specific causal variants and their effects are used instead (note $\overline{a} \approx 0.8$).

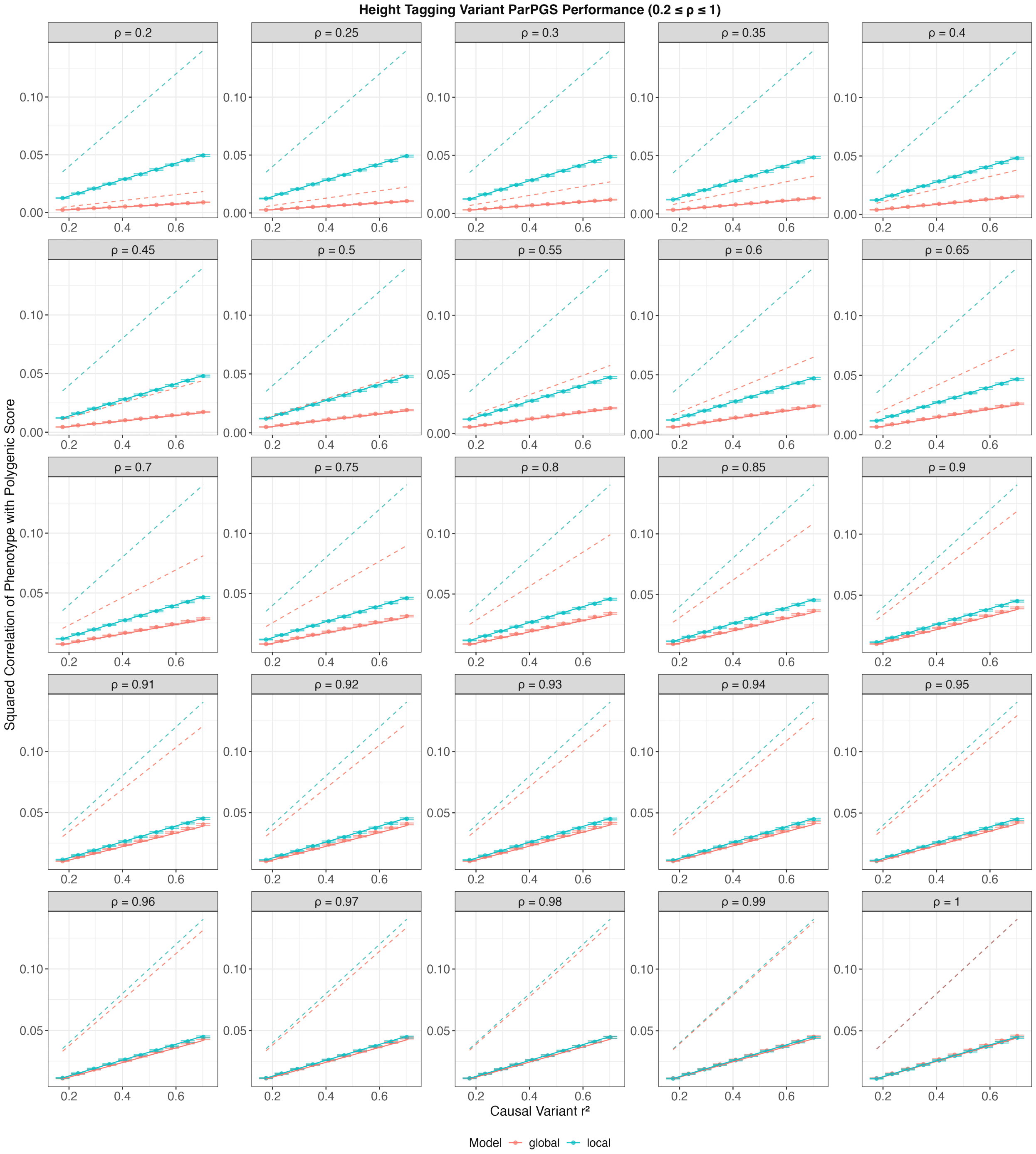

Figure S10: Partial PGS (ParPGS) performance when polygenic scores are computed using height-specific tagging variants and simulated tagging effect sizes under Eq. (11) of Main Text. Solid coloured curves are approximate expectations of the performance under the local and the global model. Dashed coloured curves are analytical quantities (see Proposition 4.2 of Main Text) describing model-specific approximate performances of ParPGS if height-specific causal variants and their effects are used instead (note $\bar{a} \approx 0.8$).

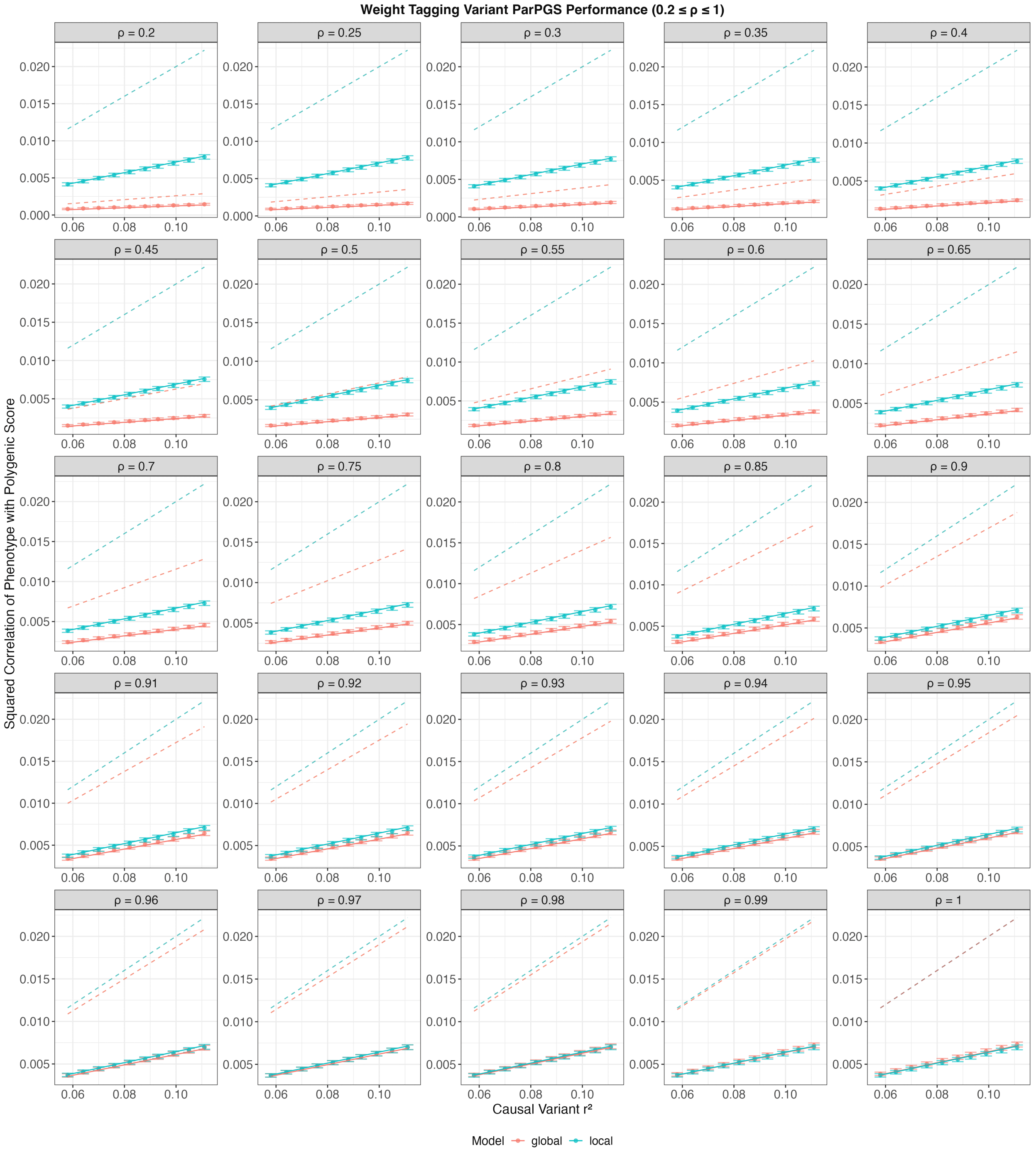

Figure S11: Partial PGS (ParPGS) performance when polygenic scores are computed using weight-specific tagging variants and simulated tagging effect sizes under Eq. (11) of Main Text. Solid coloured curves are approximate expectations of the performance under the local and the global model. Dashed coloured curves are analytical quantities (see Proposition 4.2 of Main Text) describing model-specific approximate performances of ParPGS if weight-specific causal variants and their effects are used instead (note $\overline{a} \approx 0.8$).

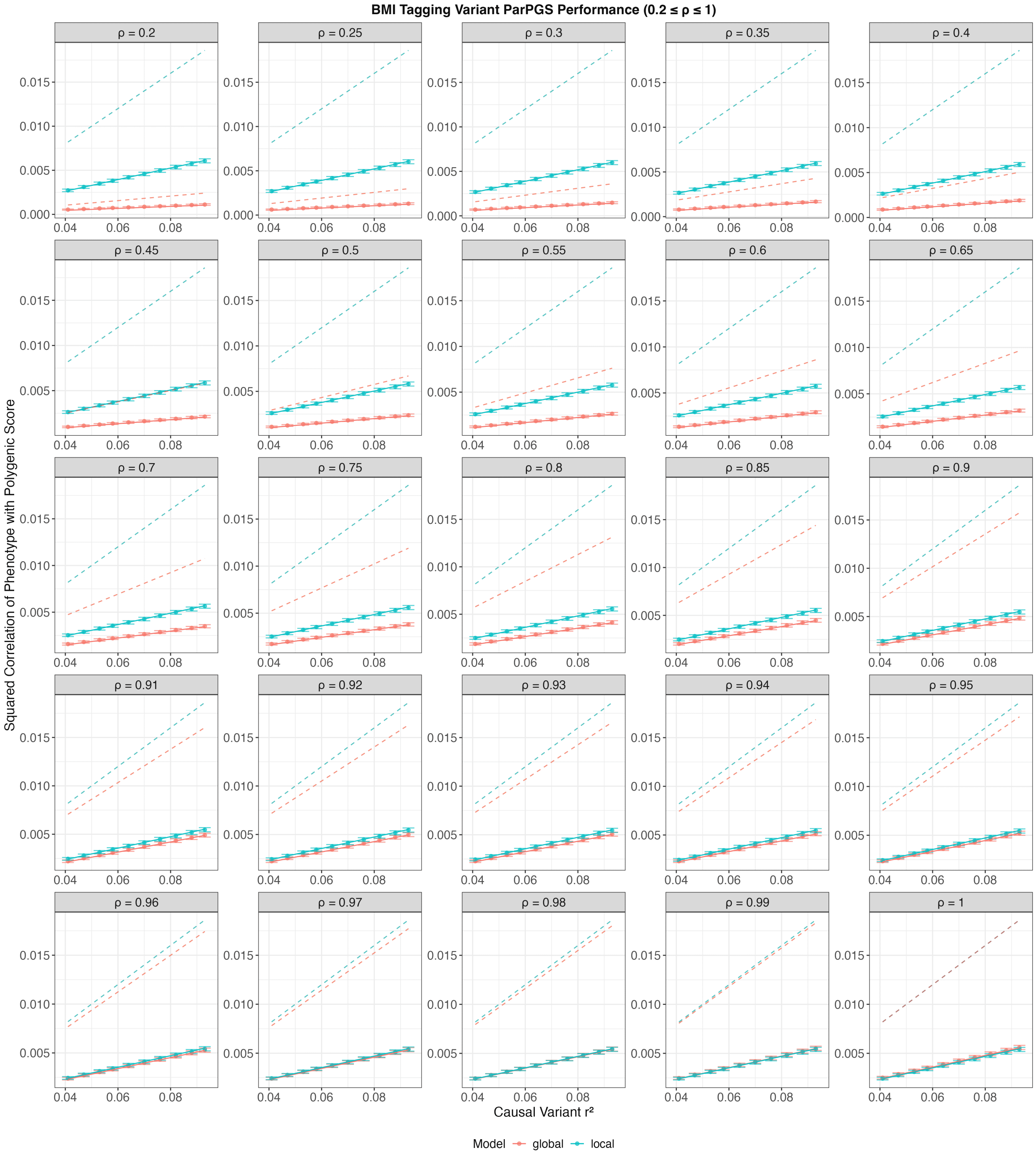

Figure S12: Partial PGS (ParPGS) performance when polygenic scores are computed using BMI-specific tagging variants and simulated tagging effect sizes under Eq. (11) of Main Text. Solid coloured curves are approximate expectations of the performance under the local and the global model. Dashed coloured curves are analytical quantities (see Proposition 4.2 of Main Text) describing model-specific approximate performances of ParPGS if BMI-specific causal variants and their effects are used instead (note $\overline{a} \approx 0.8$).

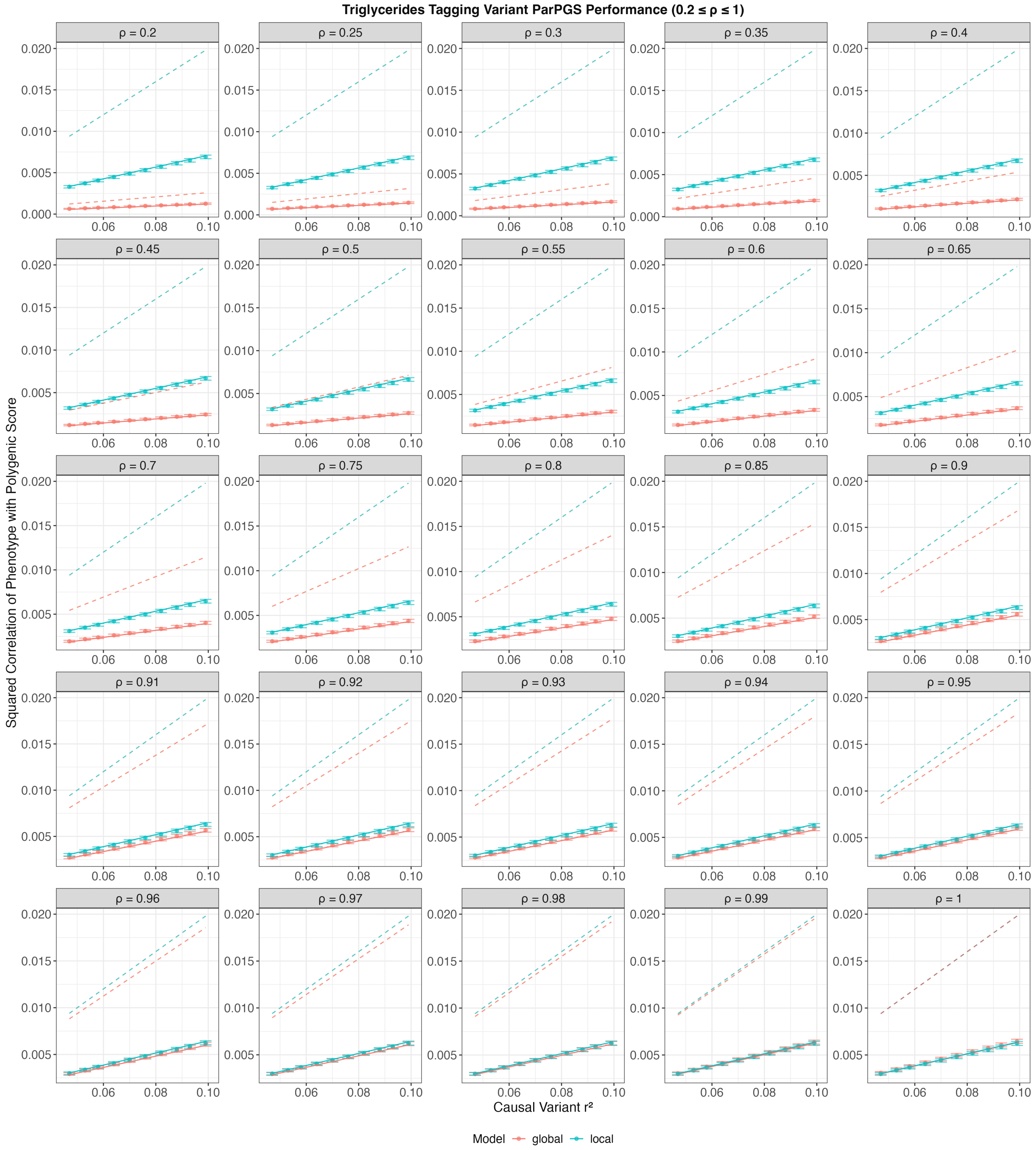

Figure S13: Partial PGS (ParPGS) performance when polygenic scores are computed using triglycerides-specific tagging variants and simulated tagging effect sizes under Eq. (11) of Main Text. Solid coloured curves are approximate expectations of the performance under the local and the global model. Dashed coloured curves are analytical quantities (see Proposition 4.2 of Main Text) describing model-specific approximate performances of ParPGS if triglycerides-specific causal variants and their effects are used instead (note $\bar{a} \approx 0.8$).

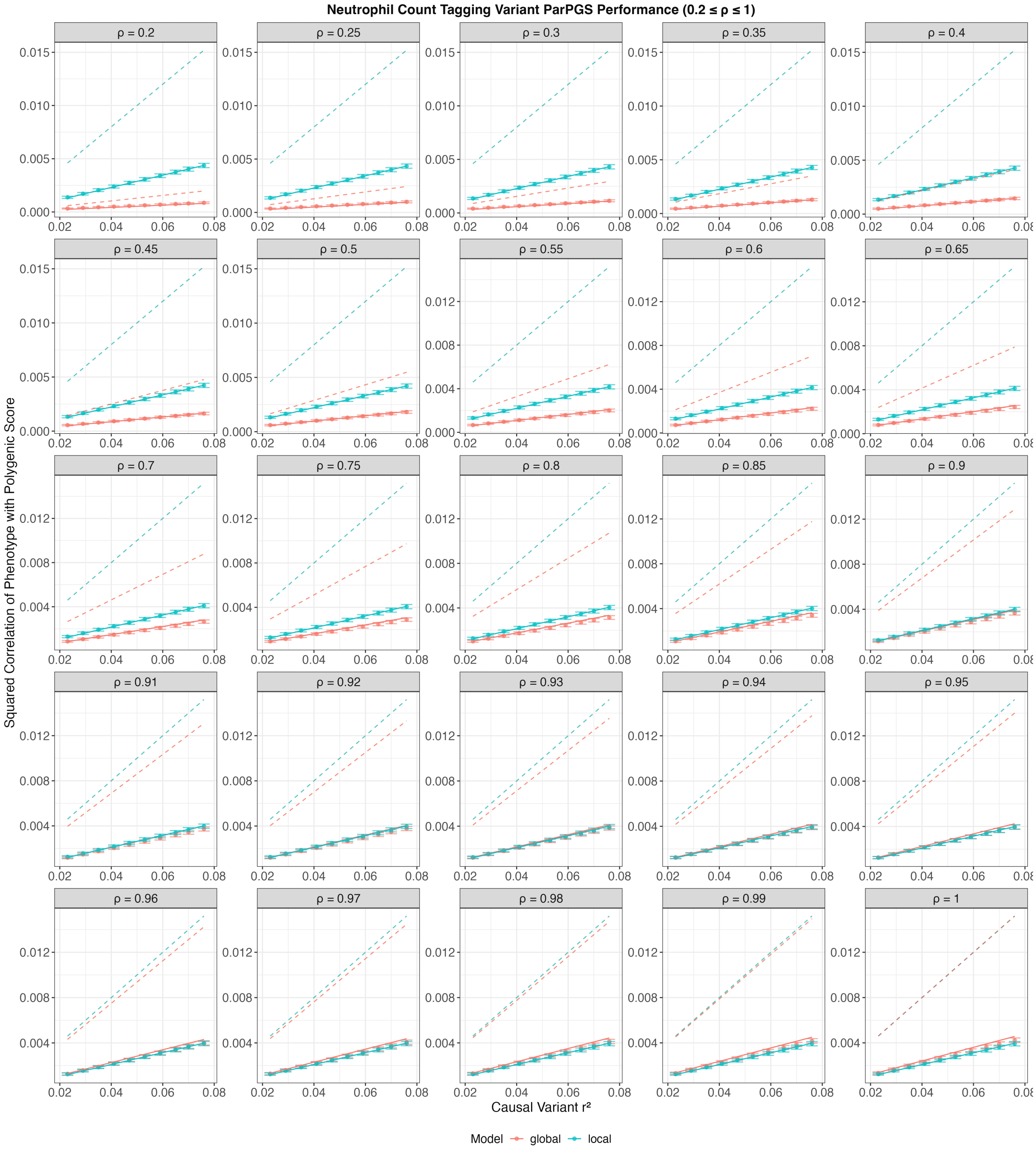

Figure S14: Partial PGS (ParPGS) performance when polygenic scores are computed using neutrophil count-specific tagging variants and simulated tagging effect sizes under Eq. (11) of Main Text. Solid coloured curves are approximate expectations of the performance under the local and the global model. Dashed coloured curves are analytical quantities (see Proposition 4.2 of Main Text) describing model-specific approximate performances of ParPGS if neutrophil count-specific causal variants and their effects are used instead (note $\overline{a} \approx 0.8$).

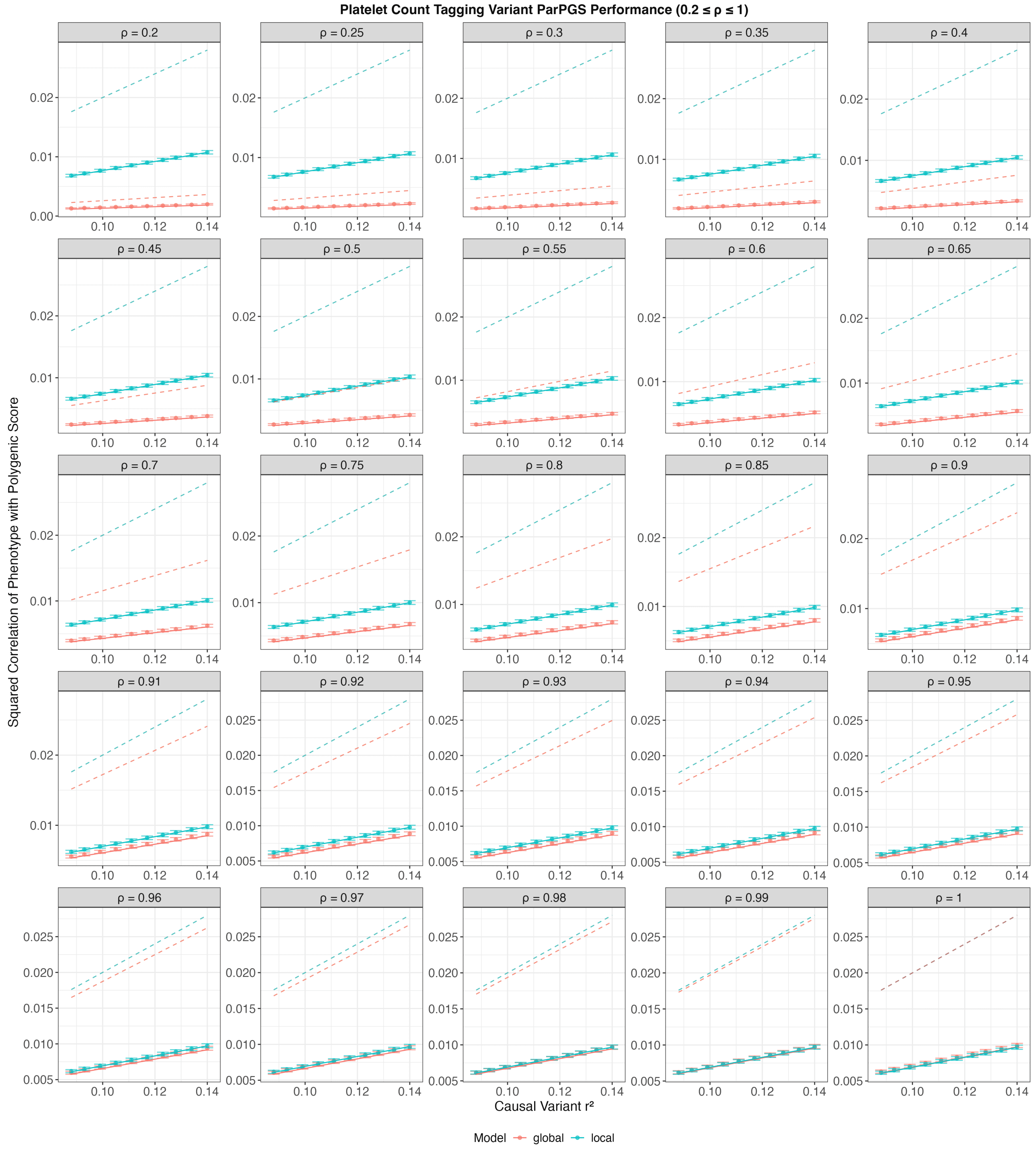

Figure S15: Partial PGS (ParPGS) performance when polygenic scores are computed using platelet count-specific tagging variants and simulated tagging effect sizes under Eq. (11) of Main Text. Solid coloured curves are approximate expectations of the performance under the local and the global model. Dashed coloured curves are analytical quantities (see Proposition 4.2 of Main Text) describing model-specific approximate performances of ParPGS if platelet count-specific causal variants and their effects are used instead (note $\overline{a} \approx 0.8$).

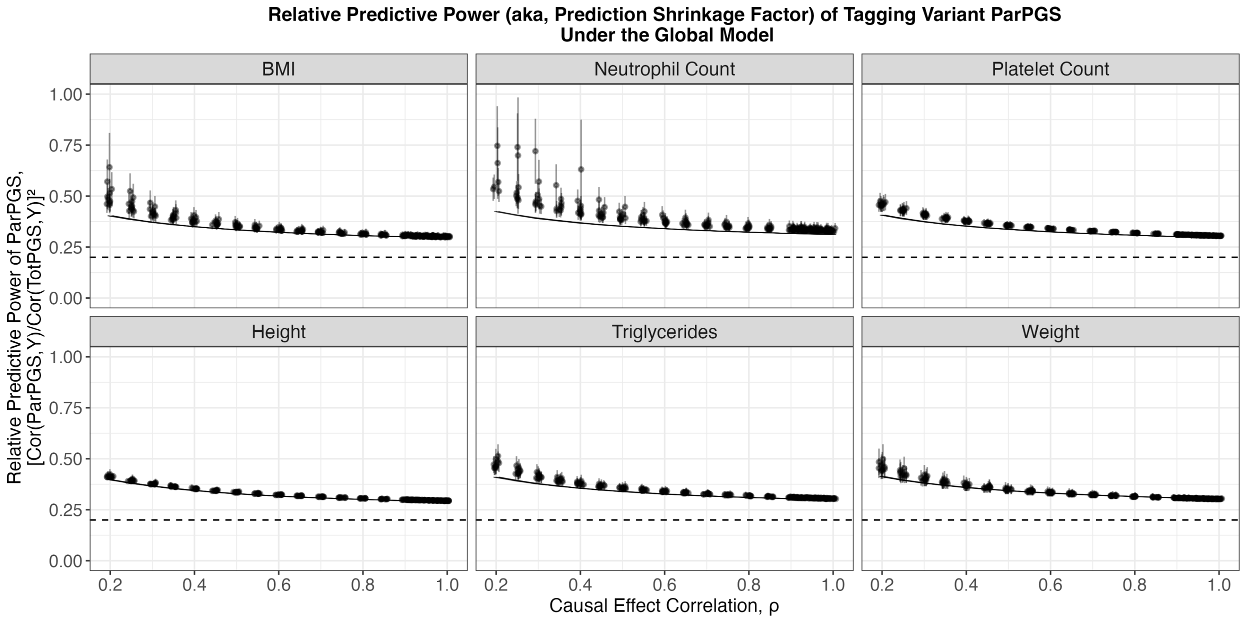

Figure S16: Relative predictive power — also referred to as the prediction shrinkage factor in Main Text — of partial polygenic scores (ParPGS) under the local model, in the general scenario whereby tagging variants are used to compute ParPGS and the standard polygenic score (TotPGS). Solid curves are grouped by $r^2$, and they depict approximate expectations of the ratio, obtained by computing the ratio of $\mathbb{E}_{\text{Glo}}[Cor(\mathbf{ParPGS}, \boldsymbol{y})^2]$ to $\mathbb{E}_{\text{Glo}}[Cor(\mathbf{TotPGS}, \boldsymbol{y})^2]$ using their approximations reported in Proposition S6. The dotted line shows $y = 0.2$, the prediction shrinkage factor predicted by the analytical approximation reported in Proposition 4.2 (note $\overline{a} \approx 0.8$).

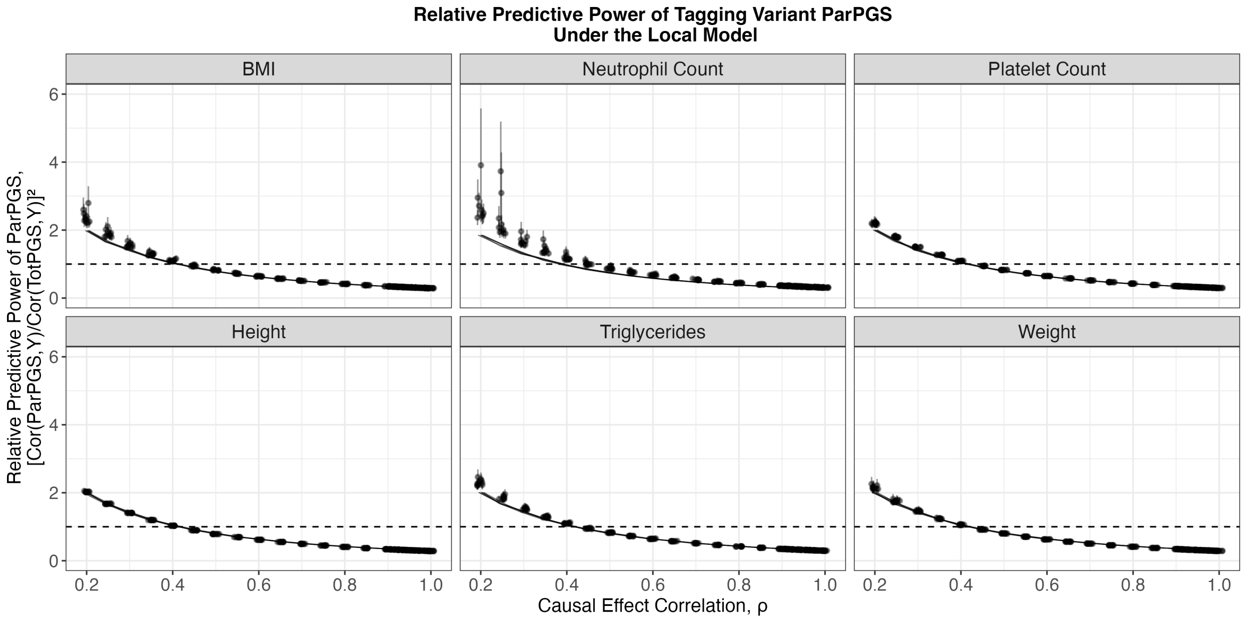

Figure S17: Relative predictive power of partial polygenic scores (ParPGS) under the local model, in the general scenario whereby tagging variants are used to compute ParPGS and the standard polygenic score (TotPGS). Solid curves are grouped by $r^2$, and they depict approximate expectations of the ratio, obtained by computing the ratio of $\mathbb{E}_{\text{Loc}}[Cor(\mathbf{ParPGS}, \boldsymbol{y})^2]$ to $\mathbb{E}_{\text{Loc}}[Cor(\mathbf{TotPGS}, \boldsymbol{y})^2]$ using their approximations reported in Proposition S6. The dotted line shows $y = 1$, such that a point lying above (resp., below) it indicates that ParPGS has greater (resp., lesser) predictive power than TotPGS.

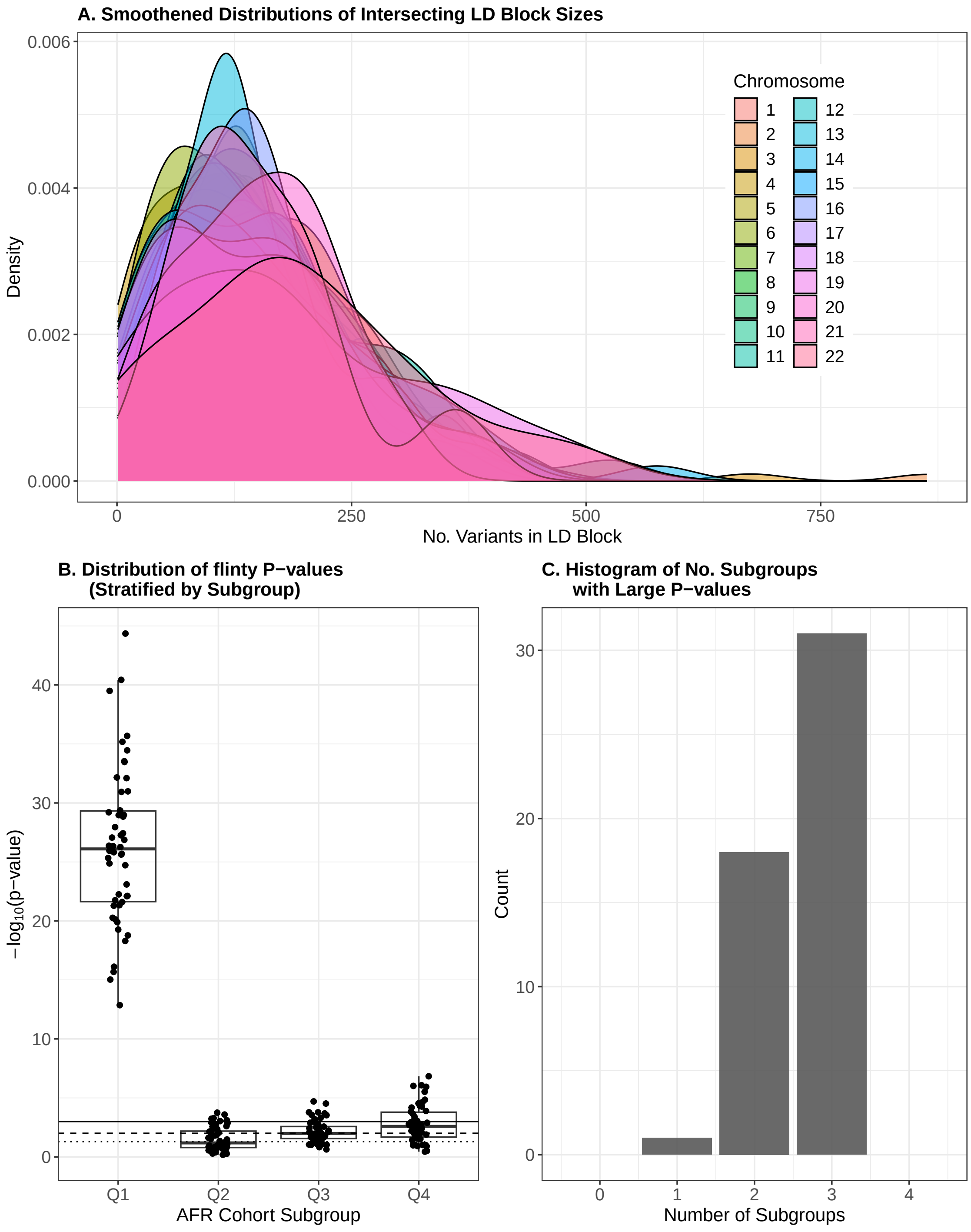

Figure S18: Selecting and Verifying Approximate Independence of Markers. **A.** Kernel density estimates of intersecting LD block sizes (i.e., number of variants in a block) obtained by intersecting LD blocks for homogeneous European and African populations. **B.** FLINTY $p$-values of feature independence and sample exchangeability, run on each subgroup of each seed. Note $p$-values are log-transformed for better visualization, and the horizontal lines depict significance thresholds $\alpha = 0.05$ (⋯⋯), $\alpha = 0.01$ (- - - -) and $\alpha = 0.001$ (——). **C.** Histogram of the number of subgroups, within a seed, for which $p$-value is greater than $\alpha = 2.5 \times 10^{-4}$. The minimum and maximum number are 0 and 4.

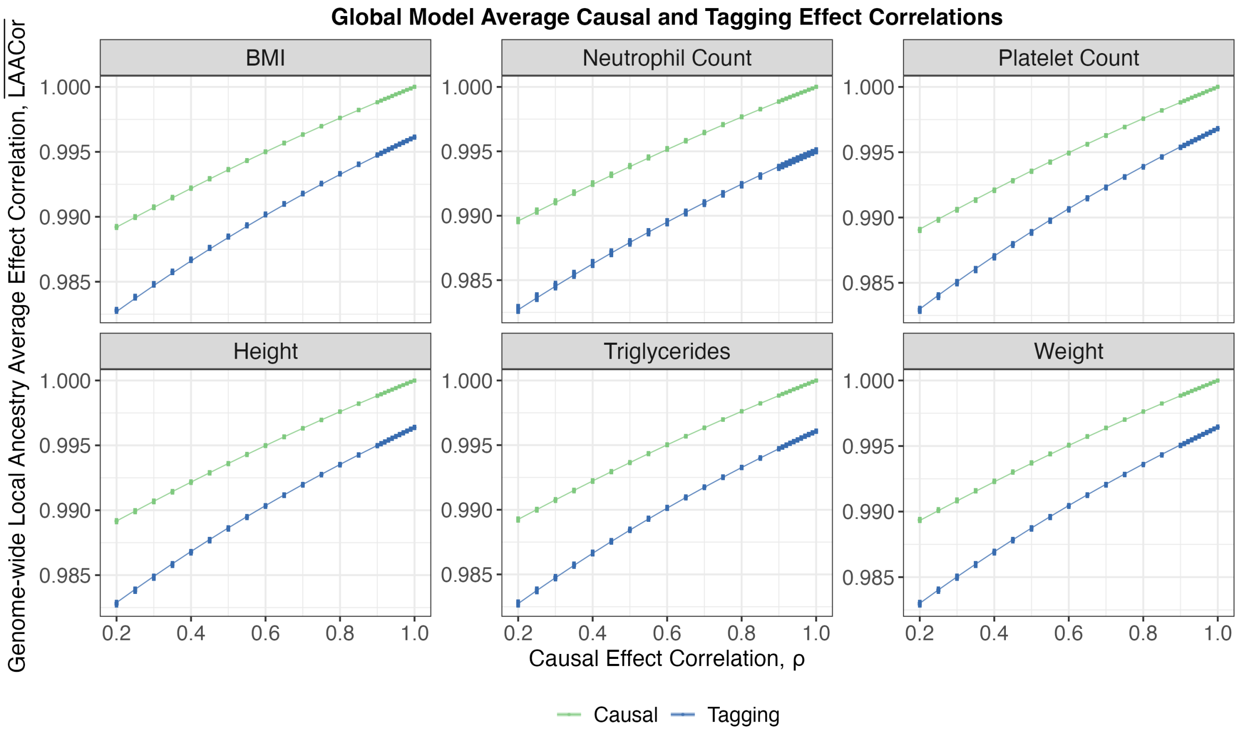

Figure S19: Local ancestry average causal and tagging effect correlations under the global model, across six phenotype-specific sets of putatively causal and tagging variants.

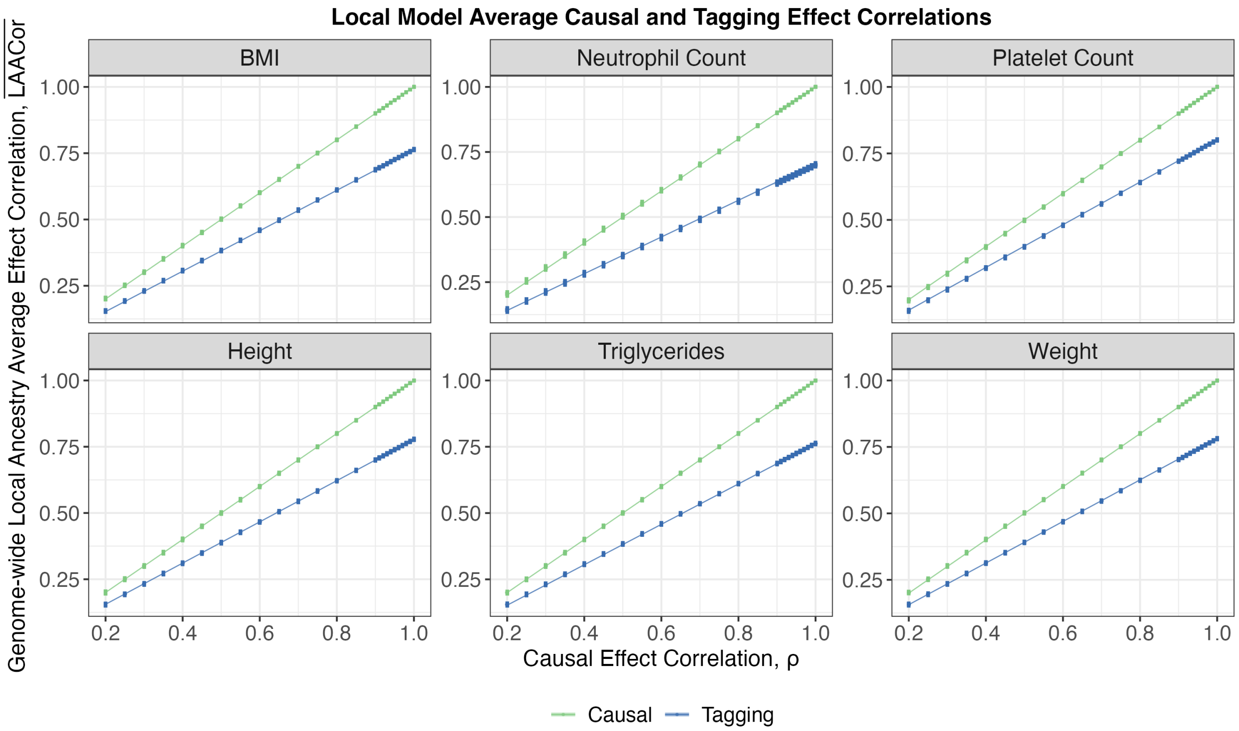

Figure S20: Local ancestry average causal and tagging effect correlations under the local model, across six phenotype-specific sets of putatively causal and tagging variants.

Table S1: Descriptions of various approaches investigating variant effect heterogeneity by ancestry.

| Reference | Finding About Variant Effect Comparisons | Loci and Populations Underlying Comparisons | Methodology | Traits Studied |
| --- | --- | --- | --- | --- |
| Chen et al. (2023) | "Trans-ethnic" genetic correlations are high | Loci in East Asian and in White British cohorts | Genetic correlation is inner product of least squares effect sizes (i.e., not estimated effects) | 21 quantitative (q.) traits shared between three Biobanks |
| Hou et al. (2023) | Effect sizes are highly correlated by local ancestry | Loci tagged by distinct ancestries within admixed African Americans | Ratio quantity measuring effect correlation by local ancestry; estimated via likelihood maximization | 24 PAGE q. traits<br>26 UKB q. traits<br>10 All of Us traits |
| Hu et al. (2025) | Effect sizes are highly similar between ancestries | Loci in White British and loci in African British | Ratio of regression slopes of polygenic scores to phenotype in admixed and European populations | 29 UKB q. traits |
| Patel et al. (2022) | Effect sizes differ by ancestry due to genetic interactions | Loci in European Americans and loci tagged with European ancestry in African Americans | Mixture coefficient that determines extent of divergence from European estimated effect in African-tagged locus | Gene expression traits in MESA<br><br>LDL Cholesterol in Million Veterans Project |
| Shi et al. (2021) | "Trans-ethnic" genetic correlations are lower in functionally important regions | Loci in East Asian and in White British cohorts | Similar to Chen et al. (2023), but only variants labeled with specific functional annotations included | 31 diseases and q. traits |

Table S2: Trait-specific causal and tagging variant summary statistics, along with the range of $r^2$ parameters used in simulations for that trait. $\lambda^2$ is computed as the squared correlation in the PMBB EUR Cohort, and as the square of the quantity $[\mathbb{P}(X = 1, X' = 1) - f \cdot f']/\sqrt{f(1-f)f'(1-f')}$ for pairs of variants tagged with African local ancestry in PMBB ADM Cohort, as described in Subsection 3.1 of Main Text.

| Phenotype | No. Variants ($p$) | Causal Variant Mean Afr Global Ancestry ($\bar{a}'$) | Tagging Variant Mean Afr Global Ancestry ($\bar{a}$) | Afr Mean $LD^2$ | Eur Mean $LD^2$ | Range of $r^2$ Used in Simulations |
|---|---|---|---|---|---|---|
| Standing Height | 2,476 | 0.795 | 0.795 | 0.435 | 0.854 | [0.176, 0.702] |
| Weight | 2,516 | 0.795 | 0.795 | 0.438 | 0.856 | [0.058, 0.111] |
| Body Mass Index | 2,958 | 0.795 | 0.795 | 0.435 | 0.856 | [0.041, 0.093] |
| Triglycerides | 2,331 | 0.795 | 0.795 | 0.437 | 0.856 | [0.047, 0.099] |
| Neutrophil Count | 738 | 0.796 | 0.796 | 0.427 | 0.852 | [0.023, 0.076] |
| Platelet Count | 2,322 | 0.795 | 0.795 | 0.439 | 0.857 | [0.088, 0.14] |

Table S3: Terms involved in calculating variances of the total and partial polygenic scores, which do not depend on the model choice (global vs local).

| **Quantity** | **Simplest Approximation** | **First Order Approximation** |
|---|---|---|
| $\mathbb{E}\left[\textcircled{1}\right]$ | $\mathbb{E}\left[\left(\beta_j^{\text{Eur}}\right)^2\right]\left[\left(1-\overline{a_{\cdot j}^{(1)}}\right)f_j^{\text{Eur}}\left(1-f_j^{\text{Eur}}\right)+\overline{a_{\cdot j}^{(1)}}f_j^{\text{Afr}}\left(1-f_j^{\text{Afr}}\right)\right]$ | $\frac{r^2(\lambda_j^{\text{Eur}})^2}{2p}\left[1-\overline{a_{\cdot j}^{(1)}}+\frac{f^{\text{Afr}}(1-f^{\text{Afr}})}{f^{\text{Eur}}(1-f^{\text{Eur}})}\overline{a_{\cdot j}^{(1)}}\right]$ |
| $\mathbb{E}\left[\textcircled{2}\right]$ | $\mathbb{E}\left[\left(\beta_j^{\text{Eur}}\right)^2\right]\left[\left(1-\overline{a_{\cdot j}^{(2)}}\right)f_j^{\text{Eur}}\left(1-f_j^{\text{Eur}}\right)+\overline{a_{\cdot j}^{(2)}}f_j^{\text{Afr}}\left(1-f_j^{\text{Afr}}\right)\right]$ | $\frac{r^2(\lambda_j^{\text{Eur}})^2}{2p}\left[1-\overline{a_{\cdot j}^{(2)}}+\frac{f^{\text{Afr}}(1-f^{\text{Afr}})}{f^{\text{Eur}}(1-f^{\text{Eur}})}\overline{a_{\cdot j}^{(2)}}\right]$ |
| $\mathbb{E}\left[\textcircled{3}\right]$ | $2\times\mathbb{E}\left[\left(\beta_j^{\text{Eur}}\right)^2\right]\frac{1}{n-1}\sum_{i=1}^n\hat{x}_{ij}^{(1)}\hat{x}_{ij}^{(2)}$ | $\frac{r^2(\lambda_j^{\text{Eur}})^2}{pf^{\text{Eur}}(1-f^{\text{Eur}})}\frac{1}{n-1}\sum_{i=1}^n\hat{x}_{ij}^{(1)}\hat{x}_{ij}^{(2)}$ |
| $\mathbb{E}\left[\textcircled{4}\right]$ | $\mathbb{E}\left[\left(\beta_j^{\text{Eur}}\right)^2\right]\left(1-\overline{a_{\cdot j}^{(1)}}\right)f_j^{\text{Eur}}\left(1-f_j^{\text{Eur}}\right)$ | $\frac{r^2(\lambda_j^{\text{Eur}})^2}{2p}\left(1-\overline{a_{\cdot j}^{(1)}}\right)$ |
| $\mathbb{E}\left[\textcircled{5}\right]$ | $\mathbb{E}\left[\left(\beta_j^{\text{Eur}}\right)^2\right]\left(1-\overline{a_{\cdot j}^{(2)}}\right)f_j^{\text{Eur}}\left(1-f_j^{\text{Eur}}\right)$ | $\frac{r^2(\lambda_j^{\text{Eur}})^2}{2p}\left(1-\overline{a_{\cdot j}^{(2)}}\right)$ |
| $\mathbb{E}\left[\textcircled{6}\right]$ | $2\times\mathbb{E}\left[\left(\beta_j^{\text{Eur}}\right)^2\right]\frac{1}{n-1}\sum_{i=1}^n\left(1-a_{ij}^{(1)}\right)\left(1-a_{ij}^{(2)}\right)\hat{x}_{ij}^{(1)}\hat{x}_{ij}^{(2)}$ | $\frac{r^2(\lambda_j^{\text{Eur}})^2}{pf^{\text{Eur}}(1-f^{\text{Eur}})}\frac{1}{n-1}\sum_{i=1}^n\left(1-a_{ij}^{(1)}\right)\left(1-a_{ij}^{(2)}\right)\hat{x}_{ij}^{(1)}\hat{x}_{ij}^{(2)}$ |

Table S4: Terms involved in calculating local model polygenic score performance.

| Quantity | Simplest Approximation |
|---|---|
| $\mathbb{E}\left[\textcircled{7L}\right]$ | $\frac{r^2\lambda_j^{\mathrm{Eur}}}{2p\sqrt{f^{\mathrm{Eur}}(1-f^{\mathrm{Eur}})f'^{\mathrm{Eur}}(1-f'^{\mathrm{Eur}})}}\frac{1}{n}\sum_{i=1}^{n}\left(1-a_{ij}'^{(1)}\right)\hat{x}_{ij}^{(1)}\hat{x}_{ij}'^{(1)}+\frac{r^2\rho\lambda_j^{\mathrm{Eur}}}{2p\sqrt{f^{\mathrm{Eur}}(1-f^{\mathrm{Eur}})f'^{\mathrm{Afr}}(1-f'^{\mathrm{Afr}})}}\frac{1}{n}\sum_{i=1}^{n}a_{ij}'^{(1)}\hat{x}_{ij}^{(1)}\hat{x}_{ij}'^{(1)}$ |
| $\mathbb{E}\left[\textcircled{8L}\right]$ | $\frac{r^2\lambda_j^{\mathrm{Eur}}}{2p\sqrt{f^{\mathrm{Eur}}(1-f^{\mathrm{Eur}})f'^{\mathrm{Eur}}(1-f'^{\mathrm{Eur}})}}\frac{1}{n}\sum_{i=1}^{n}\left(1-a_{ij}'^{(2)}\right)\hat{x}_{ij}^{(1)}\hat{x}_{ij}'^{(2)}+\frac{r^2\rho\lambda_j^{\mathrm{Eur}}}{2p\sqrt{f^{\mathrm{Eur}}(1-f^{\mathrm{Eur}})f'^{\mathrm{Afr}}(1-f'^{\mathrm{Afr}})}}\frac{1}{n}\sum_{i=1}^{n}a_{ij}'^{(2)}\hat{x}_{ij}^{(1)}\hat{x}_{ij}'^{(2)}$ |
| $\mathbb{E}\left[\textcircled{9L}\right]$ | $\frac{r^2\lambda_j^{\mathrm{Eur}}}{2p\sqrt{f^{\mathrm{Eur}}(1-f^{\mathrm{Eur}})f'^{\mathrm{Eur}}(1-f'^{\mathrm{Eur}})}}\frac{1}{n}\sum_{i=1}^{n}\left(1-a_{ij}'^{(1)}\right)\hat{x}_{ij}^{(2)}\hat{x}_{ij}'^{(1)}+\frac{r^2\rho\lambda_j^{\mathrm{Eur}}}{2p\sqrt{f^{\mathrm{Eur}}(1-f^{\mathrm{Eur}})f'^{\mathrm{Afr}}(1-f'^{\mathrm{Afr}})}}\frac{1}{n}\sum_{i=1}^{n}a_{ij}'^{(1)}\hat{x}_{ij}^{(2)}\hat{x}_{ij}'^{(1)}$ |
| $\mathbb{E}\left[\textcircled{10L}\right]$ | $\frac{r^2\lambda_j^{\mathrm{Eur}}}{2p\sqrt{f^{\mathrm{Eur}}(1-f^{\mathrm{Eur}})f'^{\mathrm{Eur}}(1-f'^{\mathrm{Eur}})}}\frac{1}{n}\sum_{i=1}^{n}\left(1-a_{ij}'^{(2)}\right)\hat{x}_{ij}^{(2)}\hat{x}_{ij}'^{(2)}+\frac{r^2\rho\lambda_j^{\mathrm{Eur}}}{2p\sqrt{f^{\mathrm{Eur}}(1-f^{\mathrm{Eur}})f'^{\mathrm{Afr}}(1-f'^{\mathrm{Afr}})}}\frac{1}{n}\sum_{i=1}^{n}a_{ij}'^{(2)}\hat{x}_{ij}^{(2)}\hat{x}_{ij}'^{(2)}$ |
| $\mathbb{E}\left[\textcircled{11L}\right]$ | $\frac{r^2(\lambda_j^{\mathrm{Eur}})^2}{2p}\left(1-\overline{a_{\cdot j}^{(1)}}-\overline{a_{\cdot j}'^{(1)}}+\overline{a_{\cdot j}^{(1)}a_{\cdot j}'^{(1)}}\right)+\frac{r^2\rho\lambda_j^{\mathrm{Eur}}}{2p\sqrt{f^{\mathrm{Eur}}(1-f^{\mathrm{Eur}})f'^{\mathrm{Afr}}(1-f'^{\mathrm{Afr}})}}\frac{1}{n}\sum_{i=1}^{n}\left(1-a_{ij}^{(1)}\right)a_{ij}'^{(1)}\hat{x}_{ij}^{(1)}\hat{x}_{ij}'^{(1)}$ |
| $\mathbb{E}\left[\textcircled{12L}\right]$ | $\frac{r^2\lambda_j^{\mathrm{Eur}}}{2p\sqrt{f^{\mathrm{Eur}}(1-f^{\mathrm{Eur}})f'^{\mathrm{Eur}}(1-f'^{\mathrm{Eur}})}}\frac{1}{n}\sum_{i=1}^{n}\left(1-a_{ij}^{(1)}\right)\left(1-a_{ij}'^{(2)}\right)\hat{x}_{ij}^{(1)}\hat{x}_{ij}'^{(2)}+\frac{r^2\rho\lambda_j^{\mathrm{Eur}}}{2p\sqrt{f^{\mathrm{Eur}}(1-f^{\mathrm{Eur}})f'^{\mathrm{Afr}}(1-f'^{\mathrm{Afr}})}}\frac{1}{n}\sum_{i=1}^{n}\left(1-a_{ij}^{(1)}\right)a_{ij}'^{(2)}\hat{x}_{ij}^{(1)}\hat{x}_{ij}'^{(2)}$ |
| $\mathbb{E}\left[\textcircled{13L}\right]$ | $\frac{r^2\lambda_j^{\mathrm{Eur}}}{2p\sqrt{f^{\mathrm{Eur}}(1-f^{\mathrm{Eur}})f'^{\mathrm{Eur}}(1-f'^{\mathrm{Eur}})}}\frac{1}{n}\sum_{i=1}^{n}\left(1-a_{ij}^{(2)}\right)\left(1-a_{ij}'^{(1)}\right)\hat{x}_{ij}^{(2)}\hat{x}_{ij}'^{(1)}+\frac{r^2\rho\lambda_j^{\mathrm{Eur}}}{2p\sqrt{f^{\mathrm{Eur}}(1-f^{\mathrm{Eur}})f'^{\mathrm{Afr}}(1-f'^{\mathrm{Afr}})}}\frac{1}{n}\sum_{i=1}^{n}\left(1-a_{ij}^{(2)}\right)a_{ij}'^{(1)}\hat{x}_{ij}^{(2)}\hat{x}_{ij}'^{(1)}$ |
| $\mathbb{E}\left[\textcircled{14L}\right]$ | $\frac{r^2(\lambda_j^{\mathrm{Eur}})^2}{2p}\left(1-\overline{a_{\cdot j}^{(2)}}-\overline{a_{\cdot j}'^{(2)}}+\overline{a_{\cdot j}^{(2)}a_{\cdot j}'^{(2)}}\right)+\frac{r^2\rho\lambda_j^{\mathrm{Eur}}}{2p\sqrt{f^{\mathrm{Eur}}(1-f^{\mathrm{Eur}})f'^{\mathrm{Afr}}(1-f'^{\mathrm{Afr}})}}\frac{1}{n}\sum_{i=1}^{n}\left(1-a_{ij}^{(2)}\right)a_{ij}'^{(2)}\hat{x}_{ij}^{(2)}\hat{x}_{ij}'^{(2)}$ |

Table S5: Terms involved in calculating global model polygenic score performance.

| Quantity | Simplest Approximation |
| --- | --- |
| $\mathbb{E}\left[\textcircled{7G}\right]$ | $\frac{r^2\lambda_j^{\text{Eur}}}{2p\sqrt{f^{\text{Eur}}(1-f^{\text{Eur}})f'^{\text{Eur}}(1-f'^{\text{Eur}})}}\frac{1}{n}\sum_{i=1}^{n}\left(1-a'^{(1)}_{ij}\right)\hat{x}^{(1)}_{ij}\hat{x}'^{(1)}_{ij}+\frac{r^2\rho\lambda_j^{\text{Eur}}}{2p\sqrt{f^{\text{Eur}}(1-f^{\text{Eur}})f'^{\text{Afr}}(1-f'^{\text{Afr}})}}\frac{1}{n}\sum_{i=1}^{n}a'^{(1)}_{ij}\hat{x}^{(1)}_{ij}\hat{x}'^{(1)}_{ij}$ |
| $\mathbb{E}\left[\textcircled{8G}\right]$ | $\frac{r^2\lambda_j^{\text{Eur}}}{2p\sqrt{f^{\text{Eur}}(1-f^{\text{Eur}})f'^{\text{Eur}}(1-f'^{\text{Eur}})}}\frac{1}{n}\sum_{i=1}^{n}\left(1-a'^{(2)}_{ij}\right)\hat{x}^{(1)}_{ij}\hat{x}'^{(2)}_{ij}+\frac{r^2\rho\lambda_j^{\text{Eur}}}{2p\sqrt{f^{\text{Eur}}(1-f^{\text{Eur}})f'^{\text{Afr}}(1-f'^{\text{Afr}})}}\frac{1}{n}\sum_{i=1}^{n}a'^{(2)}_{ij}\hat{x}^{(1)}_{ij}\hat{x}'^{(2)}_{ij}$ |
| $\mathbb{E}\left[\textcircled{9G}\right]$ | $\frac{r^2\lambda_j^{\text{Eur}}}{2p\sqrt{f^{\text{Eur}}(1-f^{\text{Eur}})f'^{\text{Eur}}(1-f'^{\text{Eur}})}}\frac{1}{n}\sum_{i=1}^{n}\left(1-a'^{(1)}_{ij}\right)\hat{x}^{(2)}_{ij}\hat{x}'^{(1)}_{ij}+\frac{r^2\rho\lambda_j^{\text{Eur}}}{2p\sqrt{f^{\text{Eur}}(1-f^{\text{Eur}})f'^{\text{Afr}}(1-f'^{\text{Afr}})}}\frac{1}{n}\sum_{i=1}^{n}a'^{(1)}_{ij}\hat{x}^{(2)}_{ij}\hat{x}'^{(1)}_{ij}$ |
| $\mathbb{E}\left[\textcircled{10G}\right]$ | $\frac{r^2\lambda_j^{\text{Eur}}}{2p\sqrt{f^{\text{Eur}}(1-f^{\text{Eur}})f'^{\text{Eur}}(1-f'^{\text{Eur}})}}\frac{1}{n}\sum_{i=1}^{n}\left(1-a'^{(2)}_{ij}\right)\hat{x}^{(2)}_{ij}\hat{x}'^{(2)}_{ij}+\frac{r^2\rho\lambda_j^{\text{Eur}}}{2p\sqrt{f^{\text{Eur}}(1-f^{\text{Eur}})f'^{\text{Afr}}(1-f'^{\text{Afr}})}}\frac{1}{n}\sum_{i=1}^{n}a'^{(2)}_{ij}\hat{x}^{(2)}_{ij}\hat{x}'^{(2)}_{ij}$ |
| $\mathbb{E}\left[\textcircled{11G}\right]$ | $\frac{r^2(\lambda_j^{\text{Eur}})^2}{2p}\left(1-\overline{a^{(1)}_{\cdot j}}-\overline{a'^{(1)}_{\cdot j}}+\overline{a^{(1)}_{\cdot j}a'^{(1)}_{\cdot j}}\right)+\frac{r^2\rho\lambda_j^{\text{Eur}}}{2p\sqrt{f^{\text{Eur}}(1-f^{\text{Eur}})f'^{\text{Afr}}(1-f'^{\text{Afr}})}}\frac{1}{n}\sum_{i=1}^{n}\left(1-a^{(1)}_{ij}\right)a'^{(1)}_{ij}\hat{x}^{(1)}_{ij}\hat{x}'^{(1)}_{ij}$ |
| $\mathbb{E}\left[\textcircled{12G}\right]$ | $\frac{r^2\lambda_j^{\text{Eur}}}{2p\sqrt{f^{\text{Eur}}(1-f^{\text{Eur}})f'^{\text{Eur}}(1-f'^{\text{Eur}})}}\frac{1}{n}\sum_{i=1}^{n}\left(1-a^{(1)}_{ij}\right)\left(1-a'^{(2)}_{ij}\right)\hat{x}^{(1)}_{ij}\hat{x}'^{(2)}_{ij}+\frac{r^2\rho\lambda_j^{\text{Eur}}}{2p\sqrt{f^{\text{Eur}}(1-f^{\text{Eur}})f'^{\text{Afr}}(1-f'^{\text{Afr}})}}\frac{1}{n}\sum_{i=1}^{n}\left(1-a^{(1)}_{ij}\right)a'^{(2)}_{ij}\hat{x}^{(1)}_{ij}\hat{x}'^{(2)}_{ij}$ |
| $\mathbb{E}\left[\textcircled{13G}\right]$ | $\frac{r^2\lambda_j^{\text{Eur}}}{2p\sqrt{f^{\text{Eur}}(1-f^{\text{Eur}})f'^{\text{Eur}}(1-f'^{\text{Eur}})}}\frac{1}{n}\sum_{i=1}^{n}\left(1-a^{(2)}_{ij}\right)\left(1-a'^{(1)}_{ij}\right)\hat{x}^{(2)}_{ij}\hat{x}'^{(1)}_{ij}+\frac{r^2\rho\lambda_j^{\text{Eur}}}{2p\sqrt{f^{\text{Eur}}(1-f^{\text{Eur}})f'^{\text{Afr}}(1-f'^{\text{Afr}})}}\frac{1}{n}\sum_{i=1}^{n}\left(1-a^{(2)}_{ij}\right)a'^{(1)}_{ij}\hat{x}^{(2)}_{ij}\hat{x}'^{(1)}_{ij}$ |
| $\mathbb{E}\left[\textcircled{14G}\right]$ | $\frac{r^2(\lambda_j^{\text{Eur}})^2}{2p}\left(1-\overline{a^{(2)}_{\cdot j}}-\overline{a'^{(2)}_{\cdot j}}+\overline{a^{(2)}_{\cdot j}a'^{(2)}_{\cdot j}}\right)+\frac{r^2\rho\lambda_j^{\text{Eur}}}{2p\sqrt{f^{\text{Eur}}(1-f^{\text{Eur}})f'^{\text{Afr}}(1-f'^{\text{Afr}})}}\frac{1}{n}\sum_{i=1}^{n}\left(1-a^{(2)}_{ij}\right)a'^{(2)}_{ij}\hat{x}^{(2)}_{ij}\hat{x}'^{(2)}_{ij}$ |
